# Robust and efficient online auditory psychophysics

**DOI:** 10.1101/2021.07.17.452796

**Authors:** Sijia Zhao, Christopher A. Brown, Lori L. Holt, Frederic Dick

**Affiliations:** Department of Experimental Psychology, University of Oxford, Oxford OX1 3PH, UK; Department of Communication Science and Disorders, University of Pittsburgh, 4028 Forbes Tower Pittsburgh, PA 15260; Department of Psychology, Carnegie Mellon University, 5000 Forbes Avenue, Pittsburgh, PA 15232; Neuroscience Institute, Carnegie Mellon University, 5000 Forbes Avenue, Pittsburgh, PA 15232; Department of Psychological Sciences, Birkbeck College, University of London, Malet Street, London WC1E 7HX, UK; Department of Experimental Psychology, PALS, University College London, 26 Bedford Way, London, WC1H 0AP, UK

**Keywords:** Online testing, Probe-signal, Psychophysics, Auditory Thresholds, Motivation

## Abstract

Most human auditory psychophysics research has historically been conducted in carefully controlled environments with calibrated audio equipment, and over potentially hours of repetitive testing with expert listeners. Here, we operationally define such conditions as having high ‘auditory hygiene’. From this perspective, conducting auditory psychophysical paradigms online presents a serious challenge, in that results may hinge on absolute sound presentation level, reliably estimated perceptual thresholds, low and controlled background noise levels, and sustained motivation and attention. We introduce a set of procedures that address these challenges and facilitate auditory hygiene for online auditory psychophysics. First, we establish a simple means of setting sound presentation levels. Across a set of four level-setting conditions conducted in person, we demonstrate the stability and robustness of this level setting procedure in open air and controlled settings. Second, we test participants’ tone-in-noise thresholds using widely adopted online experiment platforms and demonstrate that reliable threshold estimates can be derived online in approximately one minute of testing. Third, using these level and threshold setting procedures to establish participant-specific stimulus conditions, we show that an online implementation of the classic probe-signal paradigm can be used to demonstrate frequency-selective attention on an individual-participant basis, using a third of the trials used in recent in-lab experiments. Finally, we show how threshold and attentional measures relate to well-validated assays of online participants’ in-task motivation, fatigue, and confidence. This demonstrates the promise of online auditory psychophysics for addressing new auditory perception and neuroscience questions quickly, efficiently, and with more diverse samples. Code for the tests is publicly available through Pavlovia and Gorilla.

## Introduction

Much of what we know about the function of the auditory system is due to a century of auditory psychophysical behavioral paradigms in human listeners. Auditory psychophysics tends to rely on strongly sound-attenuated environments, finely calibrated equipment, and small numbers of expert or highly trained listeners who are motivated and compliant with task demands. This high level of what we term ‘auditory hygiene’ is important: seemingly minute differences in stimulus delivery and timing, background noise levels, or participant engagement during attention-demanding paradigms for measuring perceptual thresholds can dramatically affect experimental results (Green, 1995; Manning, Jones, Dekker, & Pellicano, 2018; Rinderknecht, Ranzani, Popp, Lambercy, & Gassert, 2018).

The COVID pandemic taught us the utility of online testing and challenged how we maintain auditory hygiene when lab facilities are inaccessible; the need to include more diverse and representative participant samples has also driven a move toward more inclusive experimental environments (Henrich, Heine, & Norenzayan, 2010; Rad, Martingano, & Ginges, 2018) particularly using online experimentation services (Anwyl-Irvine, Massonnié, Flitton, Kirkham, & Evershed, 2020; Buhrmester, Kwang, & Gosling, 2011; Peirce et al., 2019, p. 2; Sauter, Draschkow, & Mack, 2020). As highlighted in a recent report by the ASA Task Force on Remote Testing (https://tcppasa.org/remote-testing/) human auditory researchers have created a number of methods to maintain high standards using out-of-laboratory testing. For instance, several groups have created tests for ensuring participants are using headphones rather than speakers (Milne et al., 2020; Woods, Siegel, Traer, & McDermott, 2017), and that they are engaging with the experimental task, rather than haphazardly pressing buttons (Bianco, Mills, de Kerangal, Rosen, & Chait, 2021; Mok et al., 2021; Zhao et al., 2019). Such innovations notwithstanding, uncontrolled online experimental situations are particularly challenging for auditory paradigms that deliver stimuli within a range of sound pressure levels, or that require sustained vigilance to respond consistently to an ever more difficult-to-perceive target sound.

Control of the range of sound pressure levels is important for ensuring participants’ well-being, making sure they are not exposing themselves to overly loud sounds. Sound pressure level is also important because neuronal responses from the cochlea to cortex are known to differ as a function of overall level. For instance, subpopulations of auditory nerve fibers differing in spontaneous firing rates respond at different acoustic stimulation levels (Horst, McGee, & Walsh, 2018; Taberner & Liberman, 2005). Across the peripheral and central auditory systems, single neuronal responses tend to be level-dependent, with frequency selectivity typically broadening with increasing sound amplitude levels (Bizley, Nodal, Nelken, & King, 2005; Schreiner, Read, & Sutter, 2000). Behaviorally derived auditory filter widths have also been shown to be level-dependent (Glasberg & Moore, 2000; Pick, 1980). This is particularly important for experiments that aim to compare perceptual versus attentional auditory filters, such as in the classic ‘probe-signal’ paradigm presented below (Anandan, Husain, & Seluakumaran, 2021; Borra, Versnel, Kemner, van Opstal, & van Ee, 2013; Botte, 1995; Dai & Buus, 1991; Dai, Scharf, & Buus, 1991; Dai et al., 1991; Green & McKeown, 2001; Greenberg & Larkin, 1968, 1968; Macmillan & Schwartz, 1975; Moore, Hafter, & Glasberg, 1996; Scharf, Quigley, Aoki, Peachey, & Reeves, 1987; Scharf et al., 1987; Tan, Robertson, & Hammond, 2008).

Many auditory experiments, including the probe-signal paradigm, typically ask listeners to perceive stimuli at or near their perceptual thresholds for hearing out a stimulus in quiet or in a masking noise or background. These thresholds can differ considerably across individuals, so often experimental sessions will begin by running adaptive psychophysical paradigms to estimate the individual’s relevant perceptual thresholds. Obtaining reliable auditory psychophysical thresholds can be challenging, even in laboratory conditions with experienced and motivated adult listeners. For example, thresholds-in-quiet have been shown to be affected by the duration of time spent in a ‘quiet’ environment (Bryan, Parbrook, & Tempest, 1965; Steed & Martin, 1973) such as an audiometric booth. Even supra-threshold detection tasks performed by experienced listeners can be affected by presentation level (Williams, Elfner, & Howse, 1978). Determining reliable psychoacoustical thresholds may be especially hard with inexperienced listeners (Kopiez & Platz, 2009) or in the presence of distracting events (Ruggles, Bharadwaj, & Shinn-Cunningham, 2011) typical of a home environment.

Especially for online studies where participants are in their home environments, reduced levels of engagement and vigilance due to listeners’ motivation, fatigue, and confidence can inject additional noise and bias (general discussion in Elfadaly et al., 2020). This is particularly true when paradigms required to set perceptual levels for the actual experiments of interest are themselves potentially tedious and unrewarding (reviewed in Jones, 2019). Multiple long thresholding tracks also add considerable expense to online experiments, which tend to rely on shorter experimental sessions with larger numbers of participants to compensate for participant variability. A number of investigators have optimized psychophysics techniques for measuring perceptual thresholds in different populations. For instance, Dillon et al. (Dillon, Beach, Seymour, Carter, & Golding, 2016) used Monte Carlo simulations to create an efficient adaptive algorithm for telephone-based speech-in-noise threshold measurement. Others have designed ‘participant-friendly’ procedures for pediatric psychoacoustics testing (for example, Halliday, Tuomainen, & Rosen, 2017) that manipulate different stepping rules, for instance changing reversal rules once a first error has been made (Baker & Rosen, 2001).

Nonetheless, lapses in attentive listening in repetitive and challenging tasks like the staircase threshold setting procedures described above can dramatically impact experimental results. Thus, concern that anonymous, online participants may be less motivated to perform to the best of their abilities, as compared to more traditional in-person expert listeners has contributed to reticence in moving auditory investigation online.

In a set of three experiments, we address the challenges of sound level setting, psychophysical threshold estimation, and participant motivation, engagement and vigilance in online auditory psychophysics experiments. To this end, we test new online versions of level setting and threshold-in-noise paradigms, as well as a short-duration online version of the aforementioned probe-signal paradigm. We also evaluate whether results are potentially modulated by participants’ motivation and fatigue levels.

In Experiment 1, we assess a method for controlling the range of experimental stimulus levels (within ± 10 dBA SPL) in online testing conducted in uncontrolled environments. To do this, we have participants act as a ‘self-calibrated audiometer’ by listening to a white or pink noise stimulus with a particular root-mean-square amplitude (RMS), then adjusting the volume setting on their own computer to a just-detectable threshold^1^. To assess the validity of this approach, participants take part in the online amplitude setting task in uncontrolled environments and in the laboratory.

In Experiment 2, we incorporate the level-setting paradigm introduced in Experiment 1, then ask whether small adjustments to standard thresholding procedures for a classic psychophysical task (tone detection in white noise) will permit fast (2-3 minute) and reliable estimation of thresholds among participants recruited and tested online. Specifically, we evaluate three factors. One, we test the reliability of estimates over three short (40-trial) staircase-based thresholding tracks. Two, we examine whether a simple estimator of psychophysical threshold - the statistical mode of levels across a thresholding track (e.g., the most frequently visited level) - is as robust or more robust at estimating threshold as traditional estimators based on staircase reversals. Three, we determine whether and how online psychophysical thresholds are related to established assays of participant fatigue, apathy, and task confidence.

In Experiment 3, we use the online tone-in-noise thresholding procedure from Experiment 2 to set participants thresholds for a new online version of the probe-signal paradigm (Botte, 1995; Dai & Buus, 1991; Dai et al., 1991; Greenberg & Larkin, 1968; Moore et al., 1996; Scharf et al., 1987). After completing the online threshold-setting procedure of Experiment 2, the same online participants heard continuous noise in which an above-threshold tone was followed by two listening intervals. Participants reported the interval in which a near-threshold tone was embedded in the noise, with the tone frequency matching the cue on 75% of trials and mismatching the cue at one of four other frequencies on 25% of the trials. We sought to determine whether patterns of frequency-selective attention: 1) can be replicated in uncontrolled online testing environments with naive listeners; 2) are evident in the short testing sessions necessitated by online testing; 3) change and develop over testing trials; and 4) are related to established assays of participant fatigue, apathy, and task confidence.

We provide code for each of these approaches to facilitate improved ‘auditory hygiene’ in online experiments, and to demonstrate the possibilities for asking new questions in auditory science with classic, yet challenging, online psychophysical paradigms. Our goal is to test and validate procedures for good ‘auditory hygiene’ in less controlled environments so that online studies can be as rigorous as (and directly compared to) in-lab studies.

## Experiment 1

In the four conditions of Experiment (Expt) 1a-d (see Table 1), we ask whether we can control the range of experimental stimulus levels in online testing conducted in different environments. Our approach involves playing a reference white or pink noise segment and having young adult online participants with healthy hearing adjust the volume setting on the computer to just-detectable levels. Rather like the “biological check” employed daily to confirm (though not adjust) level calibration in most audiology clinics, this procedure allows for each participant to use their normal hearing thresholds to adjust for their unique testing equipment and acoustic environment. The RMS amplitude of the white noise stimulus used for setting this detection threshold is then used as a reference value for setting the amplitude of subsequent experimental stimuli during the same session.

**Table 1:**
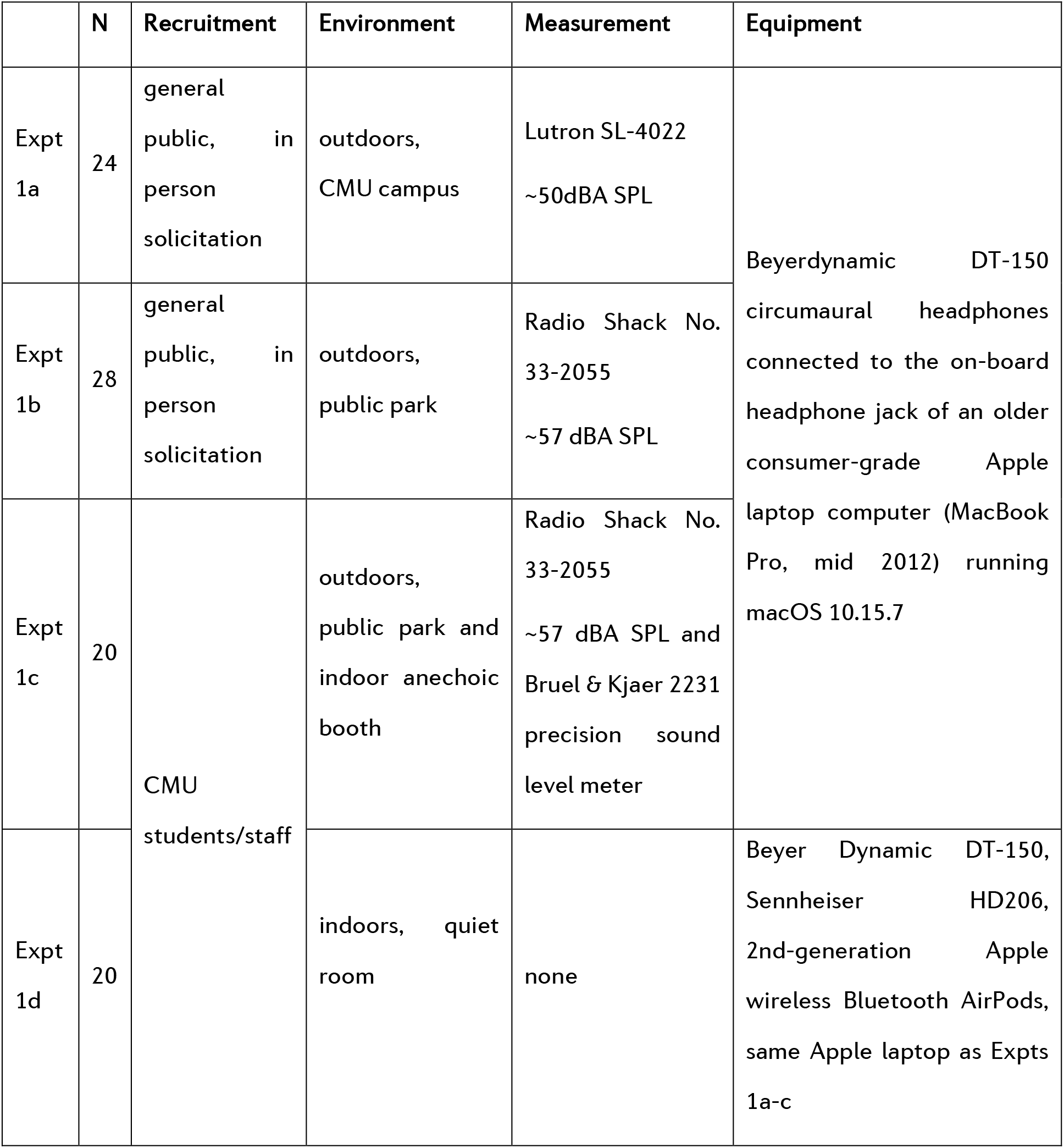
Overview of Experiment 1. Details differentiating Experiments 1a-d are shown.

In Expt conditions 1a and 1b, we tested different members of the general public outdoors using a pulsed bandpass-filtered white noise; given the level of distraction and background sound, these experiments provide initial real-world tests of the level setting paradigm. In condition 1c, we tested a group of Carnegie Mellon University affiliates to assess the reliability of the level setting paradigm over different listening conditions by having the same participants complete the task outdoors and in an anechoic chamber. Finally, in condition 1d, we tested another group of Carnegie Mellon University affiliates with bandpass-filtered white and pink noise to ask how level setting might be affected by spectral shape; to assess consistency across headphones, the same participants were also tested with white noise only using two different headphones as well as a popular brand of earbuds.

## Methods

### Participants

Validation of the online level setting procedure required testing in-person participants on a common consumer laptop with consumer headphones (with headphone type manipulated across conditions). For Expts 1a and 1b, recruitment was primarily conducted via informal in-person solicitation in outdoor environments due to COVID-related restrictions on indoor activities that were in place during data collection, and because the total task duration was approximately 2 minutes. For Expt 1a, participants (N=24) were recruited in an open lawn on the Carnegie Mellon University campus; a subset of participants were graduate students at a departmental gathering, others were undergraduate students as well as parents visiting for graduation ceremonies. Expt 1a participants were asked only whether they were at least 18 years of age, and considered their hearing to be within normal ranges, similar to the information that is solicited in many online studies. For Expt 1b, participants (N=28) were recruited in a central Pittsburgh park from a more heterogenous pool; here, participants were asked to note their age (mean age = 27.9 years (SD 10.2), ranging between 18 and 55 years). One of these participants mentioned that they occasionally wore hearing aids. For Expt 1c, all participants were Carnegie Mellon or University of Pittsburgh students or staff (N=20; mean age = 30.1 years (SD 9.2), age range 17-47 years); here the same individuals were tested in the outdoor environment as well as in an anechoic sound booth under well-controlled laboratory conditions. For Expt 1d, all participants were Carnegie Mellon students or staff (N=20, mean age = 25.4 years (SD 5.2, age range 18-37 years; these were not the same participants as Expt 1c).

The study was approved by the Birkbeck College ethics committee (181941/200518) for online testing without geographic restrictions, and took approximately 2 minutes to complete, including reading and completing the consent form, reading instructions, and performing the amplitude-setting task. Face-to-face participants were covered by local Carnegie Mellon University or University of Pittsburgh IRB protocols, as appropriate.

### Stimuli and Equipment

Using Praat 6.0.17 (Boersma & Weenink, 2021) a 1-second Gaussian white noise was generated, and band-pass filtered between 80-8000 Hz to restrict high-frequency contributions to overall intensity and low-frequency line noise. The RMS amplitude within Praat was adjusted to 0.000399 (26 dB). This amplitude setting was chosen as pilot testing suggested it allowed thresholds-in-quiet to be achieved within the range of laptop volume control settings (see Stimulus Analysis section below for analysis of analog stimulus output from two laptops).

Raised-cosine onset and offset ramps of 100-ms were added, so that when played on a continuous loop without gaps, it would sound like a sequence of pulsed noises. The audio data were stored in the WAV file format, then exported in Sox (http://sox.sourceforge.net/) to a stereo (diotic) sound file in the lossless FLAC format. This RMS level of this stimulus file serves as the amplitude reference for the sound stimuli in Expts 2 and 3. For Expt 1d only, a pink noise stimulus (with 1/f power spectral density) with the same duration, onset/offset ramps, and RMS as the white noise was also saved to FLAC format.

For Expts 1a-c, stimuli were presented using Beyerdynamic DT-150 circumaural headphones connected to the on-board headphone jack of an older consumer-grade Apple laptop computer (MacBook Pro, mid 2012) running macOS 10.15.7. For Expt 1d, which tested the procedure with different grade headphones, the Beyerdynamic DT-150 (∼$200 US), along with Sennheiser HD206 (∼$20 US), and 2nd-generation Apple wireless Bluetooth AirPods (∼$150 US) were used with the same Apple laptop.

For experiments 1a-c, outdoor sound levels were measured using a Lutron SL-4022 (Expt 1a) or a Radio Shack Cat No. 33-2055 sound level meter (Expt 1b and 1c). For Expt 1a, Baseline average sound levels were ∼50 dB SPL A-weighted; for Expt 1b, they were somewhat higher, with an average of ∼57dBA SPL, ranging between ∼53-67 dBA SPL. For Expt 1c, sound levels were an average of 31 dBA SPL indoors in the anechoic chamber (using a Bruel & Kjaer 2231 precision sound level meter) and 57 dBA SPL outdoors. As with many real-world listening environments, the outdoor environments included frequent sound events of somewhat higher amplitude (bird chirps, conversations of passing people, motorized skateboards, and helicopters flying overhead). See Supplemental Figure S1 for power spectral densities of the acoustic environments used in Expt 1c. (Expt 1d was conducted indoors in quiet rooms so we did not measure ambient sound levels).

### Calibration

For each volume setting increment on the MacBook Pro, dB SPL measurements were obtained using a Bruel & Kjaer 2231 precision sound level meter set to slow averaging and A-scale weighting and Bruel & Kjaer 4155 ½” microphone mounted in a Bruel & Kjaer 4152 artificial ear with a flat-plate coupler, coupled to the same set of Beyer DT-150 headphones used for data collection. Stimuli were played with exactly the same procedure and Macbook Pro as used for participant testing. This calibration routine was conducted in an anechoic chamber located on the University of Pittsburgh campus with an ambient noise floor measured to be about 31 dBA SPL using the same Bruel & Kjaer meter and coupler set up, as detailed above, but with the headphones disconnected. Note that the coupler simulates ear canal resonance, which when paired with A-Scale weighting, magnified the associated band-pass filtering and thus likely underestimated SPL at the eardrum.

Because the SPL of the stimulus at the lowest volume settings was below this noise floor, the white noise stimulus was digitally increased in level by 10 and 20 dB, and SPL values were then recorded at all volume settings for these two more intense stimuli, as well as for the original stimulus used during testing. The SPL-Volume setting functions generated using the more intense stimuli were then used to extrapolate the same function from the original stimulus below the noise floor (See Figure 1). Volume setting adjustments were determined to be linear on the MacBook Pro used in the amplitude setting experiment, e.g., a given increment in volume setting generated a relatively consistent change in dBA SPL at both high and low overall levels. This result gave us confidence that we could extrapolate downward to and below the noise floor. For a fuller picture of measuring below the noise floor, please see Ellingson, Gallun, & Bock (2015) and Whittle & Evans (1972).

**Figure 1.**
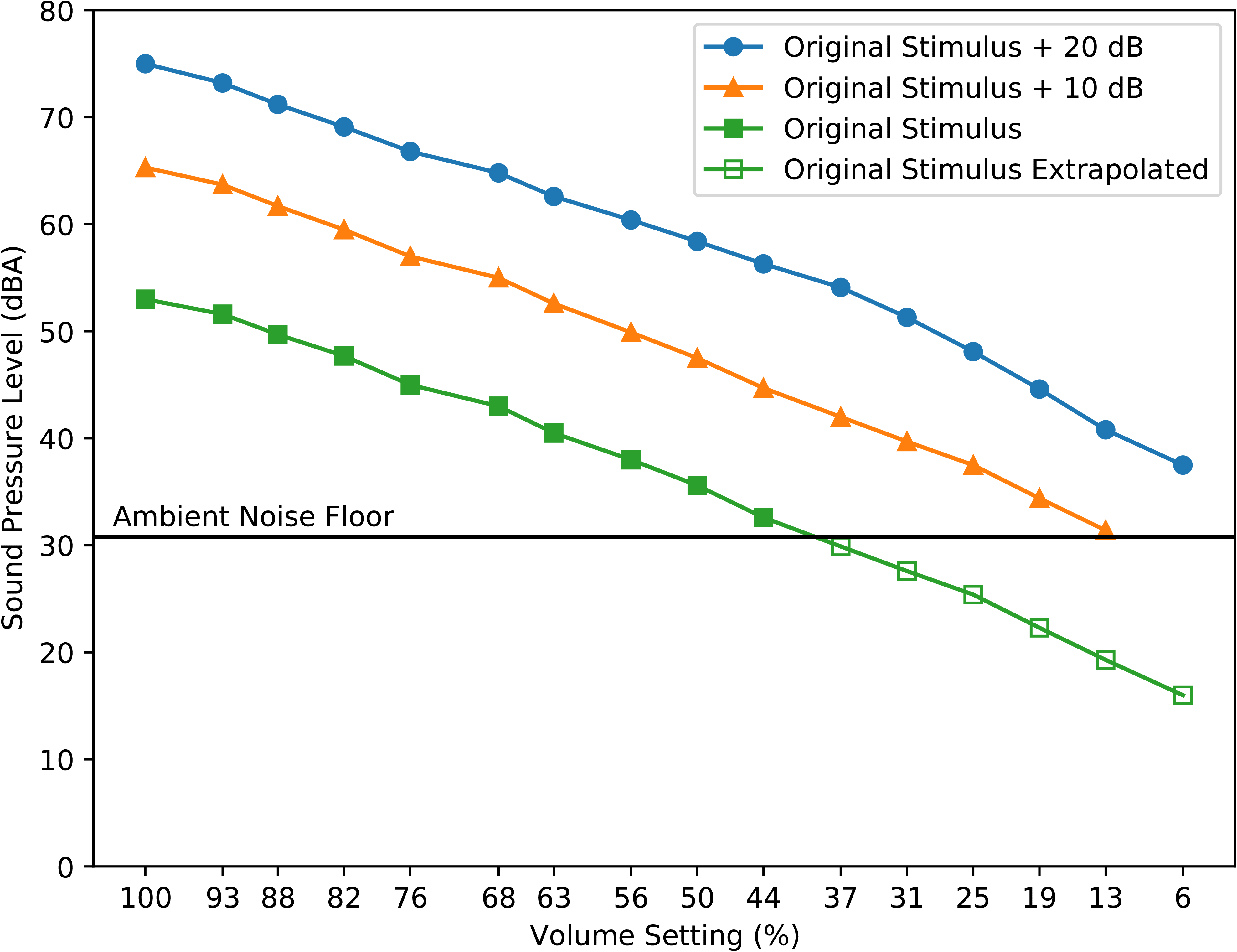
Sound pressure levels of the noise stimulus as a function of computer volume setting percentages. The noise stimulus was the same bandpass-filtered white noise used for testing or was increased in intensity by 10 or 20 dB. Measurements were made by playing each stimulus at each volume setting of the Macbook Pro using the headphones used in Experiments 1a-1c, coupled to an artificial ear. Because the SPL of the stimulus at the RMSv used for testing was below the ambient noise floor at lower volume setting values, the volume-setting functions at +10 and +20 dB were used to extrapolate the test stimulus function. SPL is in dBA.

The results of this acoustic analysis indicated that the highest volume setting (100%) produced a stimulus presentation level of 55 dBA SPL, and the lowest (6%) corresponded to 19.3 dBA SPL. Figure 1 shows dBA SPL values for the band-passed white noise stimulus at various levels (original level used during testing, and +10 and +20 dB) at each volume setting.

### Recording and analysis of laptop stimulus output to headphones

In order to deliver sound levels near detection threshold via standard laptops and headphones, the RMS of the white noise audio file needed to be very low (0.000399), raising the possibility that the signal would be distorted due to low bit depth, and would also fall below the noise floor of the sound card. To test this, we recorded the electric headphone jack output of a MacBook Pro as well as an older Asus Windows laptop, and compared the power spectrum of line noise alone to that of the white noise stimulus at the laptop volume settings corresponding to the range of participants’ reported thresholds (See Supplemental Materials and Figure S1 for full details). Power across stimulated frequencies was consistently above noise floor for all volume settings reported as white noise thresholds (from ∼+5dB to +∼14dB for MacBookPro volume setting 18 to 44%), did not change appreciably in spectral shape, and floor noise levels are consistent across volume settings. We also tested the pink noise thresholding stimulus with same RMS as the white noise (used in Expt 1d below); as would be expected, at lower frequencies (< ∼1 kHz) there was a greater difference in power between the pink noise stimulus and noise floor than with the white noise (see Supplemental Materials).

### Experimental Procedure

For all experimental conditions, sounds were presented with the Pavlovia.org (Peirce et al., 2019) online experimental platform using Google Chrome version 09.0.4430.212 via wireless connections to various broadband providers. Written instructions presented on the laptop screen asked participants to adjust the computer’s volume setting to about 50% and then to click a button labeled ‘play’ to hear the pulsed noise played on a continuous loop until the participant pressed pause or proceeded to the next page. The continuous loop was achieved by in-house JavaScript code and no gap was inserted between the repetitions. Next, participants were instructed to use the computer’s volume setting buttons on the keyboard to adjust the level of the noise so that it was barely audible. Specific instructions directed participants to slowly lower the volume setting until they could no longer hear the noise, and then to increase the volume setting one increment at a time, until they could again just barely hear the noise. After the participant was satisfied with their setting, the experimenter manually recorded the final volume setting as a percentage of full volume. As with many computers, the Mac volume setting buttons permit only a discrete range of percentage values. The only possible percentage settings were [0 6 12 19 25 31 38 44 50 56 62 69 75 81 88 94 100]. A demonstration of the procedure is available at [https://run.pavlovia.org/sijiazhao/volumechecking_demo]. The implementation is available in JavaScript [https://gitlab.pavlovia.org/sijiazhao/volumechecking_demo] and via the Gorilla experimental platform [https://gorilla.sc/openmaterials/261557].

For Expt 1a and 1b (outdoor experiments), participants only performed the task once. For Expt 1c, participants performed the task once outdoors, and then once in the anechoic chamber. For Expt 1d, participants performed four variants of the paradigm. Wearing the Beyer Dynamic DT-150 headphones, participants set levels using 1) white and 2) pink noise stimuli. They also set levels using the white noise stimulus only while wearing 3) Sennheiser HD206 headphones and 4) Apple AirPods. The order of these four variations was counterbalanced over the 20 participants.

## Results

### Experiment 1a (participants tested outdoors at Carnegie Mellon University)

Participants set their “just detectable” levels, an estimate of the audibility threshold, by choosing volume settings that were between 19 - 50%, a range that corresponds to 22.3 – 35.6 dBA SPL, with a mean dBA SPL setting of 29.43 (standard deviation (SD) 3.95, Figure 2A).

**Figure 2.**
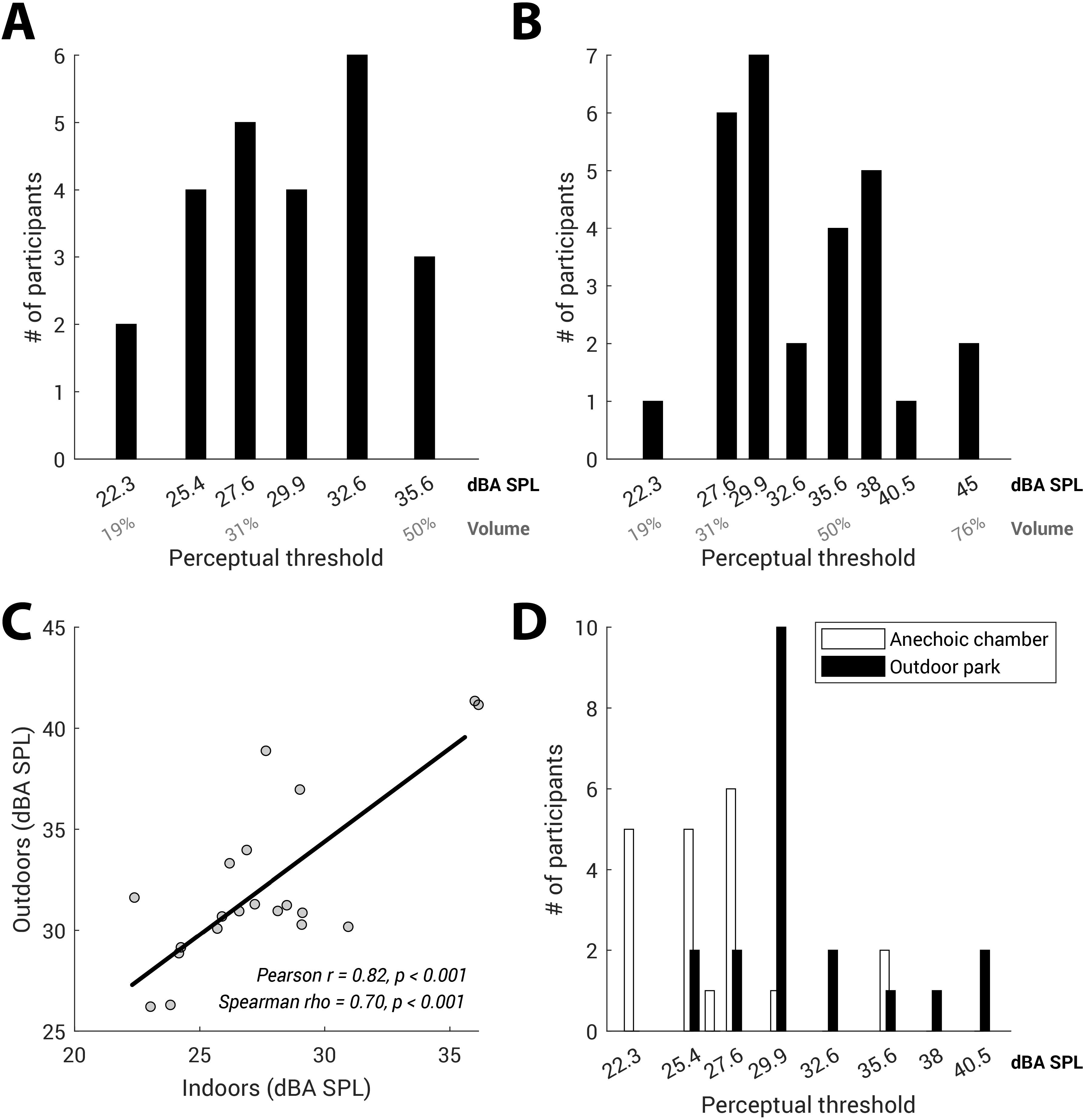
Perceptual thresholds set in Expt 1a-c. (A) Frequency histogram showing the number of Expt 1a participants who set their perceptual threshold at each volume setting/dBA SPL level, as established in the anechoic calibration procedure. The top row of the x-axis shows estimated dBA SPL level; the bottom row of the x-axis shows the range of the corresponding MacBook Pro volume settings. (B) Frequency histogram showing the number of Expt 1b participants who set their perceptual threshold at each dBA SPL. The top row of the x-axis shows estimated dBA SPL; the bottom row of the x-axis shows the range of the corresponding MacBook Pro volume settings. (C) Scatterplot showing Expt 1c data from the same participants, collected indoors in the anechoic chamber (x-axis), and outdoors in a Pittsburgh park (y-axis). The black line shows best linear fit; individual data points are slightly jittered to show all 20 individuals. (D) Frequency histogram showing the number of Expt 1c participants who set their perceptual threshold at each dBA SPL, for indoor (anechoic chamber) and outdoor (park) settings.

### Experiment 1b (participants tested outdoors in central Pittsburgh park)

Participants’ white noise perceptual thresholds were somewhat broader than in Expt 1a. Volume settings were between 19 and 76%, a range corresponding to 22.3 – 45.0 dBA SPL, and a mean dBA SPL of 33.05 (SD 5.62, Figure 2B).

### Experiment 1c (participants tested both outdoors and in psychoacoustic laboratory settings)

As with the previous experiments, participants’ white noise detection thresholds were converted from the MacBook Pro percent volume setting to dB SPL using the data and extrapolation shown in Figure 1. Results in both settings replicated the previous experiments, with participants’ indoor volume settings ranging between 19 - 50% (22.3 – 35.6 dBA SPL, mean 26.59 dBA SPL, SD 3.83), and outdoor settings ranging between 25 - 63% (25.4 – 40.5 dBA SPL, mean 31.24 dBA SPL, SD 4.31).

Figure 2C shows that participants’ noise detection thresholds in anechoic and outdoor conditions were highly correlated (Pearson r = 0.82, p < 0.001, verified using nonparametric Spearman rho = 0.70, p < 0.001). There was a modest average increase of 4.66 dBA SPL in the threshold values from anechoic to outdoor settings (Figure 2D). This mean increase in threshold seems reasonable despite the relatively large difference in ambient noise levels (31 dBA SPL indoors, and 57 dBA SPL outdoors). An inspection of the relative power spectral densities (see Supplementary Materials Figure S2) shows that while there are large differences at low frequencies, those differences are smaller near the upper end of the frequency band of the test stimulus (indicated by the shaded area). It may also be that the outdoor noise sources are relatively localizable, and thus more easily segregated from the stimulus during testing.

Because participant age can interact with both pure-tone hearing thresholds as well as listening in noise, we assessed the potential effects of age on estimated thresholds in outdoor settings by combining data from Expts 1b and 1c (Figure 3A). Using a regression analysis including age in years as well as cohort (participants in Expt 1b or Expt 1c), the overall model was significant (ANOVA, F(2,45) = 4.87, *p* < 0.0121), with no significant effect of cohort (*t* = 1.60, *p* = 0.12), and a significant moderate effect of age (*t* = 2.84, *p* = 0.0067, slope estimate 0.204). There were two people who had relatively high thresholds (45 dBA SPL); one participant (age 40) mentioned they occasionally wore hearing aids.

**Figure 3.**
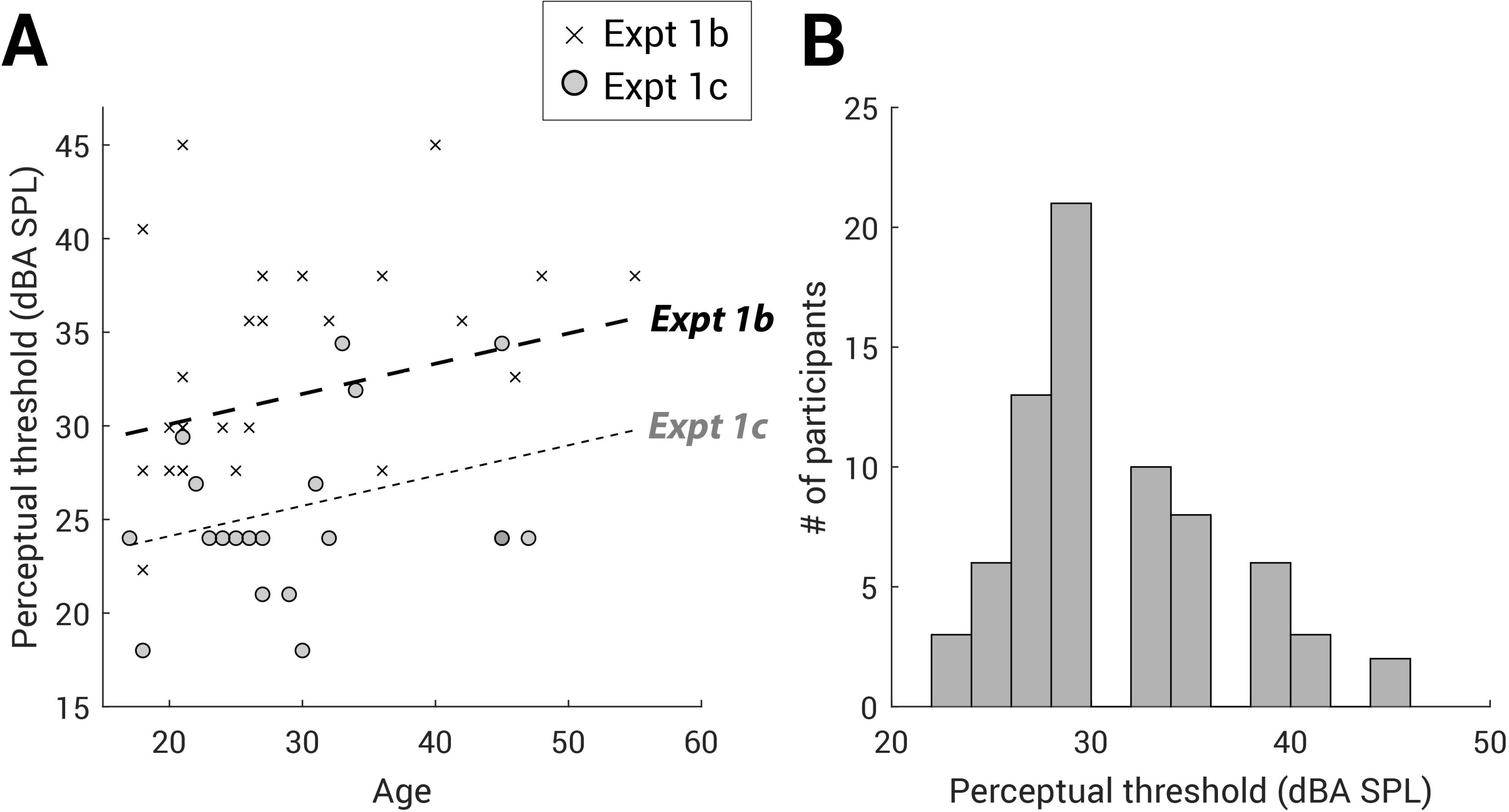
Perceptual thresholds in Expt 1a-c. (A) Scatterplot showing the relationship between Expt 1b and 1c participant age (on the x-axis) and estimated dBA SPL threshold on the y-axis. The crosses present the individual data from Expt 1b and the grey circles present the individual data from Expt 1c (two experiments N = 48 in total). The thick and thin dashed lines show the best fit between age and dBA SPL threshold when cohort (participants in Expt 1b or 1c) is included in the regression model. (B) Histogram of perceptual thresholds set by participants who were tested outdoors in all Expts 1a-1c (n=72) is shown, with the black bins indicating the number of participants who set their perceptual threshold at each dBA SPL.

Across Expts 1a-1c (N=72 total participants tested outdoors, Figure 3B), the median noise detection threshold was 29.90 dBA SPL, with the 10^th^ and 90^th^ percentiles at 25.40 and 38 dBA SPL. For very quiet indoor settings, extrapolating from the Expt 1c outdoor versus indoor within-subjects experiment showing a 4.66 dB level difference, we would expect a median detection threshold of 25.95 dBA SPL with 10^th^ and 90^th^ percentiles of 22.30 and 32.75 dBA SPL. The 25.15 dB (14.1 to 39.25 dBA SPL) range of sound detection thresholds is similar to the ∼25dB range of hearing reported for the 5th-95th percentile of normal hearing adults 18-40 years of age (Park, Yoo, Baek, Kim, & Cho, 2016); this assumes that assessment of auditory thresholds with different pure tone frequencies and 80 Hz - 8000 Hz bandpass-filtered white noise are comparable, an assumption with limited evidence, to our knowledge (Carrat, Thillier, & Durivault, 1975).

### Experiment 1d (participants tested using white and pink noise, and different headphones and earbuds)

We first compared levels set using white and pink noise while participants wore the Beyer Dynamics D-150 headphones in quiet conditions. Participants’ white noise thresholds ranged between 25-44% volume setting (25.4-32.6 dBA SPL) and were very highly correlated with their pink noise thresholds (Spearman’s rho = 0.83, p < 0.0001, see Figure 4A). There was a significant offset, where levels set with pink noise were on average one volume increment higher compared to white noise (Wilcoxon signed-rank, S=100, p < 0.0001), corresponding to a ∼2dB difference. Next, we compared white noise thresholds set when using the Beyer Dynamics D-150 versus the Sennheiser HD206 and Apple AirPods. Thresholds set with the Beyer Dynamics D-150 were significantly correlated with those set with the Sennheisers (Spearman’s rho = 0.65, p = 0.0021; Figure 4B), and with the AirPods (Spearman’s rho = 0.69, p = 0.0008; Figure 4C). Threshold volume settings were on average reliably but just slightly (0.75 volume control increments) higher with the Beyer Dynamics (mean = 33.2%) than with the Sennheisers (mean = 28.5%, Wilcoxon signed-rank, S=82.5, p < 0.001). By comparison, threshold volume settings were an average of 3.05 higher with the AirPods (mean volume setting = 52%, Wilcoxon signed-rank, S=105, p < 0.0001). As would be expected given the relatively young (18–37-year-old) cohort in this condition, there were no significant correlations between age and amplitude setting threshold (all p > 0.1).

**Figure 4.**
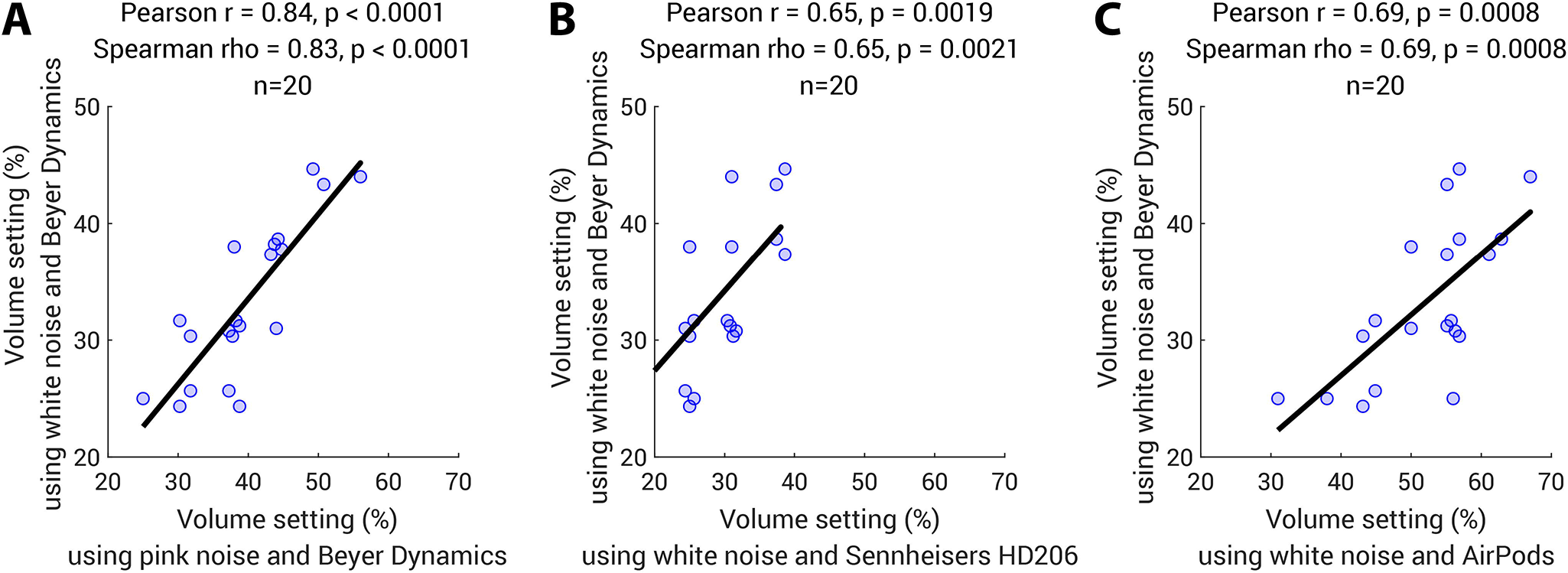
Comparison of volume using different noises and different headphones in Expt 1d (N=20). (A) Scatterplot showing the relationship between the thresholds set using white noise (y-axis) and those using pink noise (x-axis) while listeners wore Beyer Dynamics D-150 headphones in quiet conditions. The light blue circles present the individual data (N=20). A small amount of jitter (<10% of one standard deviation of the value range) was applied to the overlapping points in both x and y directions. The black line shows the best fit between two estimates. Both Pearson and Spearman’s correlations statistics are shown above the plot. (B) Scatterplot showing the relationship between the thresholds using white noise wearing Beyer Dynamics D-150 (y-axis) and Sennheisers HD206 (x-axis). (C) Scatterplot showing the relationship between the thresholds using white noise wearing Beyer Dynamics (y-axis) and AirPods (x-axis).

In sum, Expt 1 establishes the feasibility of having participants act as their own reference for setting sound levels, even under worst-case listening conditions in public outdoor spaces. Although the approach is quite a departure from the high level of control typical of laboratory studies, it presents a practical alternative for online auditory psychophysical paradigms in which stimulus amplitude must fall within a constrained range of audibility.

## Experiment 2

Experiment 2 makes use of the noise detection threshold setting procedure, validated in Expt 1, to set stimulus levels for a classic psychophysical task -- tone detection in noise -- among online participants. We first ask if reliable, well-behaved psychophysical threshold tracks can be obtained online. Second, we examine whether small adjustments to traditional threshold-setting procedures might permit fast (1-3 minutes) and reliable threshold estimates online. Given the risk of reduced participant vigilance and attentiveness during online studies, minimizing the amount of time devoted to establishing a psychophysical threshold is particularly important. Thus, the first goal of Expt 2 is to investigate the minimum number of trials needed to derive a reliable threshold estimate. Modern online testing platforms also make the study of human psychophysics available to a wide cross-section of would-be researchers, including students and other non-experts. In this light, another goal of Expt 2 is to determine whether the standard method of estimating a threshold -- the mean across a set number of reversals -- can be simplified while still upholding high psychophysical standards. We examine whether a simple estimate of the mode across all levels encountered in the staircase procedure is as robust or more robust at estimating threshold as traditional estimators based on staircase reversals. This adds to previous efforts to optimize the efficiency and precision of auditory threshold setting techniques (e.g., Dillon et al., 2016; Gallun et al., 2018; Grassi & Soranzo, 2009). A third goal of Expt 2 is to ask whether individual differences in threshold levels might be influenced by online participants’ arousal, engagement, or fatigue (Bianco et al., 2021; Libera & Chelazzi, 2006; Shen & Chun, 2011). To this end, we surveyed these characteristics at multiple timepoints during the threshold setting procedures.

## Methods

### Participants

60 online participants took part via the Prolific recruitment platform (prolific.co, Damer and Bradley, 2014; see Table 2 for demographics); all gave electronic informed consent prior to the experiment, with ethical approval granted by the Birkbeck College Psychological Sciences ethics committee (see Expt 1). Data collection occurred between 11^th^ and 14^th^ May 2021 with participants paid to complete the study.

**Table 2.**
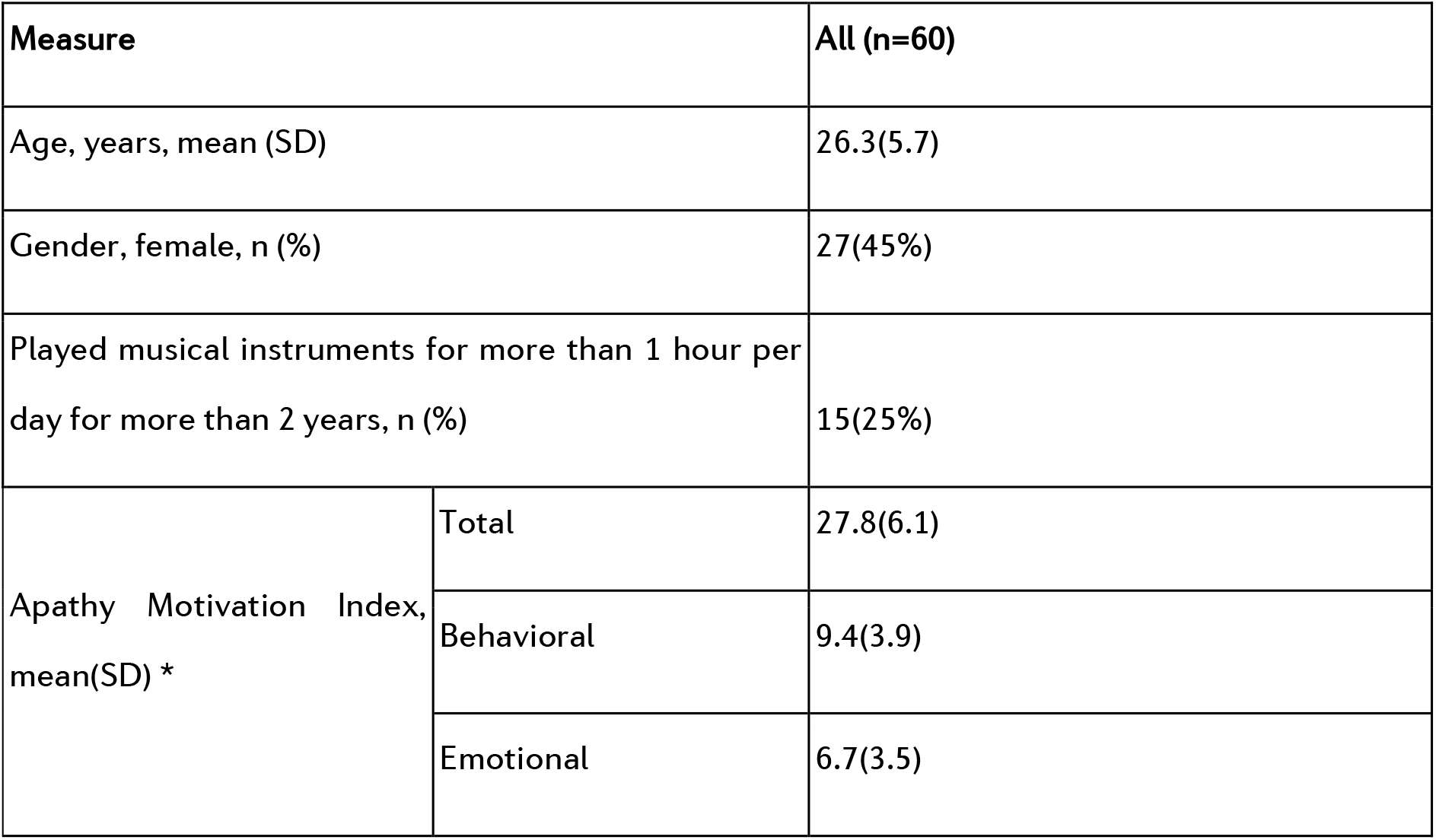
Self-reported participant demographics. *One participant did not complete the Apathy Motivation Index questionnaire.

Participants were selected from a large pool of individuals from across the world. As Prolific is available in most of OECD countries except for Turkey, Lithuania, Colombia and Costa Rica and also available in South Africa, most prolific participants are residents in these countries. In our sample, the 60 participants were residents from 13 different countries including United States, United Kingdom, Canada, Poland, Spain, Germany, South Africa, Belgium, Chile, Mexico, Portugal, France and New Zealand. We utilized Prolific.co pre-screening options to refine eligible participants to those who were between 18 and 40 years of age, reported no hearing difficulties, and had a 100% Prolific.co approval rate. 91 participants began the experiment online, and of these, 31 dropped out either before or after the headphone test (see below), or during the main experiment.

### Stimuli and Procedure

The experiment was implemented using PsychoPy v2021.1.2 and hosted on PsychoPy’s online service, Pavlovia (pavlovia.org). A demo is available at [https://run.pavlovia.org/sijiazhao/threshold_demo]. Participants were required to use the Chrome internet browser on a laptop or desktop computer (no smartphone or tablet) to minimize the variance in latency caused by differences among browsers and devices. Operating system was not restricted. Before the start of the online experiment, participants were explicitly reminded to turn off computer notifications.

### Amplitude Setting

Participants first followed the amplitude setting procedure described for Expt 1. As described above, this brief (<2 min including form-filling) procedure had participants adjust the volume setting on their computer so that the stimulus was just detectable, thereby serving as their own level reference.

### Headphone Check

After that, we screened for compliance in wearing headphones using the dichotic Huggins Pitch approach described by Milne et al. (2020). Here, a faint pitch can be detected in noise only when stimuli are presented dichotically, thus giving higher confidence that headphones are being worn. The code was implemented in JavaScript and integrated with Pavlovia using the web tool developed by author SZ [https://run.pavlovia.org/sijiazhao/headphones-check/].

The headphone check involved 6 trials, each with three one-second-long white noise intervals. Two of the intervals presented identical white noise delivered to each ear. The third interval, random in its temporal position, was a Huggins Pitch stimulus (Cramer & Huggins, 1958) for which white noise was presented to the left ear and the same white noise, phase shifted 180° over a narrow frequency band centered at 600 Hz (±6%), was presented to the right ear to create a Huggins Pitch percept (Chait, Poeppel, & Simon, 2006; Yost & Watson, 1987).

Participants were instructed that they would hear three noises separated by silent gaps and that their task was to decide which noise contained a faint tone. Perfect accuracy across six trials was required to begin the main experiment. Participants were given two attempts to pass the headphone check before the experiment was terminated. The procedure took approximately 3 minutes to complete.

To get an overall idea of attrition, we counted how many participants returned the test on Prolific. A total of 91 participants started the test, 7 participants quit the test after passing the headphone test, and 24 returned the test before the main experiment started. However, of these 24 returned participants, it is unclear whether they completed the headphone test or not, as they might have quit even before the headphone test started. Nevertheless, our total attrition for Experiment 2 (including before or after the headphone test and drop-out during the main experiment) is 34.1% (31/91).

### Adaptive Staircase Threshold Setting Procedure

Two simple acoustic signals comprised the stimuli for the adaptive threshold setting procedure. A 250-ms, 1000-Hz pure tone with 10-ms raised-cosine amplitude onset/offset ramps was generated at a sampling rate of 44.1kHz (16-bit precision) in the FLAC format using the Sound eXchange (SoX, http://sox.sourceforge.net/) sound processing software. This tone served as the target for detection in the threshold setting procedure.

A 300-sec duration white noise with 200-ms cosine on/off ramps served as a masker; this was generated using the same procedure as described for Expt 1, except that it was adjusted in amplitude to 0.0402 RMS rather than 0.000399 RMS as in the amplitude setting experiment (Expt 1). The white noise masker was thus 40 dB suprathreshold (20 * log(.0402 / .000399) = 40.07. To estimate the sound pressure level of the masker as delivered to Expt 2 participants, we averaged Expt 1c’s indoor and outdoor extrapolated dBA SPL (mean 22 dBA SPL, SD 4.3) and added 40 dB, arriving at an estimate of 66 (±4.3) dBA SPL average masker intensity. This is similar to many probe signal experiments, including the original Greenberg & Larkin (1968) study (65 dBA SPL), as well as a recent replication and extension (65 dBA SPL, Anandan et al., 2021).

The noise masker was continuous, with onset commencing as soon as participants began the threshold procedure and looping until the end of the experiment. At the end of each five-minute loop, there was a slight ‘hiccup’ as the noise file reloaded which occurred at different times for each participant, as several of the experimental parts were self-paced. Simultaneous presentation of a long masking sound - or indeed any long continuous sound - is challenging for experimental presentation software, particularly online. However, transient noise onsets and offsets - for instance, starting and stopping the noise masker for each trial - can have surprisingly large effects on perception, with Dai & Buus (1991) showing that use of noise bursts versus continuous noise maskers essentially eliminates the probe signal effect (Dai & Buus, 1991).

The staircase threshold procedure followed the headphone check. The threshold procedure trial design is shown in Figure 5. Each trial was a three-interval forced choice: the 1000-Hz signal tone could appear during any one of the three 250-ms response intervals with equal probability. Response intervals were separated from each other by 250 ms. The intervals were labelled with the digits ‘1’, ‘2’ and ‘3’ displayed visually at the center of a screen and participants responded using their computer keyboard by pressing the number corresponding to the interval in which they heard the signal. All symbols and instructions were presented as black text on a white background.

**Figure 5.**
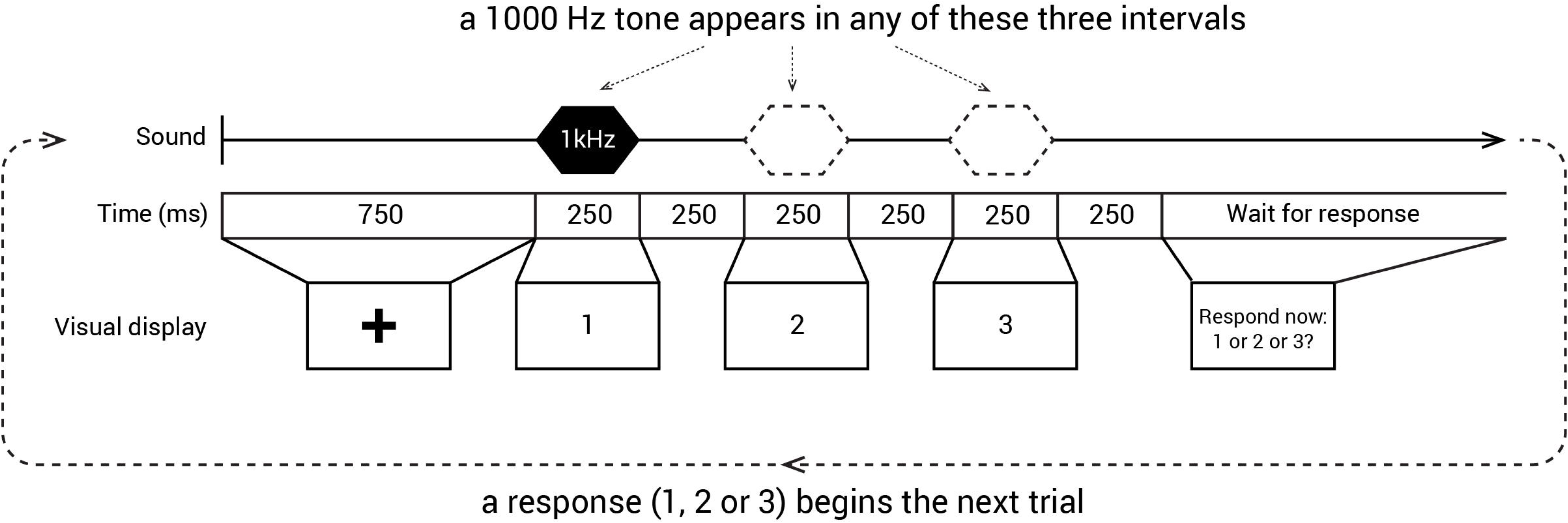
Trial structure in the threshold staircase procedure. In Expt 2, only one of the three intervals (1, 2, or 3) contained the signal, a 250 ms, 1 kHz pure tone. Responses were collected by participants pressing the corresponding numerical key on their computer keyboards.

The level of the signal relative to noise that was required to produce 79.4% correct detection was determined using an adaptive ‘three-down, one-up’ staircase procedure (Levitt, 1971). The procedure started at a signal-to-noise ratio (SNR) of −13.75 dB (calculated as dB difference in RMS between the background white noise and pure tone). Each track began with an initial descent to approximate threshold, with every correct response leading to a decrease in signal intensity by 1.5 dB with the decrement reducing to 0.75 dB once the level fell below −19.75 dB SNR or after the first incorrect response. At this juncture, the three-down, one-up staircase procedure started.

As practice before the first of three adaptive threshold staircase tracks, participants completed six trials with the signal presented at −13.8 dB SNR (i.e., the easiest level) and with performance feedback provided (“correct” or “wrong” shown for 1 sec on-screen after each response). The average performance of this practice block was 92.78% correct (SD = 13.85%) with 41 out of 60 participants (68%) making no mistakes. No feedback was given during the adaptive staircase threshold session.

Each of the subsequent three adaptive staircase threshold tracks consisted of 40 trials. Tracks were completed consecutively, with the opportunity for a short break between tracks. However, most participants did not take a break (mean break duration = 9.03 s, SD = 11.68 s).

To keep participants engaged throughout the procedure, progress was shown on the top left of the screen (“Progress: x/40”, where *x* is the index of the current trial). Moreover, we awarded a bonus (maximum of £1.50) in addition to the base payment; after the 6 practice trials with feedback, participants were informed that if their accuracy surpassed 50%, they could earn a bonus of 50p per track. The accumulated bonus was shown at the end of each track, and all 60 participants got the full bonus of £1.50.

The threshold staircase procedure was achieved using in-house code [https://gitlab.pavlovia.org/sijiazhao/threshold_demo].

### Assessment of participant apathy, motivation, and fatigue

To measure lack of motivation (apathy), we presented the Apathy Motivation Index (AMI) questionnaire before the experiment. This 18-question survey is subdivided into three apathy subscales: emotional, behavioral and social apathy (Ang, Lockwood, Apps, Muhammed, & Husain, 2017; see Supplemental Materials for questions).

To track the dynamics of motivation and fatigue across the experiment, participants also rated their level of subjective motivation, fatigue, and confidence before and after the threshold session. They were provided with three horizontal visual analogue scales, each with equally spaced tick marks along its axis, an accompanying question positioned centrally above, and labels at the extreme left and right of the scale. The questions and labels are available in Supplemental Materials.

Responses were registered by a click on the appropriate position on each scale. After completing all three ratings, a ‘confirm’ button appeared at the bottom of the screen, allowing participants to submit their ratings.

The questionnaire and ratings were added to the experiment on the second day of data collection. Thus, of the 60 participants, 49 responded to both the AMI questionnaire and ratings of motivation and fatigue, 10 had the AMI questionnaire only, and a single participant completed neither the questionnaire nor the ratings.

On average, participants spent 39.3 minutes (SD = 10.4) on the entire experiment, including both the Adaptive Staircase Threshold procedure (Expt 2) and the Probe Signal procedure (Expt 3, below).

## Results

### Reliability of individual participant signal-to-noise thresholds in online psychophysical staircase procedure

First, we asked whether we could obtain good-quality and stable tone-in-noise thresholds online. As an initial qualitative approach, we examined the three 40-trial tracks for each participant. We found that they were generally well-behaved in terms of reaching a stable plateau with multiple reversals after the initial descent to the first error. (All threshold tracks are available at https://github.com/sijiazhao/TPS_data). To estimate threshold distribution and reliability across tracks, we calculated the mean and range of thresholds for each participant, based on the last six reversals for each of the three tracks unless that track had fewer than six reversals. (Mean reversals across tracks was 7.8. Sixteen participants had one track with fewer than 6 reversals: 2 tracks with 3 reversals, 1 track with 4 reversals, 13 tracks with 5 reversals). The mean SNR threshold was −19.54 (SD = 1.39), with the distribution of mean thresholds slightly skewed toward lower SNR levels (see Figure 6A). The mean range of estimated thresholds across the three tracks was 1.71 dB (see Figure 6B); with a 10^th^ and 90^th^ percentile range of 0.39 to 3.41 dB SNR. A repeated-measures ANOVA on the mode-based threshold for each track showed no significant order effect [F(2,118) = 2.30, p = 0.11, partial eta squared = 0.038, observed power = 0.46, no significant violations of sphericity, so sphericity assumed].

**Figure 6.**
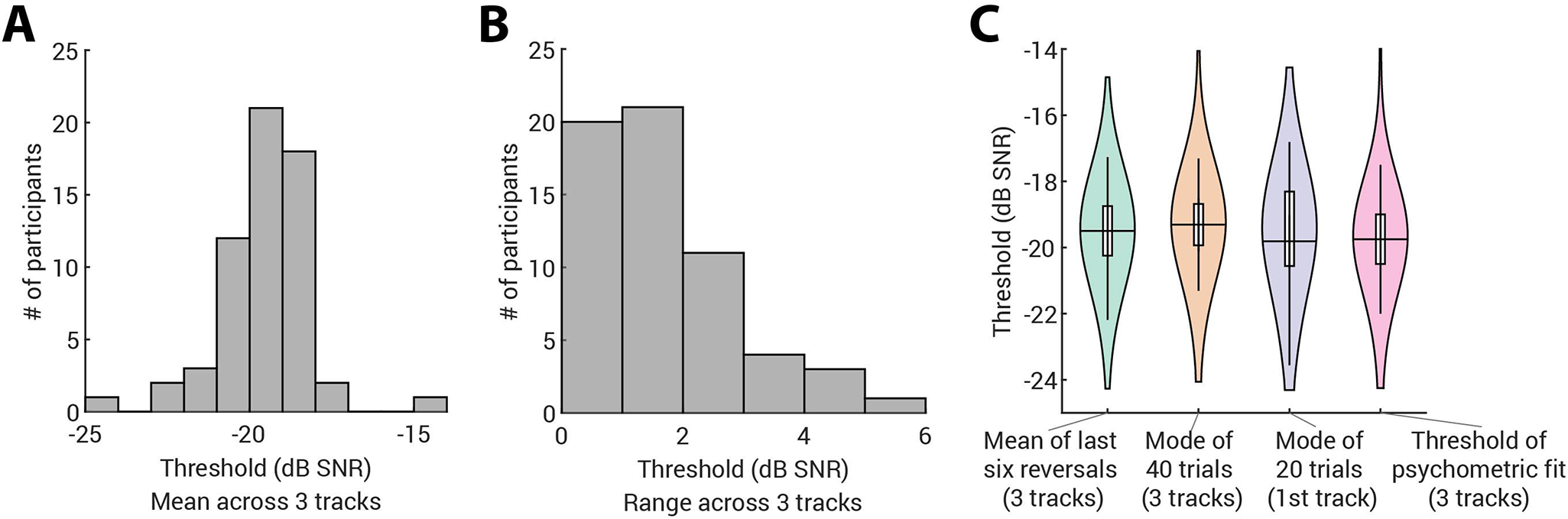
Results from Expt 2. (A) Frequency histogram of all participants’ tone-in-noise thresholds in dB SNR based on the mean of the SNR values of the last six staircase reversals (count on y-axis). (B) Frequency histogram showing the distribution of the range of 6-reversal-based thresholds across the three thresholding tracks (in dB SNR). (C) The violin plots for the tone-in-noise detection thresholds across 60 participants estimated using the four estimation methods. Each violin is a kernel density plot presenting the distribution of the estimated thresholds for each estimate method. For each violin plot, the group’s median (the horizontal black line inside the violin), interquartile range (the vertical box) and 95% confidence interval (the vertical black line) are shown.

### Evaluation of mode-derived thresholds compared to reversal counting

We compared four different methods of deriving a threshold from psychophysical data collected in the 3-down/1-up adaptive staircase procedure. The goals were: 1) to determine whether reliable threshold estimates could be obtained using fewer trials; 2) to examine whether the statistical mode is a viable alternative to the standard approach (the mean across a predetermined number of reversals).

One approach to establishing a threshold is to average values at the last six reversals in each of three tracks, and to compute a grand mean ‘gold standard’ threshold for each participant from these three-track means (green violin in Figure 6C). Another is to estimate a threshold from the psychometric function reconstructed from all 120 trials using maximum likelihood procedures carried out in the *psignifit* toolbox in MATLAB (Schütt, Harmeling, Macke, & Wichmann, 2016; pink violin plot in Figure 6C). We also calculated the statistical mode for all 40 trials in each of the three tracks per participant, and generated a grand mean from these three modal values for each participant (orange violin in Figure 6C). The rationale for using the mode is that it can be thought of as a measure of the ‘dwell time’, e.g., how long a participant spends at a particular level in the adaptive staircase procedure. Finally, we computed the mode from the first 20 trials in each participant’s first track in order to assess the goodness of a mode-based threshold estimate from a single short track (purple violin in Figure 6C). On average, the number of reversals when the 20^th^ trial was reached in the first track was 3.5 (SD = 1.0).

We compared these four metrics using a Bayesian repeated measures ANOVA in JASP (JASP Team, 2020; Morey and Rouder, 2015; Rouder et al., 2012), which revealed a very low Bayes factor compared to the null hypothesis (BF_10_ = 0.592), as would be expected given the ≤ 0.2 dB SNR mean difference between any of the four metrics. This suggests that there is little, if any, significant bias in using either modal measure versus the more standard approaches.

However, a potentially more consequential difference between obtaining a single 20-trial threshold track estimate versus using the three-track 40-trial 6-reversal-based estimate would be unacceptably high variability in the former case. To quantify the degree of variability associated with the number of trials used to calculated the threshold, we compared the distributions of differences between the 3-track grand average and single-track thresholds calculated using the mode of 1) the first 20 trials; 2) or 30 trials; 3) all 40 trials; or 4) the mean of last six reversals. Each participant contributed 3 difference scores (one per track) to each distribution. Figure 7 shows the range of deviation from the gold-standard that is observed when using mode-based estimation. As would be expected, dispersion decreases as more trials are used to calculate the threshold.

**Figure 7.**
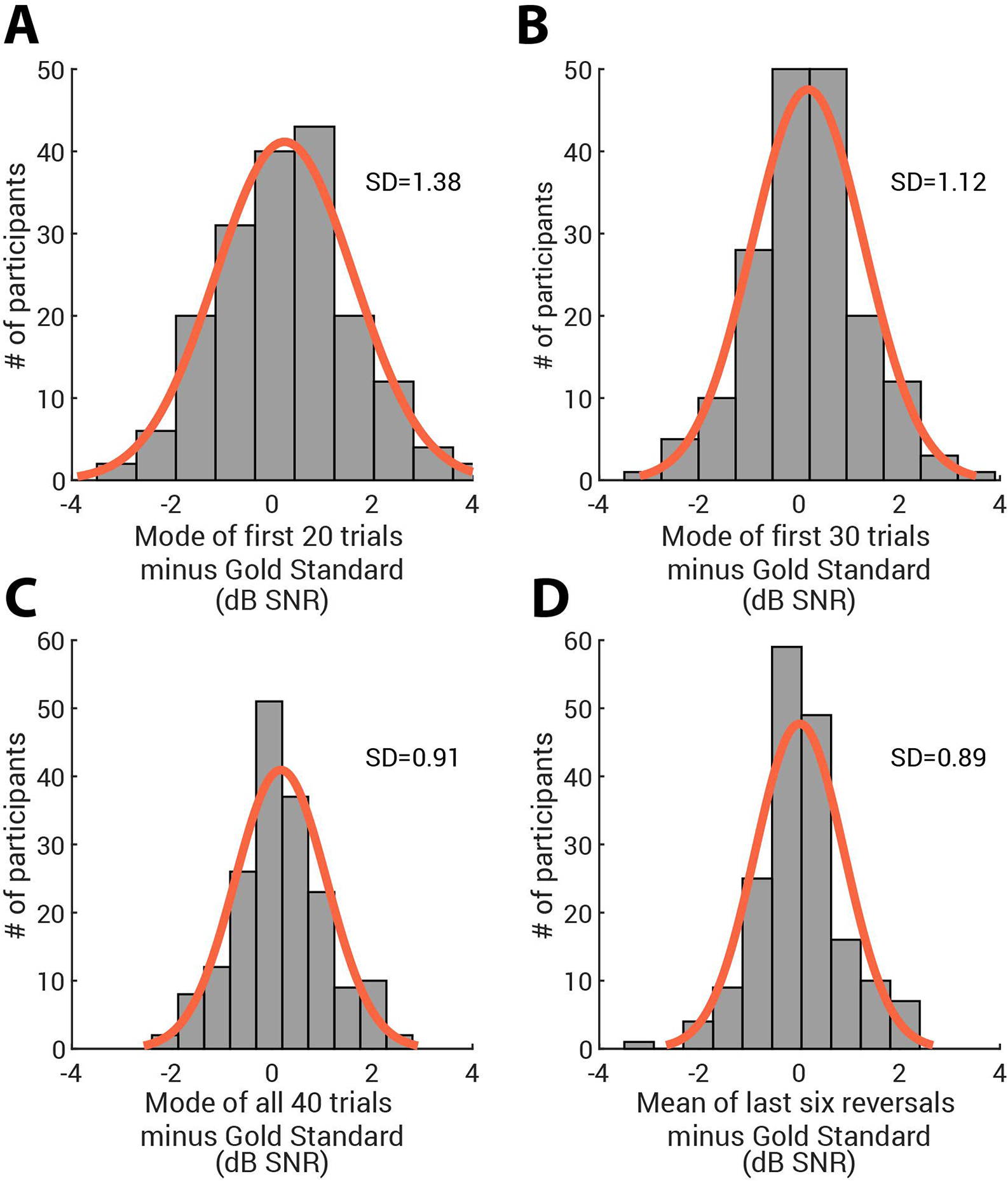
Difference in dB SNR of each participant’s single tone-in-noise threshold tracks derived from the mode of the first 20 (Fig 7a), 30 (Fig 7b), and all 40 trials (Fig 7c) when compared to the ‘gold standard’ mean of three reversal-based thresholds. As a comparison, Fig 7d shows the analogous difference between the gold standard mean, and the track-wise mean of the last six reversals. Note that each participant contributes three datapoints (one from each track) to each distribution.

We also assessed the adequacy of single-track mode-based threshold estimates using the initial 20, 30, or all 40 trials. To do so, we examined the correlation of each mode-based threshold with the 3-track threshold across participants, and then statistically compared the difference in correlations. As tested using the r package *cocor* using the Hittner et al. and Zou tests (Diedenhofen & Musch, 2015; Hittner, May, & Silver, 2003; Zou, 2007), the fit between the gold standard and mode-based thresholds differed across tracks ^2^ (Figure 8, *r-values* shown in figure). Here, the correlations between each mode-derived threshold from first thresholding track and the gold standard threshold were all significantly (p < 0.05) lower compared to when the same measure used data from the second thresholding track. Correlation differences between the first and third tracks were in the same direction, but ‘marginal’ using the Hittner et al. tests (p < 0.08). The less-robust thresholds obtained in the first track suggest that at least some psychophysics-naive online participants had not quite acclimated to the threshold setting procedure until later on in the track.

**Figure 8.**
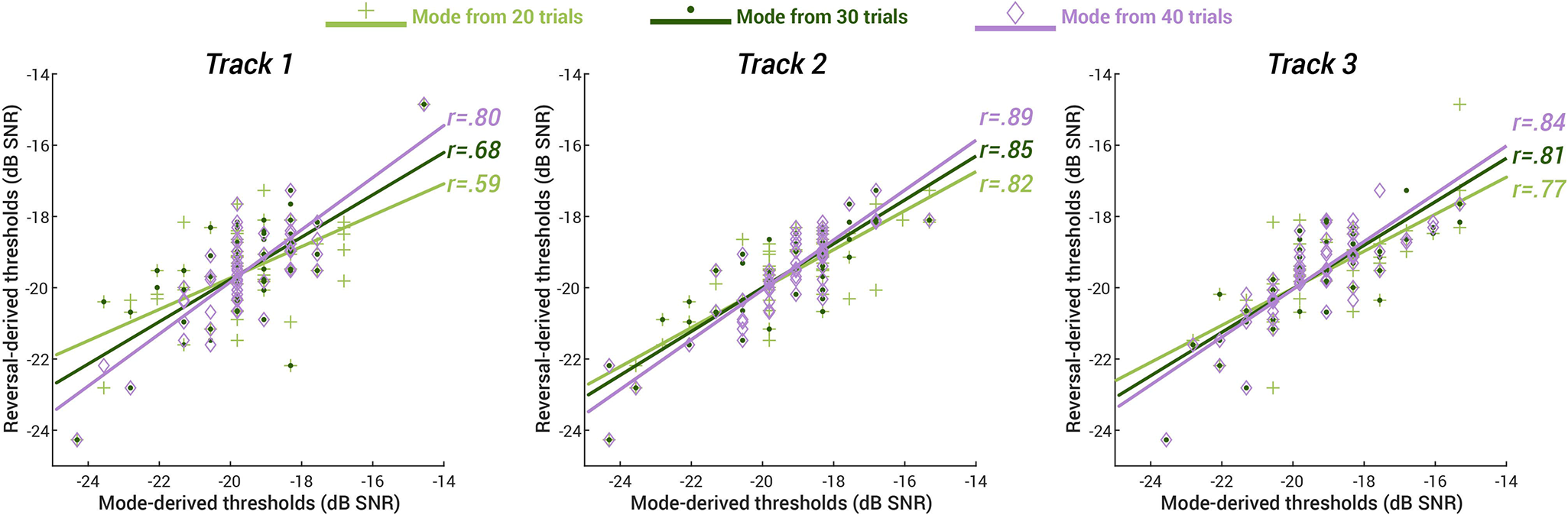
Correlations between the 3-track gold standard threshold (y-axis) and single-track threshold estimates (x-axis) based on the mode of the first 20, 30, or all 40 trials (track 1 (left), 2 (middle), and 3 (right panel)). Light green crosses and lines refer to 20-trial mode estimates, dark green to the 30-trial estimates, and purple to the 40-trial estimates. Both axes show tone-to-noise dB SNR. Pearson’s correlation coefficients for each estimate are shown on the right of the fits.

Using the same difference-in-correlation-based comparison method (and with the same statistical caveats), we also found that the relative reliability of mode-based thresholds derived from 20 or 30 versus 40 trials changed across tracks. In the first and second tracks, thresholds based on the first 20 trials were significantly less correlated with the gold standard than were those based on 40 trials (p < 0.05) but did not differ in the last track; correspondingly, first-track thresholds based on the first 30 trials were significantly less correlated with the gold standard than were those based on 40 trials (p < 0.05), but this difference was no longer significant in the second or third tracks. In addition, the overall deviation of mode-derived scores from the gold-standard approach (the standard deviation of the threshold differences; SD in upper-right corner of each panel in Figure 7) decreases with increasing number of trials, indicating a convergence of the mode-based threshold approaches toward the gold standard. A reasonable explanation for this effect is that online participants acclimated to the threshold setting procedure across the three tracks, and performance became more stable and consistent after a few minutes of practice. Nevertheless, as shown previously (Figure 8) even tone-in-noise thresholds based on the first 20 trials in the first track are reasonably accurate estimates of a participant’s ‘true’ threshold.

### Evaluation of potential motivation, confidence, and fatigue effects on tone-in-noise thresholds

Here, we asked whether estimated thresholds might in part reflect the personal motivation of online participants. To this end, we used a common self-report for a personality trait-like component of motivation among healthy populations (apathy in the AMI questionnaire, Ang et al., 2017). We also examined the dynamic change of motivation ratings across our task, measured before the first threshold track and again after the third threshold track.

Participants’ tone-in-noise thresholds from did not correlate with any aspect of the motivation trait measured by the AMI questionnaire. Neither behavioral (rho = .055, *p* = .68), emotional (rho = .079, *p* = .55), nor social apathy (rho = .053, *p* = .69) dimensions were related to tone-in-noise thresholds. Self-reported motivation across the course of the staircase thresholding procedure also did not account for threshold level either before (rho = −0.15, *p* = 0.29) or after (rho = 0.007, *p* = 0.96) the threshold procedure.

To assure ourselves that this lack of correlation was not due to a faulty instrument, we tested whether there was a correlation between the trait motivation/apathy score and the in-experiment motivation ratings. Indeed, the behavioral dimension of the apathy questionnaire was associated with the post-experiment motivation level (rho = −0.37, *p* = 0.010) and this relationship remains significant after controlling for the threshold level (partial correlation, *r* = −0.39, *p* = 0.006). This indicates that more apathetic individuals reported feeling less motivated after the threshold session regardless of their behavioral performance, although no relationship was observed prior to the experiment.

The absence of a link between motivation and task performance was further confirmed by a repeated measures general linear model on the tone-in-noise threshold level with fixed effects of the total score of the apathy questionnaire, the pre-threshold and the post-threshold motivation ratings. The thresholds could not be predicted by apathy traits (F(1,41) = 0.022, *p* = 0.88), or motivation ratings either pre-experiment (F(1,41) = 0.93, *p* = 0.34) or post-experiment (F(1,41) = 0.15, *p* = 0.71). Moreover, there were no three-way or two-way interactions (all *p* > 0.32). In sum, online participants’ motivation did not contribute significantly to their tone-in-noise thresholds.

Threshold level was also not significantly related to confidence measured either before (rho = 0.074, *p* = 0.61) or after the experiment (rho = −0.12, *p* = 0.40), suggesting that participants showed quite limited metacognitive awareness of their performance.

Finally, we investigated the relation of self-reported fatigue to thresholds. Here, the threshold level *did* positively and moderately correlate with fatigue ratings both before (rho = 0.31, *p* = 0.027) and after (rho = 0.32, *p* = 0.025) the experiment, consistent with higher (poorer) thresholds among fatigued participants.

In all, Expt 2 demonstrates that it is possible to quickly and reliably estimate a classic auditory psychophysics threshold online. Moreover, a very simple -- and easily automatized -- estimate of the level at which participants dwell for the most trials across the adaptive staircase procedure (the mode) is highly reliable, and as robust at estimating threshold as traditional estimators based on staircase reversals. We outline potential usage cases regarding the number of tracks and trials to use in the Discussion. Finally, online participant motivation level is not a significant moderator of tone-in-noise perceptual threshold (at least within the range of motivation levels and task difficulty we measured here), whereas fatigue was associated with somewhat poorer tone-in-noise detection.

## Experiment 3

Experiment 3 tests an online version of the classic probe signal paradigm to measure frequency-selective auditory attention (Borra et al., 2013; Dai & Buus, 1991; Dai et al., 1991; Green & McKeown, 2001; Greenberg & Larkin, 1968; Macmillan & Schwartz, 1975; Moore et al., 1996; Scharf et al., 1987). We ask 1) whether the Expt 2 online tone-in-noise threshold-setting procedure is sufficient for setting the SNR level to achieve a specific target accuracy in the 2AFC tone detection task used in the probe signal paradigm. We then ask 2) whether this paradigm can be replicated online in relatively uncontrolled environments; 3) if frequency-selective attention effects can be observed on an individual basis within a single short online testing session (circa 30 minutes); and 4) if these effects change across the course of a testing session. As with Expt 2, we finally ask 5) whether psychophysical thresholds and frequency-selective attention are related to well-established measures of fatigue, apathy, and task confidence before, during, or after testing.

## Methods

### Participants

All participants from Expt 2 also took part in Expt 3.

### Stimuli and Procedure

Like Expt 2, Expt 3 was implemented using PsychoPy v2021.1.2 and hosted on PsychoPy’s online service, Pavlovia (pavlovia.org). A demo is available at [https://run.pavlovia.org/sijiazhao/probesignal_demo]. All experimental restrictions used in Expt 2 also applied in Expt 3.

After completing the Amplitude Setting, Headphone Check, and Threshold Setting of Experiment 2, participants completed a classic probe-signal task (Anandan et al., 2021; Botte, 1995; Dai & Buus, 1991; Dai et al., 1991; Greenberg & Larkin, 1968; Scharf et al., 1987; Tan et al., 2008). Continuous broadband noise was present throughout all trials, as described for the threshold setting procedure.

As shown in Figure 9, each trial began with a 1000-Hz, 250-ms cue tone followed by 500 ms of silence. At this point, the first of two listening intervals was indicated by a black ‘1’ presented at central fixation on the white computer screen for 250 ms. The ‘1’ disappeared during a 250-ms silent interval at which time a black ‘2’ was presented at fixation to indicate a second listening interval.

**Figure 9.**
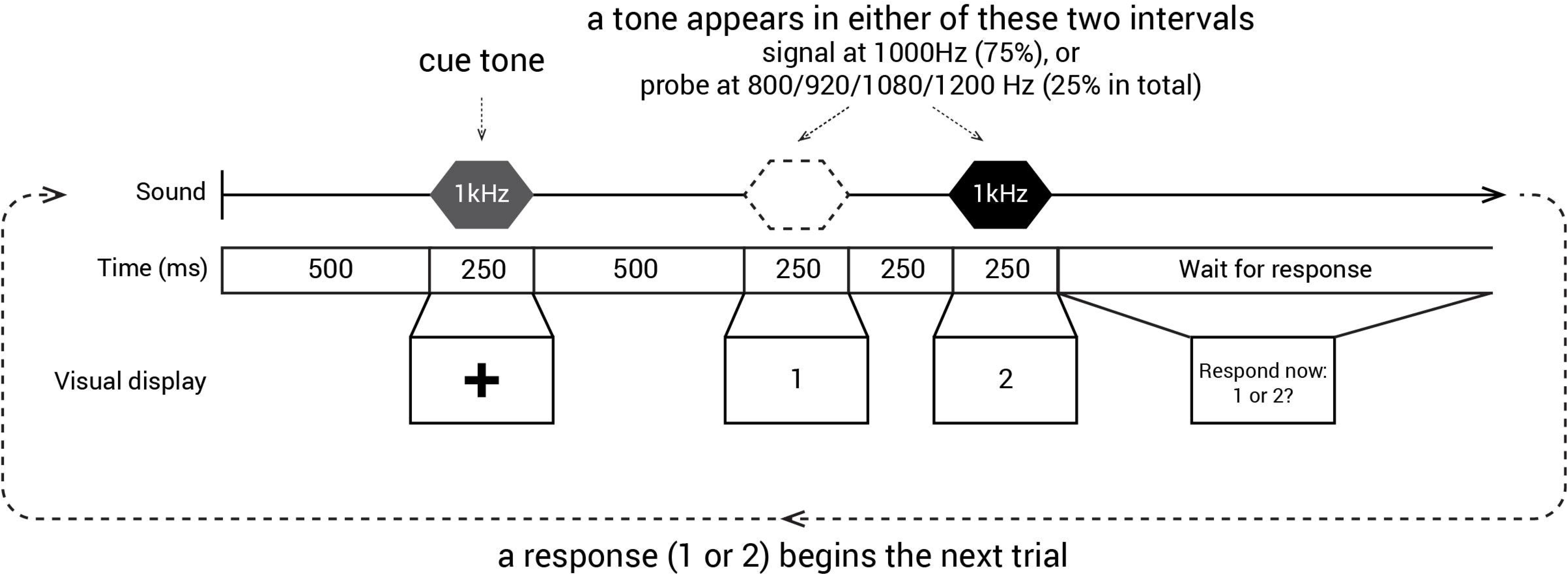
Trial structure in the probe signal task. In Expt 3, one of the two intervals (1, 2) contained the signal, a 250 ms, 1 kHz pure tone. Responses were collected by participants pressing the corresponding numerical key on their computer keyboards.

A 250-ms tone was presented with equal probability in either the first or the second listening interval; participants reported which interval contained the tone with a keypress. *Signal* trials involved a tone that matched the 1000-Hz cue frequency; these trials comprised 75% of the total trials. Another four *probe* tones with 800, 920, 1080, and 1200 Hz frequencies were presented with equal probability across the remaining 25% of trials (6.25% likelihood for each tone frequency).

To assure ourselves that the full sample did not perform at ceiling, we adjusted each individual’s probe-signal SNR threshold slightly, lowering it by one step size (0.75 dB) from the threshold estimated in Expt 2. The signal and probe tones were always presented at the adjusted threshold level; the preceding cue tone was suprathreshold, set at 14 dB above the adjusted threshold SNR level.

Participants first completed five practice trials with suprathreshold signal and probe tones presented at −13.8 dB SNR. Immediately thereafter, another five practice trials involved signal and probe tones at the adjusted individual threshold. Performance feedback (‘correct’ or ‘wrong’) was provided on-screen for one second following each response to a practice trial.

Each of the subsequent 12 blocks consisted of 32 trials (384 trials total), with 24 signal trials (1000-Hz tone) and 2 probe trials at each of the other frequencies (8 probe trials total) in random order. Blocks were completed consecutively, with the opportunity for a short break between blocks (mean break duration = 10.44 s, SD = 22.44 s). There was no feedback for these trials.

Participants were informed that if their overall accuracy across the 12 blocks surpassed 65%, they would earn a bonus of £1.00 at the end of the experiment. In all, 63% of participants earned the bonus.

## Results

### Adequacy of online thresholding for setting SNR levels for 2AFC task

We first asked how effective the online tone-in-noise threshold measurement was in setting the SNR level for the probe signal task. The adaptive staircase procedure (3-down, 1-up) was designed to set the threshold to detect a 1000-Hz tone in noise at 79.4% accuracy. However, to retain additional ‘head room’ for accuracy in the probe signal task we lowered the actual SNR level by 0.75 dB for each individual (as noted above). In order to map how changes in tone-in-noise SNR levels mapped to changes in 2AFC tone-in-noise detection accuracy, we ran a small study and found that each 0.75 dB increment in SNR corresponded to a detection accuracy change of 4.2%. Thus, if the Expt 2 online threshold setting functioned correctly, Expt 3 participants should achieve tone-in-noise detection of 75.2%. As shown in Figure 10A, average signal detection accuracy was 72.45% (SD = 8.86), just slightly (2.75%) yet significantly lower than the predicted accuracy (t(59) = 62.67, *p* < 0.001, BF > 10^50^).

**Figure 10.**
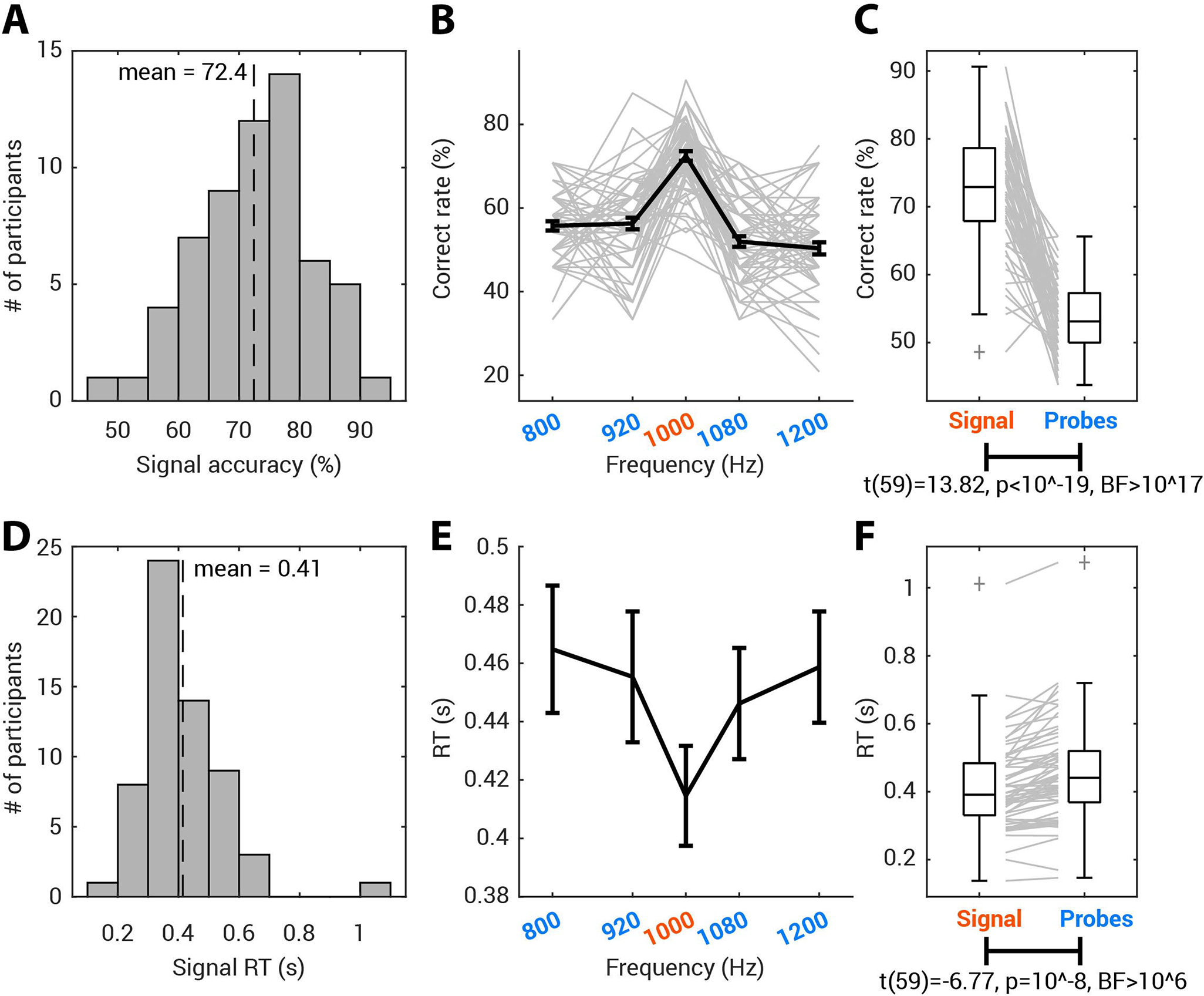
Experiment 3: Probe-signal (N=60). (A) Distribution of the signal accuracy using the mode-derived threshold. The population mean is labelled as a dashed vertical line. (B) The percent correct detection of 1000-Hz signal and each of the four probes (800, 920, 1080 and 1200 Hz). The thick black line presents the group mean, with error bar = +/- 1 SEM. Each grey line indicates individual data. (C) The accuracy to detect signals (highly probable 1000Hz tones) was significantly higher than the average detection accuracy for probe tones (the less probable 800, 920, 1080, and 1200Hz tones) The population data is presented as a boxplot with the outliers marked as grey crosses. Each grey line indicates individual data, and paired t-test stats reported below the graph. The RT data is shown in the same manner below, in panels D, E, and F. For the visualization, individual data are not presented in (E), but a summary of individual data is shown in (F).

If the Expt 2 mode-derived threshold adequately estimated tone-in-noise thresholds, then a participant’s tone-in-noise detection accuracy in Expt 3 should be independent of their tone-in-noise threshold. In other words, even if two participants have very different tone-in-noise thresholds, their accuracy on the 2AFC probe-signal task should be more or less equivalent. Indeed, probe-signal detection accuracy was not correlated with the mode-derived threshold level (Spearman rho = −0.08, *p* = .544; Pearson *r* = −.01, *p* = .929).

### Robustness of the probe-signal effect at group and individual level

As shown in Figure 10B, online participants detect the high-probability 1000-Hz signal at levels that are approximately at the predicted target accuracy (72.45% (SD = 8.86), Figure 10A and 10C), whereas tones with less-probable frequencies are much less accurately detected (53.59% (SD = 5.36), Figure 10C). Figure 10C plots a direct comparison of what is visually apparent in Figure 10B. The signal tone was detected significantly more accurately than were probe tones (*t*(59) = 13.82, *p* < .00001, BF > 10^17^; Figure 10C). This classic pattern of frequency-selective auditory attention is echoed in faster reaction times for the 1000-Hz signal tone compared to the probe tones (*t*(59) = 6.77, *p* < .00001, BF>10^6^; Figure 10E, 10F). These results replicate the frequency-selective attention effects that have been documented in laboratory studies for decades (Anandan et al., 2021; Botte, 1995; Dai & Buus, 1991; Dai et al., 1991; Green & McKeown, 2001; Greenberg & Larkin, 1968; Moore et al., 1996; Scharf et al., 1987; Tan et al., 2008) using a naive online sample of participants who utilized variable consumer equipment in uncontrolled home environments. This effect was notably robust even at the individual participant level: 56 of the 60 participants (93.33%) showed at least a 5% detection advantage for signal versus probe frequencies. Moreover, the effect of the high-probability signal was established rapidly among naïve listeners. This supports models of frequency-selective attention dependent upon a system that adjusts very rapidly to input regularities (Fritz, Shamma, Elhilali, & Klein, 2003; Hafter, Schlauch, & Tang, 1993).

### Time course of the probe-signal effect

Here we asked how the probe-signal effect may change as participants become more practiced over time. As in the literature, we calculate the probe-signal effect as the difference between accuracy for the most probable frequency (the ‘signal’) and average accuracy for the least probable frequencies (the ‘probes’ in Figure 11A). A linear mixed-effect model (LMM) using block index as a fixed effect and participants as a random effect showed that the probe-signal effect diminished slightly as the task progressed (F(1,718) = 7.87, *p* = .0052). This result was mirrored in RTs (Figure 11B); although response times to both signal and probes decreased over time, the difference between the two was overall smaller at the end of the experiment (LMM, effect of block index: F(1,711) = 10.75, *p* = .0011).

**Figure 11.**
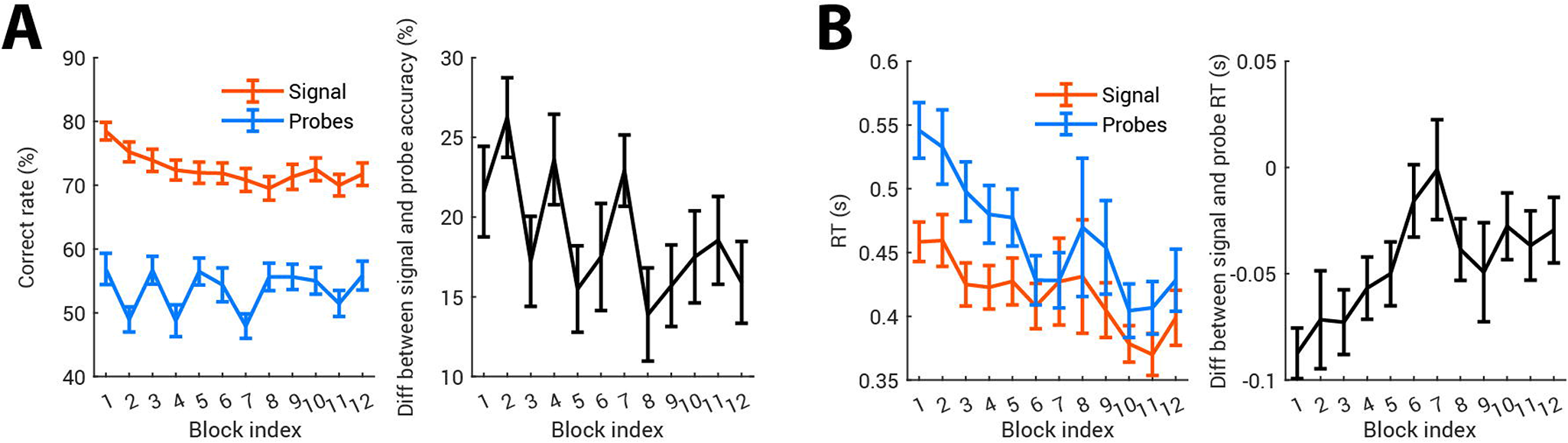
Dynamics of the probe-signal effect. In all plots, the error bar shows ±1 SEM. (A) Probe-signal effect in accuracy decreased over time. In the left panel, signal and probe accuracy are computed for each block and averaged across participants. The probe-signal effect is computed as Signal Accuracy - Average Probe Accuracy; the group average probe-signal effect is plotted in black in the right panel. (B) The probe-signal effect in RT is shown in the same manner; RTs to signal and probe tones are plotted against the block index, with their difference shown in the right panel. Note that since the probe-signal effect in RT is computed as RT-to-signal minus RT-to-probe, more negative values mean larger probe-signal effects.

### Effect and time course of motivation variables across the probe-signal experiment

During the 12-block probe-signal task, participants were instructed to rate how well they felt they performed, how motivated they were, and how tired they felt at the end of each block. This allowed us to examine how the probe-signal effect evolves along with individuals’ dynamics of confidence, fatigue and motivation.

As would be expected given the difficulty of the probe signal task, confidence remained low throughout (Figure 12A). An LMM on confidence rating showed that as the task progressed, confidence decreased slightly, but not significantly so (F(1,595) = 3.50, *p* = .062), with higher confidence associated with better overall accuracy (F(1,595) = 5.21, *p* = .023). With increasing time on task, fatigue accumulated (Fig 12B, LMM on fatigue with block and accuracy, effect of block: F(1,595) = 47.01, *p* < 10^-10^) and motivation diminished (Fig 12C, LMM on motivation with block and accuracy, effect of block: F(1,595)=59.36, p < 10^-13^). However, ratings of fatigue and motivation were not significantly related to the probe signal performance of that block (LMM with block and accuracy, effect of overall accuracy on fatigue: F(1,595) = 0.32, p = 0.57; effect of overall accuracy on motivation: F(1,595) = 1.06, p = 0.30).

**Figure 12.**
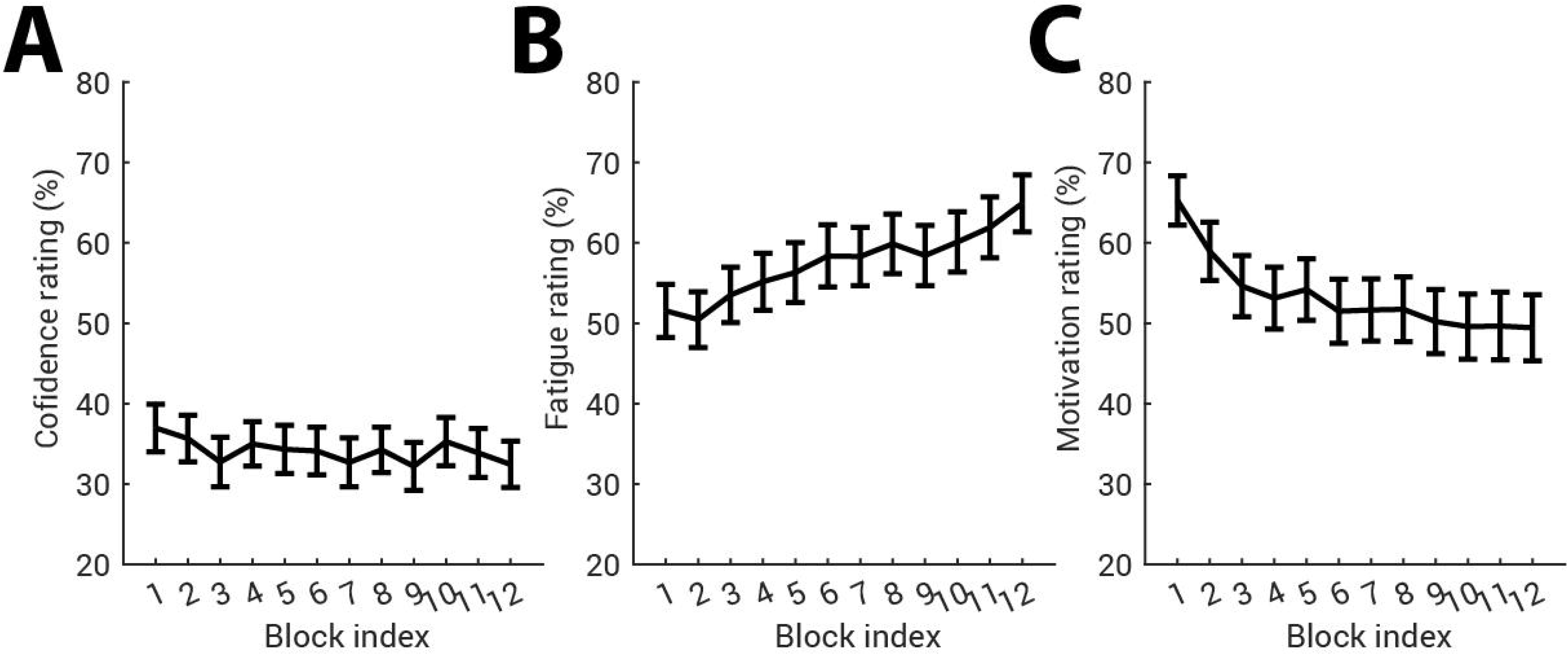
Dynamics of confidence, fatigue, and motivation across the probe signal experiment. In all plots, the error bar shows ±1 SEM.

Finally, to investigate the effect of motivation and fatigue on the probe-signal accuracy effect, we ran an LMM with block index, motivation rating and fatigue rating as fixed effects and participants as a random effect^3^. While fatigue did not show an influence on the probe-signal effect (F(1,594) = 0.067, *p* = .80), the probe-signal effect decreased over blocks (F(1,594) = 5.54, *p* =.019) and increased slightly with motivation (F(1,594) = 5.61, *p* = .018). This suggests that a larger probe-signal effect is predicted by high motivation, but not low fatigue.

## Discussion

Here, we developed and tested new approaches to making auditory psychophysical methods viable for online studies with psychophysics-naive participants. We first showed that the problem of limiting the range of stimulus sound levels can be addressed by using each participant as their own reference for setting stimulus levels at a given dB RMS above their noise detection threshold. We then showed that online participants’ perceptual tone-in-noise thresholds could be reliably estimated, not only by combining data from multiple tracks as is classically done, but also with a single short staircase track with a simple mode-based analysis that is easily implemented even by novice researchers. Individual differences in online participants’ apathy, confidence, and motivation did not significantly influence their perceptual thresholds, although those who were more fatigued tended to show somewhat less-sensitive thresholds. Online tone-in-noise thresholds also were reasonably reliable in setting the desired accuracy level for a new online version of the classic probe-signal task (Dai & Buus, 1991; Dai et al., 1991; Greenberg & Larkin, 1968; Moore et al., 1996; Scharf et al., 1987). Moreover, despite using only a third of the trials of a recent and efficient in-lab version (Anandan et al., 2021), we found a robust frequency-selective auditory attention effect overall, and in 93% of individual participants. This compares well with results from studies with few participants each undergoing thousands of trials. Indeed, the probe signal effect itself could be clearly detected at a group level from the first block of trials (Figure 11A). The magnitude of the attentional probe-signal effect decreased somewhat as the task proceeded, which was related somewhat to a decrease in motivation over time, but was not significantly associated with overall participant fatigue, or with changes in fatigue ratings over time. In sum, these experiments show that using such vetted ‘auditory hygiene’ measures can facilitate effective, efficient, and rigorous online auditory psychophysics.

### A method for remotely setting stimulus amplitude levels

The human auditory system is capable of successful sensing signals across a remarkable range of acoustic intensity levels, and many perceptual and cognitive phenomena are robust to level changes (Moore, 2013). However, a lack of control over auditory presentation levels - as is often the case in online experiments - is far from desirable on several grounds. Hearing safety is of course a potential concern for online experiments, particularly when presenting punctate sounds for which onset times are considerably faster than the ear’s mechanical protective mechanisms can respond. Sounds presented at different absolute levels evoke responses in distinct auditory nerve fibers, which can be selectively affected by pathological processes (Schaette, 2014; Verhulst, Altoè, & Vasilkov, 2018). As noted above, the frequency selectivity of subcortical and cortical auditory neurons can vary systematically as a function of sound pressure level (Moore, 2013; Schreiner et al., 2000). Of course, absolute sound pressure level is not the only factor to consider: individual participants with normal hearing will show thresholds with a range of up to 30 dB HL, and therefore a fixed absolute amplitude level can result in quite different perceptual experiences for participants who lie at one end or the other of this hearing range.

In Expts 1a-d, we found that community-recruited participants could very quickly adjust levels via the computer volume setting to estimate their hearing threshold using diotic pulsed white noise. The 20-25 dBA SPL range accords well with that of normal hearing (Park et al., 2016); and the extrapolated threshold levels are highly consistent across the outdoor settings of the three experiments. Expt 1c and Expt 1d showed that participants’ thresholds indoors and outdoors were highly correlated; this not only shows excellent reliability (albeit in a relatively small sample given the strictures of working during the COVID pandemic), but also demonstrates the robustness of this method to different acoustic environments. The spectra of both background noises have generally low-pass characteristics, so headphone attenuation should not be appreciably different; thus, we measured attenuation in only one of the backgrounds (the anechoic chamber). The fact that the thresholds were quite similar in indoor and outdoor environments, despite the large difference in ambient noise levels, may be due to the non-stationary nature of the noise, providing gaps in which the listeners could detect the presence of the signal. We plan a larger-N follow-up when in-person studies in indoor environments are more feasible than at the time of writing.

For experimenters who need to present auditory stimuli within a given range of intensities or at a particular level above perceptual threshold, the presentation level can be referenced to the RMS level of the white noise stimulus used in the amplitude setting procedure. For instance, say an experimenter wants to set her stimulus presentation level at ∼ 60 dBA SPL. an average of. If she assumes the typical participant will be in an acoustic environment similar to the outdoor setting (with an average 50 dBA SPL ambient noise level) the average stimulus soundfile *RMS* to produce an average 60 dBA SPL level in the headphones can be estimated. Recall that the RMS of the white noise file used in Experiment 1a (background noise level ∼50dBA) was 0.000399; using this stimulus, participants set their thresholds to an average of 29.4 dBA SPL (range 22.3 – 35.6 dBA SPL).

To achieve the desired average SPL of 60 dBA for the experimental stimulus, the experimenter can scale the RMS amplitude of the experimental stimulus soundfile as follows. First, calculate the difference in dbA between the desired average SPL and the SPL associated with the average participant’s threshold for white noise: 60dBA - 29.4dBA = 31.6dB SPL. Second, calculate the RMS of the experimental stimulus; for the present example, we will assume the sound has an RMS of 0.0080. Third, calculate the RMS amplitude difference in dB between the experimental stimulus (0.0080) and the white noise stimulus used for thresholding (0.000399), using the following formula: dB ratio = 20 * log_10_(*experimental stimulus RMS* / *white noise RMS*) = 20 * log_10_ (*0.0080/0.000399) = 26.04dB.* Fourth, calculate the difference in dB between the results of step (1) and step (3), e.g., 30.6dB minus 26.04dB = 4.54dB. Finally, scale the experimental stimulus file amplitude by this amount to achieve the desired RMS, either in an audio editing program like Audacity, or through calculation on the soundfile values itself in a program like Matlab, e.g., output_stimulus = input_stimulus * 10 ^ (4.54/20).

Assuming our Expt 1a-c noise detection threshold results generalize to the online population, the 10^th^ and 90^th^ percentiles of presented levels across all participants should be approximately 54- and 68-dBA SPL. Alternatively, the stimulus RMS could simply be scaled 30 dB above each individual participant’s white noise threshold level to ensure that stimuli are sufficiently audible to the vast majority of participants. One very important caveat to this approach is in the case where the spectrum of experimental stimuli is far from the 1-4kHz band that will drive much of the detectability of the white noise stimulus (for instance, pure tone stimuli at lower or very high frequencies). Here, it is important that either additional checks be placed on stimulus amplitude, or that a different thresholding stimulus be used (for instance a narrower-band noise centered around the stimulus frequency).

Of course, individuals will have different laptops with different sound card characteristics, different quality headphones etc. Although we chose to use a band-limited noise as our stimulus to help mitigate these potential confounds, this does not ensure that there are no differences across subjects. Within the selected band, however, the frequency response of each participant’s setup will be constant between the amplitude setting procedure and the psychophysical test of interest, which renders across-subject differences in technology less critical, especially when common-sense steps are taken in designing each online experiment.

For example, avoiding both narrow-band stimuli like tones as well as stimuli that are not band-limited like broadband noise will limit the effects of across-subject hardware frequency response differences on results. Ensuring that subjects are working at SPLs that are reasonably above threshold will help ensure that audibility is not a confound. Better still would be to design studies in which the experimental SNR ensures that stimulus noise levels are likely to overwhelm the levels of environmental noise sources.

Asking participants to avoid using open-back headphones, and instead to use closed-back or insert phones with soft rubber or latex tips will likely help alleviate the intrusion of environmental noise on psychophysical data. To establish the potential amount of insertion loss that might be expected from closed back headphones like those used here, we placed an acoustic manikin (Knowles Electronics Manikin for Acoustic Research) in the anechoic chamber, and presented the band-limited white noise stimulus from approximately 2m away and directly in front. Recordings were made from KEMAR’s microphones with and without the Beyer Dynamics DT150 headphones used in the study, in position. We then compared the RMSv levels of each recording and found that the headphones provided about 9 dB of attenuation. We re-ran this analysis with various other headphone models that were readily available to us (as well as 3M foam ear plugs as a reference) to determine the degree of variability. These data are shown in Supplementary Materials Figure S3. Among the circumaural phones we tested, the AKG K271’s provided the least amount of attenuation, at about 6 dB, while the Beyer Dynamics used in the study provided about 9 dB of attenuation.. The two sets of supra-aural phones we tested – RadioEar DD45 and TDH-49 – provided the poorest attenuation, along with the Apple AirPods, which is not surprising given their non-pliable hard plastic shell. While far from exhaustive, this analysis suggests that even inexpensive circumaural closed-back headphones, will likely provide at least 6 dB or so of attenuation.

### Time-efficient and reliable estimation of tone-in-noise thresholds

We used the ‘amplitude-setting’ method of Expt 1 with all Expt 2 participants. Based on this, the online continuous white noise masker played during both parts of Expt 2 was set to 40 dB above each participant’s white noise detection threshold, resulting in an average of 66 dBA SPL (SD = 4.3). Using a standard staircase technique to estimate tone-in-noise thresholds, we were able to obtain stable threshold estimates in online participants (Figure 6), not only by using the traditional method of averaging the means of the last six reversals from three staircase threshold tracks, but also using an easy-to-calculate and robust mode of the SNR levels from the first 20, 30, or all 40 trials (Figure 7). We also found that it was possible to obtain a reliable threshold from a single track of 20 trials (Figure 8), entailing about a minute of online testing.

If a psychophysical task takes about 3 sec per trial, then a standard thresholding track of 40 trials would take two minutes, and three tracks would take 6 minutes excluding time between tracks. Using the same assumption, the mode-of-20-trials approach would take about one minute to generate a threshold, a significant reduction in testing time. This streamlined threshold setting approach may be very attractive for online testing settings, as the vigilance of participants might not be as high as it would be during in-person testing, where experienced participants can typically be expected to generate reliable data for 1.5 hours or more. This fact places a premium on time-to-threshold for online studies. However, further investigation is needed into estimation with multiple modes, with multiple tracks, or more disperse SNRs. Although in more traditional psychophysical testing scenarios, this reduced thresholding time would not be worth the corresponding increased variability associated with the mode-based approaches described here, online testing easily offers larger sample size from a more diverse population than traditional in-person testing on the university campus. Thus, it is suggested that the streamlined thresholding approach described here, along with shortened online testing sessions, and increased sample sizes can yield better, more reliable outcomes when testing online.

There are other issues to consider in maximizing the efficacy of online testing using streamlined thresholding. For example, psychophysical tasks that require participants to work near or at their thresholds-in-quiet are likely not suitable, because overall ambient sound level is less controlled in online studies (as described in Expt 1), which raises signal audibility as an issue. This adds more uncertainty by the participant to the task, which makes short, 20-trial tracks less reliable. In a similar way, tasks in which the required perceptual decision is based on subtle cue differences like those that are often categorized as timbral may not be good choices for online study, again because bad tracks are more likely. Generally, it is recommended to choose psychophysical tasks that are easy for novice listeners to understand and ‘hear out,’ and to implement a training regimen that is carefully designed to clarify the perceptual task for listeners to avoid bad tracks, which are more difficult to discern with streamlined threshold setting procedures.

Finally, another potential advantage of the mode-based approaches might lie in their ease of computation. It is undeniable that online platforms such as Pavlovia.org and Gorilla.sc make psychophysical testing accessible to many, including students and other non-experts. These novice psychophysicists may have valid and interesting scientific questions. However, they may not have algorithms at-the-ready to estimate thresholds from staircase reversals using traditional approaches, a limitation that should never be a barrier to entry into the field.

### Rapid and robust online auditory psychophysics of frequency-selective auditory attention in single participants

The probe-signal paradigm (Borra et al., 2013; Dai & Buus, 1991; Dai et al., 1991; Green & McKeown, 2007, 2001; Greenberg, Bray, & Beasley, 1970; Greenberg & Larkin, 1968; Macmillan & Schwartz, 1975; Moore et al., 1996) would not seem to be a promising target for online research. Classic and more recent psychophysical studies have both recruited highly experienced participants for multi-day experiments with extensive tone-in-threshold measurement, multiple practice sessions, and thousands to tens-of-thousands of trials in the primary experiment, all conducted with specialized equipment in acoustically isolated laboratory settings (Borra et al., 2013; Dai & Buus, 1991; Dai et al., 1991; Green & McKeown, 2007, 2001; Greenberg et al., 1970; Greenberg & Larkin, 1968; Howard, O’Toole, Parasuraman, & Bennett, 1984; Macmillan & Schwartz, 1975; Mondor & Bregman, 1994; Moore et al., 1996; Tan et al., 2008; Wright & Dai, 1994). Here, Expt 3 violated each of these experimental *desiderata* in a single, brief online session with psychophysically naïve participants using their own computers and headphones in uncontrolled home environments. Nonetheless, we observed a probe-signal effect in most participants, with a signal-to-probe accuracy advantage of about 20-25%, on par with the magnitude of frequency-selective attention observed in studies with tens of thousands of trials (Dai et al., 1991; Greenberg et al., 1970; Greenberg & Larkin, 1968; Macmillan & Schwartz, 1975). Despite the relatively uncontrolled online experimental setting, the probe-signal effect was apparent even in response time; participants were faster in noise at detecting the signal, as compared to probe tones.

Beyond the convenience of recruiting participants online, there is power in demonstrating psychophysical effects like the probe-signal effect in a diverse sample of psychophysically naïve participants. Rather than rely on highly expert listeners, or even naïve listeners sampled from the relative homogeneity of a university campus, Expts 2 & 3 involved a world-wide sample. Behavioral science is increasingly recognizing that human behavior sampled for convenience only across university populations may be WEIRD (Western, Educated, Industrialized, Rich and Democratic; Henrich et al., 2010), and therefore not necessarily representative of populations at large. Although there are sound reasons to expect many psychophysical paradigms to generalize beyond WEIRD samples, this assumption has not often been tested (but see McDermott et al., 2016). The present results demonstrate that, with the right approach, it is indeed feasible to successfully conduct even challenging psychophysical paradigms dependent on thresholds online, and among inexpert participants. This substantially broadens the reach of psychophysics and opens the door to the possibility of large-scale psychophysics. Here, even with modest sample sizes (that nonetheless exceed typical probe-signal samples by an order of magnitude) Expt 3 demonstrated that it is possible to observe the evolution of frequency-selective attention via the probe signal effect from the first block onward, in both accuracy and RTs.

### Motivation in online participants

Another concern with online experimentation is participants’ motivation; low levels may result in high drop-out rates and poor task engagement and performance, in turn affecting the validity of the experimental results (Shen & Chun, 2011). Compared with online participants, those attending in person might be expected to be more motivated since they have already made the effort to visit the lab, and social evaluative stress caused by the presence of the experimenter can motivate them to some degree (Bianco et al., 2021), as in the long-documented Hawthorne effect (McCarney et al., 2007).

Meanwhile, online experiments are normally completely anonymous and without supervision, leading to a common worry that the online population might be more apathetic than in-lab participants. Because of these concerns, we expected that the estimated thresholds might, at least in part, reflect motivation level. However, the estimated thresholds in Expt 2 showed no relation with motivation, neither as expressed by the apathy index (a personality-trait-like component of motivation derived from a well-established apathy questionnaire, Ang et al., 2017), nor the motivation ratings before and after Expt 2. Similarly, in Expt 3, the self-reported questionnaire-derived apathy index, as well as its subdomains, could not explain the strong probe-signal effects observed. However, we did find a weak but significant effect of in-experiment motivation on the probe-signal effect: blocks in which listeners were more motivated generated a larger probe-signal effect. Interestingly, we also found that in motivated people, high confidence strongly prevented motivation loss over time, while in apathetic people this protective effect was diminished.

One might worry that the online threshold estimation may be affected by the on-task motivation of the participants. It is interesting that, at least that in this study, we did not observe any influence of self-reported motivation on the threshold estimation amongst the remotely tested participants. On the other hand, motivation showed a small influence on the probe signal effect. This is in line with the previous work (Watson & Clopton, 1969) which found motivation — regulated by applying electric shock on incorrect trials— increased sensitivity in a simple tone-in-noise detection task but the increase was rather small. One explanation for the absent effect of motivation on the threshold estimation here is that the effect of motivation on performance is sensitive to the length of the experiment; the probe signal experiment was longer (around 20 minutes) and was run after the threshold estimation (a length of around 10 minutes). This, with no observed effect of motivation in Expt 2, indirectly suggests an advantage of keeping experiment time shorter. In summary, any generalizations of the motivation-related findings here should be taken carefully.

## Acknowledgements

We thank Christi Gomez, Erin Smith, and Sydney Sepkovic for their assistance in collecting in-person data. We thank Dr. Alessandro Rinaldo, Carnegie Mellon University Department of Statistics and Data Science, for statistical consultation.

## Data Availability

Test implementations of the amplitude setting (Expt 1) are available in JavaScript [https://gitlab.pavlovia.org/sijiazhao/amplitudechecking_demo] and Gorilla [https://gorilla.sc/openmaterials/261557]. A demo can be found on Pavlovia [https://run.pavlovia.org/sijiazhao/volumechecking_demo].

The guide to implement the staircase procedure (Expt 2) is available at [https://sijiazhao.github.io/how-to-staircase/]. A demo is available on Pavlovia [https://run.pavlovia.org/sijiazhao/threshold_demo/] with publicly available in-house code [https://gitlab.pavlovia.org/sijiazhao/threshold_demo].

The probe-signal task (Expt 3) can be tried at [https://run.pavlovia.org/sijiazhao/probesignal_demo] and its code can be found at [https://gitlab.pavlovia.org/sijiazhao/probesignal_demo].

The raw data of this study are available on GitHub [https://github.com/sijiazhao/TPS_data].

## Author Contributions

Sijia Zhao: Conceptualization, Methodology, Software, Validation, Data Curation, Writing Chris Brown: Conceptualization, Methodology, Validation, Writing, Funding Acquisition Frederic Dick: Conceptualization, Methodology, Validation, Writing, Funding Acquisition Lori L. Holt: Conceptualization, Methodology, Validation, Writing, Funding Acquisition

This manuscript has been accepted by Trends in Hearing on 21 July 2022. We want to thank Dr. Andrew Oxenham (Editor in Chief), Dr. Michael Stone (Action Editor), and the three reviewers for taking the time to make such helpful comments. Our response letters to the three revision rounds are attached below.

## Authors’ Response Letter – Revision Round 1

Thank you for the opportunity to submit a revised version of our manuscript. Below we reply to all Action Editor (AE) or Reviewer (R) comments, *italicized* and labelled as AE or R [Reviewer number]-[comment number], with Author Responses labelled AR[Reviewer number]-[comment number]. We begin with your editorial note.

### Comments from Editor Dr. Andrew Oxenham

*As you will see, both reviewers and the Associate Editor, Michael Stone, find the topic to be important and timely. However, all three have a number of major criticisms that would need to be thoroughly addressed before the manuscript could be considered further for publication. In my reading, it would require a very major rethinking and restructuring (as well as retitling, as well as potentially new data collection, given the very low levels used and consequent small number of bits exploited, as described by the Associate Editor). However, I am leaving the door open for you to submit a revised version in case you feel able to address the comments below*.

AR-E: We extend our thanks to you, the Associate Editor, and the reviewers for the ample suggestions and critiques. As we describe below, we address concerns through new behavioral experiments, acoustic and stimulus measurements, analyses, and extensive revision of the manuscript itself (including the title, as requested).

### Associate Editor (AE) Michael Stone comments, and Author Responses (AR-AE)

*AE-1.0: This paper reports two sets of experiments that investigate methods for improving reliability of thresholds obtained in psychophysical tasks performed online. The use of online, rather than laboratory-based, tests results in reduced ability to use quality equipment, consistent methodology and reliable calibration*.

*As such it is a timely contribution to the literature, but makes little or no mention to much larger recent and current discussion of the issues in the community, such as the ASA Task force report eg* https://asa.scitation.org/doi/10.1121/2.0001409 *although that does conclude “Longer-term goals include identifying best practices and providing resources for evaluating outcomes of remote testing”, ie something which this paper could contribute to*.

AR-AE-1.0: Thank you for bringing the very relevant report to our attention; we were aware of the task force via the Auditory mailing list but had not seen the proceedings paper when it was published in late April 2021. We have incorporated this into the introduction and discussion and appreciate you pointing it out. We have also submitted the online sound level setting (https://www.spatialhearing.org/remotetesting/Examples/ExampleLevel) and probe signal paradigm (https://www.spatialhearing.org/remotetesting/Examples/ExamplePitch) to the experimental methods repository associated with the task force.

*AE-1.1: It also makes no mention of alternative stopping criteria in psychophysical tracks, such as that used by Dillon et al. 2016*.

AR-AE-1.1: Thank you for pointing out this paper on speech-in-noise thresholds via telephone, we were unfamiliar with it. More generally, we now more explicitly mention other approaches to psychophysical thresholding on pages 25-26 and the new Figure 6C.

*AE-2: The technical descriptions do not give enough detail. Why was an 8-kHz bandwidth chosen ?. What does this mean for estimation of thresholds, especially if there is an underlying hearing lossWhy white noiseThis biases detection to mid/high frequencies, as well as being environmentally unrealistic, and also confusable with the soundcard electrical noise. What about an inharmonic tone series, possibly with a uniformly exciting spectrum ?*

AR-AE-2: We should have made it clear why we chose white noise and those bandwidths. We now explicitly state our rationale to limit bandwidth between 80 Hz-8000 Hz, based on exactly the issue the associate editor raises: the high-frequency bias of white noise, as well as the presence of low-frequency line noise. Nonetheless, we take your later point regarding the potential bias from white noise given ambient spectrum, and have re-run the level setting experiment (quiet environment only) with both white and pink noise (within subjects design), reporting the data here in the response to reviewers as well as in the paper itself. The results show that there is very good agreement (r=0.84 over volume settings, see page 17-19 and new Figure 4A), with the pink-noise stimulus yielding volume settings (see below for terminology) that are on average one Mac volume setting higher, ∼2dB SPL (see figure and below).

*AE-3: p2 line 7 “auditory hygiene” An unfamiliar term with no prior definition. Not really useful here without such, so possibly best left for later*.

AR-AE-3: We agree, and have operationally defined the term here (page 4).

*AE-4: p2 line 17 “volume setting” : “volume” is not in the lexicon for auditory psychophysics*.

AR-AE-4: While this is certainly true, it is the term that essentially all sound device and computer manufacturers use for their controls and what participants understand (as opposed to the more precise intensity or amplitude adjustment). We deliberately chose to use ‘volume setting’ only when referring to the arbitrary sound amplitude changes that results from manipulating the volume setting buttons on the laptop (while the steps are fairly linear, the step size is arbitrarily chosen by the manufacturer). We think it is important to use a term to refer to the ordinal data of volume setting steps, and differentiate it from the interval data associated with amplitude and intensity. We have gone the manuscript to make sure that this is consistent throughout.

*AE-5: P4 line 30 “segment of broadband Gaussian white noise generated to have a particular root-mean-square (RMS) amplitude.” Only later are we told that this noise is, in fact, band limited to 8,000 Hz. This makes a large difference in the understanding of self-calibration. The RMS is referenced to the “broad band” noise, yet for a “white spectrum, the bulk of the power is in the top octave. Therefore this calibration is largely set by the top octave, or rather, the audibility of the top octave through the headphone & soundcard combination. Since most headphones are pretty imbalanced above 4 kHz, partly due to ear canal loading, as well as the design philosophy (hype) from the manufacturer, then the mentioning of potential error sources in calibration is completely overlooked at this stage*.

AR-AE-5: We addressed these issues in a number of ways. First, we compared pink and white noise thresholds for the same participants, showing that they are highly correlated (near the limits of test-retest reliability, see new Figure 4A) but do show higher thresholds for pink noise of ∼3dB SPL. Second, as described below, we also re-ran the experiment (within subjects) with both the Beyer Dynamic headphones and a $20 pair of Sennheiser HD206 circumnaurals, which showed extremely good agreement (see new Figure 4B). Finally, we now use the same stimulus for calibration as we did for testing.

*AE-6: P6 line 10 mentions A-weighting, yet the weighting scale is dropped in the remaining numbers on line 11-13*.

AR-AE-6: A-scale weighting was used throughout and is now labelled explicitly in all instances.

*AE-7: p6 line20 : UK-obtained ethics (with no reference number, also on p13 line 4) being used to support a study on US soil (ExptS 1a, b & c) legally unlikely without clarification*.

AR-AE-7: Approval was obtained in the UK for the study to be conducted online, with the express understanding that anyone could participate in any country, as is now typical for online studies conducted across multiple platforms. Clarification has been added (page 9).

*AE-8: P6 line 32 : RMS of 0.000399. ie −68 dB. So the signal range peaks at about −58 dB. The coding barely requires a 6-bit signal in a 16-bit format. For many cheaper soundcards, the broadband SNR is not much lower than about −70dB RMS. This signal is awfully close to the noise floor of many domestic appliances…. As figure 1 shows (“noise, extrapolated”)*.

AR-AE-8: Please see *AR-AE-11* for discussion and data on SNR.

*AE-9: P6 line 59-60 : Mac volume levels : are these as well thought our as Windows levels which are a linear scale (rendering levels above “70” barely detectable in difference)Figure 1 shows this to be nearly the case with a step variation of between 2 & 8 dB, and confirmed on p7*.

AR-AE-9: As we lay out below, we re-ran the calibration with bandpassed white noise, and the generated functions are quite linear; an inspection of the figure shows that the majority of non-linearities appear to result from the non-linear % volume increments (which are arbitrary values).

*AE-10: P 7 line 9-16 . Too much detail here: the headphones are closely coupled to the artificial ear, so precise details of “anechoic” are unnecessary since the Beyer headphones are circumaural and closed back, so have some low-frequency attenuation of external sounds. The important issue is the NR of the room. Which then gets mentioned in the very next paragraph, but next section. 28 dB SPL : flat weighting, C or AThis needs re-ordering*.

AR-AE-10: These changes to the manuscript are now made.

*AE-11: There is then a potential flaw here. Because the ambient noise floor was ca NR20, the authors extrapolate the effect of their inferred volume setting on the white noise SPL produced in the coupler. This is predicated on the soundcard producing electrical noise at its output << 3.5 dB SPL broadband, ie much lower than the replay level of the white noise at the lowest volume setting. This is highly unlikely. Figure 1 says a 0.5 V signal produces near 100 dB SPL, and we are not told if this is full scale sine. We are then asked to believe that the output SNR of the sound card < −(100-3.5) ie < −96 dB. This is only really quoted/achieved on the aftermarket 24 bit soundcards*.

AR-AE-11: In response to this criticism, we re-ran the SPL analysis using the 80-8000 Hz band-passed white noise stimulus used for testing, rather than using pure tones. As mentioned above, we averaged 3 sets of measurements over three different days to maximize stability. We measured dB SPL from the headphone using the noise stimulus file adjusted to three amplitude levels: at the RMS level used for data collection, as well as that stimulus increased in level by +10 and +20 dB. Figure 1 has been updated to show these results. As might be expected, the gain functions are more linear than they were for pure-tone stimuli, particularly at low volume settings. We also found average overall levels that were a few dBA higher than we measured in the first submission. The net result of these two differences is that the SPL at the lowest Mac volume setting is now about 16 dBA SPL with white noise, as opposed to about 3.5 dBA SPL is originally reported. These new data are much more in line with our experience, which is that when we listen to the stimulus at the lowest volume setting in a controlled environment, it is just audible to us and sounds like white noise. We thank the Associate Editor for prompting us to be more thorough in carrying out and reporting these measurements.

All figures and statistics related to Experiment 1 have been updated. See pages 10 to 18. Regarding soundcard SNR, please see AR-AE-17 for response and new data.

*AE-12: Figure 1 caption does not describe how the 1-kHz curves were derived, and what the scaling between their electrical and digital levels was. Therefore we are further in the dark as to the likely electrical SNR of the sound card*.

AR-AE-12: This is now described in Figure 1; as noted above, sound card SNR is addressed below in AR-AE-17.

*AE-13: P10 lines 41-43 : 28 dB SPL vs 57 dB SPL. This comparison is fairly meaningless unless one knows the relative spectra of these two noises, an can compare to the ear-coupled spectrum of the “white noise”. Since the thresholds will largely be driven by the 2-4 kHz region, and the power of the white noise increases by +3 dB/oct, while the ventilation/park noise will be decreasing by typically between −3 to 9 dB/oct, then the SNR in the 2-4 kHz region will be +12 to +18 dB relative to that around ventilation noise peak of 500 Hz. Obtaining noise thresholds of <, 30 dB SPL is therefore not unexpected*.

AR-AE-13: We have now included a spectral analysis of the respective noise floors as Supplemental Figure 1 and we have updated the discussion of this issue as a result (page 39).

*AE-14: P10 line 54 “s/he” : English has slightly neater way by use of the gender-nonspecific “they”*.

AR-AE-14: We are pleased that the journal will accept the singular they (Foertsch & Gernsbacher, 1997) and have changed to the more modern usage.

*AE-15: Figure 3 caption : “with the black bins indicating” On my copy they were grey. P23 line 42, they are referred to as “grey lines”. Inconsistent*.

AR-AE-15: Figure 3B’s bin colour has now been changed to a lighter grey colour (RGB (128,128,128)) to avoid confusion between black and grey. The caption has been changed to “with the grey bins indicating…”. (Note that the “grey lines” in the original manuscript P23 line 42 (original Figure 8 caption and new Figure 10 caption) was an accurate description of original Figure 8B and new Figure 10B).

*AE-16: P11 line 41 (and elsewhere) : Moore 2013 : What page number in text bookWhy quote in units of SLHere the authors are talking about detection of a white noise, but I do not recall Moore talking specifically about that, especially in a clinical setting*.

AR-AE-16: Thank you for picking this up - we meant that normal hearing was defined with a 30dB range, but now are more precise about this, and also note the difference between thresholds based on pure tones and those on white noise (see page 16).

*AE-17: P11 line: I dispute that Expt 1 “validated” the procedure. The authors have not convinced the reader that the volume control was linear on the noise level due to the extrapolation procedure employed. It is highly likely that volume settings < 44 were likely contaminated by electrical noise in the output stages of the soundcard, ie AFTER level control had been applied. Inspection of Fig 1c suggests that it was impossible to obtain output levels below 15 dB SPL*.

AR-AE-17: We tackled these points in a number of new measures and analyses. To address both the question of output signal relative to line/audio card noise floor as well as generalizability across computers (*AE-18*), we recorded the experimental stimulus (white and pink noise), output at different volume settings from the headphone jack of both a new MacBook Pro and an old, entry-grade Asus Windows laptop. To avoid the additional electrical noise introduced by external amplification in our previous measurements, we routed the headphone jack output the line input of a 2011 MacBookPro. (The male-to-male cable was low quality and did pick up some electrical interference). For both MacBookPro and Asus laptops, stimuli were played on the Pavlovia online experimental platform as was experienced by participants; input was recorded and analyzed using Adobe Audition X and Matlab 2020b.

This is now reported in Supplemental Materials.

Figure S1 (see also below) shows the spectral power (in dB from peak amplitude) for white and pink noise for each computer around threshold volume settings observed for reported participants with the MacBookPro and for two observers with the Asus laptop. Power across stimulated frequencies is consistently above noise floor for all thresholds (+ ∼14dB for MacBookPro volume setting 44%), and floor noise levels are consistent across volume settings. As would be expected, the Asus laptop output line noise level is generally ∼3-4 dB higher that of the MacBook Pro, with an additional ∼2-3 dB noise levels in frequencies ∼< 400Hz.

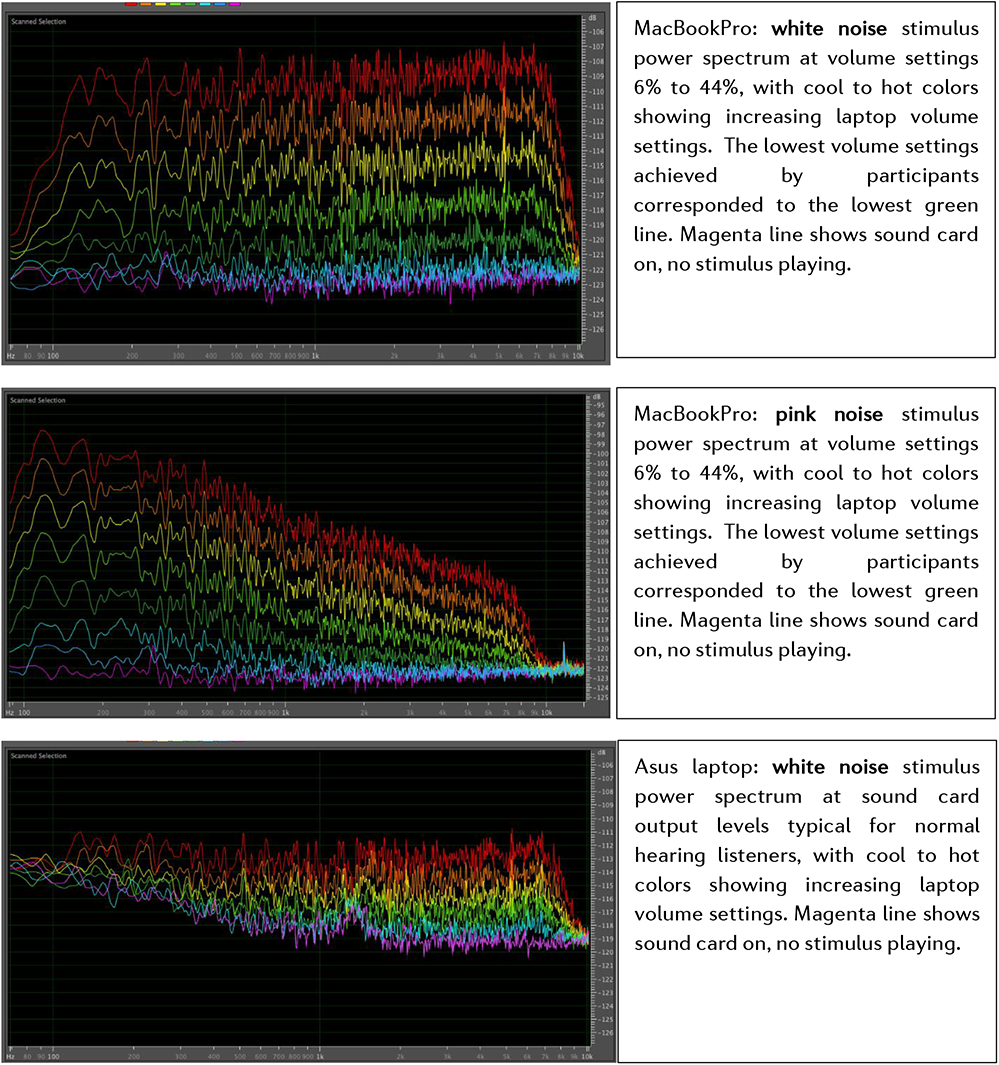
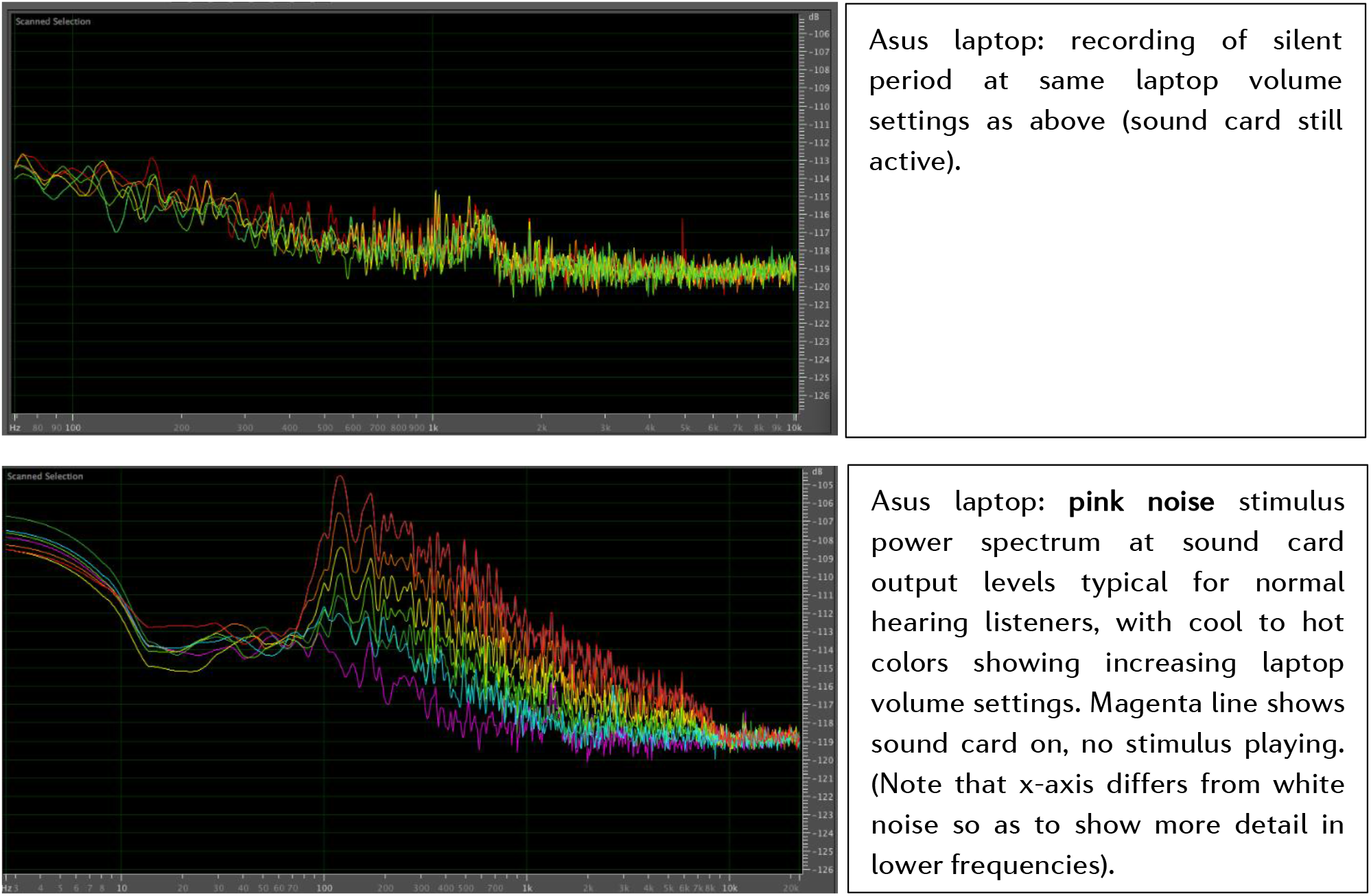

*AE-18: P11 line 58 : there is a leap of logic here. By use of one computer, with one set of headphones (with a particular response) the reader is asked to believe that the results can be extrapolated to any computer, with any headphone, without any reference to at least inter-headphones variability (as well as ability to exclude external sounds by being open-backed, or not*.

AR-AE-18: We agree with these points. In the original manuscript, as one estimate of susceptibility to background noise, we reported that participants’ white noise thresholds were highly correlated across outdoor and sound booth settings (r = 0.82, Figure 2C). In response to this comment, the revision adds an additional within-subject experiment (N=20) evaluating consistency of results over headphones. We find that white noise thresholds obtained with the Beyerdynamic headphones are strongly correlated (r = 0.65, Figure 4B) with those obtained using ($20) Sennheiser HD205 and popular Apple AirPods (r = 0.69, Figure 4C). (Note we do not recommend use of battery-powered Bluetooth earbuds for auditory experiments in the manuscript). The same participants’ white noise thresholds with Beyerdynamic headphones were also highly correlated (r=0.84, Figure 4A) with their pink noise thresholds using the same headphones (Page 17-19).

We have also added some discussion of the advantages of the closed-back design, along with a new figure comparing insertion loss across a number of different headphone models we had available to us (page 39-40).

*AE-19: P13 line 36 “thereby serving as their own sound level meter.” Here and used elsewhere . This is not really correct : really it is more a self-calibrated audiometer. (and by “self” I also include the effects of the headphones as well as any hearing variation)*

AR-AE-19: We agree, and have avoided use of the term ‘their own sound level meter’ throughout the manuscript. Thanks for the suggestion on the alternative term.

*AE-20: P14 line 16 “A long 300-sec white noise” Better as “A 300-sec duration white noise”*

AR-AE-20: Agreed and revised.

*AE-21: P14 line 43 : “with the continuous white noise background sound commencing in advance” Repetitious on previous*.

AR-AE-21: Thanks for flagging this, now fixed.

*AE-22: P15 line 55 “-13.8 dB SNR (i.e. the easiest level)” HuhBetween pages 14 & 15 we have “reducing to 0.75 dB once the level fell below 22.75 dB SNR” Some revision of signage is required around here*.

AR-AE-22: Thanks, now reads −22.75 dB SNR as it should have previously; we also clarify the way that we calculate SNR in this sentence (dB difference in RMS for sine wave signal and noise background).

*AE-23: P17 lines 42-46. This is a very long sentence*.

AR-AE-23: That it is; now split into two sentences.

*AE-24: P18 lines 16 onwards. Mode-derived thresholds. While the paper tries to claim novelty of approach, there is a distinct hole in the consideration of reversals & the criterion to stop. In an attempt to reduce trial time, Dillon et al (IJA 2016) used an adaptive stopping criterion, once the continuously updated standard error had dropped below a threshold. Ie it does not count reversals in the traditional way, but uses more information from the track than in the classic psychophysics approach*.

AR-AE-24: We now note the Dillon et al., 2016 paper as well as a few other thresholding optimization procedures. While trying not to lengthen the paper too much, in the preceding paragraphs, we also reference several papers by Peter Jones and colleagues which cover this topic in more detail.

*AE-25.1: P17 lines 45 onwards, P23 lines 51 onwards, P24 lines 38 onwards, p25 line 40 onwards. They are listed under “Results and discussion”. These are all something for the discussion section which only explicitly appears on page 27. One cannot really have two discussion sections*.

AR-AE-25.1: Here we followed the journal’s guidance on style, which itself references APA guidelines (APA journals often have experiment-wise discussion sections followed by a general discussion). However, as you and other reviewers found this structure awkward, we have moved discussion and interpretation into the formal discussion section.

*AE-25.2: P 25 line 42 : “As exciting as the results from psychophysical experiments can be,” This reads more like an opinion piece than a scientific report*.

AR-AE-25.2: We have made a number of stylistic changes to the manuscript in response to Reviewer 1’s suggestions and detail them below.

More generally, we appreciate the detailed technical suggestions and critiques. They definitely helped to strengthen the science and manuscript.

### Reviewer 1 comments (R1) and Author Response to R1 (AR1)

*R1-0.1: There are many reasons to evaluate and adopt procedures for remote psychophysical testing (for example, perception experiments conducted via online services). These include health and safety (a major current concern), along with diversity and equity of inclusion of research participants, access to more representative large-N samples or rare populations, and more. Yet many potential drawbacks of remote testing are anticipated, especially compared with traditional approaches that focus on rigorous control of stimuli, procedures, statistical approach, etc. Relaxing some of those ideas might be necessary in order to fully embrace remote psychophysics. The submitted manuscript advocates for exactly that: adopting less rigorous “auditory hygiene” in order to enable online experiments and reap some of the benefits mentioned above. It does so in three ways: first by persuasive argument against the rigid confines of “traditional” psychophysics; second by proposing “new” approaches that escape those confines (letting participants set their own levels, alternate averaging for psychophysical thresholds); and third, by demonstrating psychophysical results from online experiments that are in relative agreement with previous studies using “traditional” methods. The goal is laudable. The advantages of remote testing (mostly well-treated in the discussion) are clear. The proposed methods are reasonable. The data are reassuring*.

AR-R1-0.1: Thank you for the supportive comments.

*R1-0.2: But the manuscript itself undermines the work by attempting to move simultaneously in too many directions and making too many claims that–even if reasonable–are not specifically supported by the work. Part of the problem is that this paper itself lacks “traditional” focus. Neither a research report nor a systematic review, it runs light on novelty, scientific rigor, and thoroughness. Instead it combines a flock of brief experimental ideas that individually seek to enhance remote psychophysics in a different way. None of them is fully and rigorously evaluated in the context of relevant literature (for example, little acknowledgment is given to the wide range of rapid threshold approaches used in contemporary clinical tests, nor is the proposed solution directly compared to those procedures). Nevertheless, each element represents a thoughtful design decision that contributes positively to the success of the final, combined experiment*.

*Throughout my reading, I had the strong feeling that this would work quite well if it was presented as a study of motivation, confidence, and fatigue on probe-signal data collected via online methods. Each challenge and solution along the way (how to ensure correct SNR, how to estimate thresholds quickly, how to set the absolute level [in SL no less, possibly an advantage over traditional SPL-defined approaches]) could be presented as just that: a design problem and validated solution. In that type of presentation, the authors’ efforts to validate each choice would appear quite rigorous (How could we be sure our threshold averages were reliable enough? We compared various subsets of trial numbers to the averages we got from multiple tracks. How confident were we about the “just-audible” levels set subjectively by participants? We ran an initial experiment where we could compare to gold-standard in-lab measurements.)*

AR-R1-0.2: Indeed, the way that the reviewer lays out the paper is similar to how we had conceptualized the structure. We have revised the paper to make this much clearer to the reader.

*R1-0.3: But that is not how the paper is presented. Instead, it reads (rather confusingly, I think) as a set of three or four separate “micro-studies” (expt 1, expt 2a, etc) aimed at each question. Each attempts to stand on its own as a rigorous evaluation of a single problem, but none is complete enough in regard to the necessary comparisons, to the existing literature, etc. What would have seemed highly rigorous as an evaluation or validation of one step of the approach appears inadequate when presented as its own experiment. As a result, none of the elements of the manuscript are individually convincing, and the overall organization lacks cohesion*.

*Thus, my overall impression of the manuscript is not positive, even though I think the topic is very important, the experimental steps were valid, and that in a different presentation format it could be very strong. Below I give some more specific comments that I hope might help the authors improve the clarity of a future revision:*

AR-R1-0.3: We have taken the reviewer’s comments to heart and have revised the manuscript to improve coherence and integration with background literature. We hope that these presentational aspects no longer interfere with the experimental strength of the paper that the reviewer identified.

*R1-1: General and important*

*1) Parts, particularly the intro, are much too long and not clear (as in precise). There is a lot of jargon, slang, and approximate language throughout. I’m sure that some of these were deliberate stylistic choices, possibly meant to enhance the impact for non-expert readers. But for this journal, the language should be precise and to the point*.

*R1-1-1:a. The motivation of this work is framed almost entirely in terms of the strictures of testing during the COVID-19 pandemic. This may be easily understood by lay audiences, but there are many other reasons to consider and continue to pursue remote psychophysics*.

AR-R1-1-1:a. We agree and have added more general motivations for remote auditory psychophysics testing. See, for example, the modified Introduction starting from page 4.

*R1-1-1:b. There is a lot of very odd phrasing throughout which is quite off-putting. Some of these are clever linguistic inventions which convey something important (but maybe not as precisely as the authors imagine). Others seem intended to convey an emotional evaluation of the work, rather than allow the results to stand on their own:*

*i. “auditory hygiene” – this captures an important idea but is not precisely defined in the paper*

AR-R1-1-1b*-*i: As noted above, we now operationally define this term.

ii. “their own sound level meter” – not quite what an SLM does

AR-R1-1-1-b*-*ii: We have omitted this term, and instead refer to subjects serving as their own ‘self-calibrated audiometer’ as suggested by the Action Editor.

*iii. “abandon their deep traditions”*

*iv. “nigh unto impossible”*

*v. we “roll out” the volume setting paradigm*

*vi. “small tweaks to…procedures”*

*vii. “uncontrolled online testing” – not true, you’ve included many controls*

*viii. “quite a departure” from the high level of control*

*ix. “somewhat finicky paradigm”*

*x. “loosening of auditory hygiene”*

*xi. “potentially finicky”*

*xii. “somewhat persnickety”*

*xiii. “violated each of these experimental desiderata”*

*xiv. “ivory tower samples”*

AR-R1-1-1b*-*iii-xiv: We have gone through the paper and edited with an eye to the reviewer’s stylistic suggestions.

*R1-2: The impacts of Experiment 1 and 2a are undermined by their lack of novelty combined with little to no discussion of the relevant literature*.

*R1-2-a. Expt 1 proposes to define levels relative to participants on detection threshold. This is a rather common design decision (dB SL = dB above individual threshold), although generally based on objective measurements of detection threshold. The proposal is to let participants adjust the level of a known / standard test signal to a fixed level of detectability. In essence, this is the “biological check” employed daily to confirm (though not adjust) level calibration in most audiology clinics. Again, I think this was a reasonable design decision to support the broader project and I appreciate the authors’ careful validation of the measures obtained. It’s just not a novel result on its own to say that listeners can adjust level to just-audible and that sounds can then be presented at a known level relative to that setting*.

AR-R1-2-a: We now emphasize that our goal was to design and validate such an approach for unsupervised online listeners and have borrowed this ‘biological check’ audiology clinic metaphor, thanks for suggesting. See page 7.

*R1-2-b. Expt 2a proposes to compute “threshold” from the mode of trial values during a subset of the early part (of the asymptotic part*) of an adaptive track. The goal is to quickly obtain a threshold estimate. It’s a reasonable approach, but it’s not the first attempt to validate quick thresholds relative to “gold standard” psychophysical thresholds. Many clinical tests that use fixed stimulus sequences can estimate thresholds in just a handful of trials, for example, by simply counting the number of correct trials in a series that starts easy and gets harder. Again, I think this was a reasonable design decision in the context of the larger study, but evaluating the impact of this as its own result requires a lot more comparison to the actual state of the art (for quick thresholds, not just comparing to the slowest lab-based approaches)*.

*\*I will also note that the tracks benefitted from an “initial descent” (P14L57) that was strictly “one down” until the first incorrect response (i.e. the first reversal). The purpose is to quickly drive the track into the vicinity of threshold before beginning the fine (3 down / 1 up) track. Thus, regardless of the nature of the fine track, participants are likely to spend the next several trials in the vicinity of “threshold” which will be near to the first incorrect response during one-down descent. I suspect that a random walk of values from this point might give modal responses that are not terribly far off from those reported here*.

AR-R1-2-b: The Action Editor also noted that we did not sufficiently review the background for efficient threshold estimation; we agree and rectify this in the revised manuscript. We have also taken care not to imply a comprehensive comparison to current thresholding techniques as this is not the focus of the study, and other publications have carried out systematic comparisons of psychophysics methods.

*R1-3: I cannot figure out why Experiments 2a and 2b are labeled and described together. They address very different questions using different methods. I understand that their data were collected in the same sessions, but they are no more similar to each other than to Experiment 1. These should be labeled Experiment 2 and Experiment 3*.

AR-R1-3: As the reviewer suggested, we have restructured the manuscript so that these are separate experiments - we agree this is much clearer now, thank you.

*R1-4: The most important experiment is presumably the probe-signal experiment. Other experiments just represent careful consideration of design choices (level setting, SNR confirmation, threshold measurement) in support this experiment. It’s great that results look comparable to past work. BUT it’s also important to recognize that this experiment is limited by SNR, not absolute level. So it’s actually not that surprising that it worked. If you can ensure a regime where SNR for home-based participants is dominated by the noise presented experimentally, then other contributors (household background noise, etc.) will be minimized. Your confident assessment of the SNR should be all you need to know in order to predict that the probe signal experiment will work…This does not factor in the motivation, fatigue, etc. issues, which is why measuring those (and relating them to probe signal data) makes a lot of sense. But “it worked” is not really an interesting result*.

AR-R1-4: We did not make it clear enough that level is essentially always reported and controlled in probe-signal experiments, and that frequency sensitivity and discrimination are affected by absolute level. We also did not emphasize enough the potential importance of the motivation issues, or of the speed with which the probe signal effect can be measured online (e.g., it is evident within the first block of trials).

*R1-5.1: Throughout the discussion, there are repeated claims that the “adaptations” (subjective threshold setting, quick threshold measurement, etc) “facilitated” or “allowed” the experiments to work. There is no evidence to support these claims. That would require doing the study without those adaptations and failing to see the same results. What you showed instead is that “it worked” with the adaptations in place. Again, that does not imply that it worked BECAUSE of the adaptations, or that it wouldn’t have worked without them. Such claims need to be removed from the manuscript*.

AR-R1-5.1: Thank you for bringing this up; we agree entirely and have made sure that we do not claim that the experiments would not have worked without these adaptations.

*R1-5.2: Unfortunately, I think this undermines the whole manuscript. Because the presentation features all of these different adaptations, each evaluated on their own, the temptation is to force a conclusion that each is important even though that was not shown in the individual experiments. If the presentation was different, and focused on exper 2b and the motivation/fatigue analysis (where there are actually some very interesting results), then the adaptations make sense, as in “we had to deal with these problems and this is how we chose to do it.” It worked, so those choices must have been reasonable. Other choices might have worked, too, but we don’t know because that’s not what this is about. But that is not how the manuscript is presented, and the attempt to draw strong conclusions about the adaptations themselves is simply not correct*.

AR-R1-5.2: These are really useful suggestions for revision, and we have taken advantage of them for restructuring the paper’s argument and structure.

*R1-6: P29L29-35: This paragraph suggests that “non-expert” psychophysicists should be able to use simplified methods (rather than vetted approaches), so as to reduce a barrier to the field. That is a troubling claim. What I think it actually argues for is including correct / vetted procedures within the experimental tools available (i.e. the platforms), so that online can studies can be as rigorous as (and directly compared to) in-lab studies*.

AR-R1-6: Indeed, it is the latter point that we strove to make, not the former. We have clarified this in a couple of places in the manuscript, borrowing some of the reviewer’s phrasing.

*R1-7: Specific and important*

*R1-7-1: A number of terms and measures are used imprecisely or without definition. The following should be defined when used, in my view:*

*R1-7-1-i: P2L17: “volume” - not a defined measure. avoid use except for “volume setting” on a device that employs that terminology*

AR-R1-7-1-i: As noted in our response to the Action Editor, we have revised to make sure that its only use is the sense of ‘computer volume setting’ and have flagged this in a footnote as well (page 6).

*R1-7-1-ii-vi:*

*ii. P3L49: “critical bands” - define (jargon)*

*iii. P3L59: “converging” - define (jargon)*

*iv. P4L11: “thresholding paradigms” – in particular, “thresholding” implies setting a limit and/or adjusting values that exceed it on one side or another. Here we are in fact talking about paradigms for measuring perceptual thresholds, which is not the same thing as “thresholding”*

*v. P21L44: “cocor-based comparison” - define*

*vi. P4L39: 57-67 dB - change to dB SPL*

AR-R1-7-1-ii-vi: All defined or clarified

*R1-7-1-vii: P6L13: Sound level meter measures need to indicate weighting (A?, C?, Linear?)*

AR-R1-7-1-vii: A-scale weighting, used throughout, is now consistently indicated

*R1-7-1-viii: P6L32: What are the units of the RMS measure? If digital, can you express this in a more meaningful way, such as dB below maximum?*

AR-R1-7-1-viii: We used RMSv and have been more explicit about this throughout the manuscript. We use dB when referring to differences in RMS or SPL (see page 7).

*R1-7-2: P6L6: “Under well-controlled laboratory conditions” – Using what procedure? The same procedure, just in the lab? Or some other “gold standard” procedure?*

AR-R1-7-2-viii: This information now added.

*R1-7-3: P6L50: what is meant by “continuous loop was achieved by in-house JavaScript”? Do you mean that the experiment presentation code (in JavaScript) implemented a continuous loop for playback? Or are you talking about a separate software program?*

AR-R1-7-3: This information now added (the experimental driver required additional JavaScript to enable the looping) (Page 10).

*R1-7-4: P7L9: Artificial ear measurement: how was this coupled to the DT-150 headphones? Flat-plate adapter or other approach? What microphone was used?*

AR-R1-7-4: This information was added (Page 10).

*R1-7-5: P7L13: Why is it important that the calibration was conducted in an anechoic chamber? Presumably the sound measurement is taking place inside the coupler, and room acoustics should not play a factor*.

AR-R1-7-5: We thought it was important to report ambient noise levels that were measured under conditions as close as possible to what we used for testing. We tested subjects in an outdoor park and in the chamber, so we reported those respective ambient noise levels.

*R1-7-6: P7L26: “about 28 dB SPL” under what settings? A-weighted? Linear? In the coupler? With headphones in place on the coupler (seems high for that)?*

AR-R1-7-6: It is A-scale weighted, and newer measurements are a few dB higher, at 31 dBA SPL. All requested information has been added.

*R1-7-7: P9L41: why is calibration referred to as “the anechoic calibration procedure” if it involves*

*coupler (artificial ear) measurements?*

AR-R1-7-7: Because it took place in an anechoic chamber; we have deleted ‘anechoic’.

*R1-7-8: P10L8: I’m having trouble accepting the use of “audibility threshold” to refer to the adjusted level. I understand that you’ve asked participants to adjust the sound “to threshold,” but I cannot figure out if the value is really a “threshold” if it’s just an adjustment? I think the right language is to say that participants adjusted their “just detectable” levels to certain values, which serve as estimates of the audibility thresholds*.

AR-R1-7-8: Thanks, we have taken this wording and revised accordingly.

*R1-7-9: P10L36043: What about noise level inside the headphone? It seems that if you want to compare the level of task-influencing noise across environments, the level should be measured inside the headphone*.

AR-R1-7-9: While we did not do this, we do now report the amount of attenuation provided by the headphones we used, along with a number of others for comparison. See details in Supplementary Materials Figure S2 and also updates on page 40.

*R1-7-10: P11Fig3B: maybe it’s just the way it’s plotted but the normal fit doesn’t look very good. Do you have statistics to verify whether this is normally distributed?*

AR-R1-7-10: Thank you for pointing this out. The fit now has been removed from Figure 3B as it was irrelevant to the question at hand.

*R1-7-11: P14L39: There is a concrete claim here about canceling the probe signal effect with certain kinds of noise. It needs to be supported with some data or citations*.

AR-R1-7-11: As noted in the initial manuscript, the effect of onsets or intermittent noise on the probe signal effect was observed in our own unpublished data, so this is not citable yet. We have therefore removed that sentence.

*R1-7-12: P14L57-60: I do not understand these values. The SNR starts at negative 13.8 and then is reduced to positive 22.75 and beyond? Should that read negative 22.75?*

AR-R1-7-12: This typo (missing ‘-’) was also noted by the Action Editor and is fixed.

*R1-7-13: P16L43-57: I found the discussion all about “slightly higher than expected” and the long justification confusing. I think it’s fine to just say signal and probe tones were presented 0.75 dB below threshold. It doesn’t need to be justified why you chose that level*.

AR-R1-7-13: Thanks, we have deleted the reference to the pilot results (but did keep in the rationale for raising the threshold slightly as other readers may wonder why we did this).

*R1-7-14: P18L11-13: Find and cite some data to back up “In our experience the range…”*

AR-R1-7-14: We have just taken this out as the paper is long already.

*R1-7-15: P18Fig5C: 0 dB SNR is not a meaningful limit and bars should not be anchored there. Use a boxplot or violin plot instead*.

AR-R1-7-15: We have substituted the bar plot with violin plots. See the new Figure 6C.

*R1-7-16: P19L34: the difference “is relatively small” compared to what?*

AR-R1-7-16: Agreed that this is not helpful, we’ve deleted.

*R1-7-17: P22L20-39: this paragraph presents a cool idea but is confusingly written*.

AR-R1-7-17: Thanks for flagging this, we have rewritten it.

*R1-7-18: P22L47-54: logical fallacy here. You’re affirming the consequent: if signals were set to the same level, no correlation would be expected. We found no correlation. DOES NOT imply the signals were set to correct level*.

AR-R1-7-18: Thanks very much for flagging this, that last line was simply incorrect and should not have been there, now deleted.

Finally, thanks to Reviewer 1 for the many detailed comments.

### Reviewer 2 comments (R2) and Author Response to R2 (AR2)

*R2-0: Summary: This study presented timely investigation on data quality and validation results from multiple datasets obtained online and in other remote testing mechanisms to compare with previous in-lab studies. I am impressed with the compelling results and thorough analyses. However, I think it can benefit from revisions, particularly in streamlining the core analyses necessary to address the questions at hand. Please see my comments below*.

AR-R2-0: Thank you for the positive comments, and also for the very helpful comments for improving the manuscript.

*R2-1: General comment: The paper is very long with many figures (and subplots) with lots of information. Some of the figures can be reduced to drive home the most important messages. In the Results sections, there is much redundancy in repeating motivation that should belong to Introduction/Methods. There were many types of statistical analyses, which may be appropriate. But the presentation of stats, by mixing result interpretation followed by stats reporting, makes it very difficult for me to fully understand what test was actually performed*.

AR-R2-1: In looking over the paper again, we also agree it is too long and would benefit from edits and reorganization (including those you suggest). We have made general changes and also note specific revisions below.

*R2-2: This is a small suggestion mainly on the writing style. There is quite a bit of use of em dash throughout the manuscript that leads to long sentences. It may improve readability to break up these sentences, for instance Page 21, line 42:” … a point we return…” This can be a stand-alone sentence something like “We will further address this issue [onUnderlying mechanism? Your suggestion to resolve? Some indication on what’s discussed later will be good.] in the Discussion.”*

AR-R2-2: Agreed, and we have gone through the paper to shorten and simplify sentences as much as possible.

*R2-3: Page 4, line 27: “…effectiveness of this online approach [by comparing] with in-person…” Page 4, line 27-40: Even though Exp 1 is well motivated, I thought the metric for validation/comparison can be identified more explicitly – it’s the “range of SPL” as tested in each sample. I overlooked this and carried the assumption that the “absolute SPL” was the metric for validation into the results section. I suggest ending the paragraph by what you’d predict in the sample tested outdoor: wider range?*

AR-R2-3: Thanks for your suggestion. We have done this, and updated the manuscript accordingly.

*R2-4: Page 5, line 12: “inexpert” -> “inexperienced” or “naïve”*

AR-R2-4: Done.

*R2-5: Page 5, line 14: I do not understand what is suggested in 2) and 3)*

AR-R2-5: Thanks for pointing out the unclear phrasing, this section is completely rewritten.

*R2-6: Page 5, line 29-36: The motivation of the 3 listener groups is missing. It may be better address in the INTRO in line 27-40 as well*.

AR-R2-6: Here we did not quite understand the reviewer’s comment in that the three groups are the samples from three different experimental conditions (expt 1a-c), conducted on different days.

*R2-7: Page 6, line 20: the information on ethics approval (e.g., international data collection) and informed consent can benefit from additional details at the same level outlined in Exp 2*.

AR-R2-7: Thanks, this was also commented on by the Action Editor, and we have now included those details.

*R2-8: Page 7, Calibration sub-section: I thought this is better suited in the Methods section. I questioned the validity of using 1 kHz tone as a way to confirm SPL linearity as the basis for broadband SPL extrapolation. It seemed like a better approach is to set up the same noise signal at different voltage levels (the same way the pure tone was manipulated) and explore linearity that way for better equivalence*.

AR-R2-8: We have done this and updated the manuscript accordingly. More specifically, we created three versions of the noise signal, at +0, +10, and +20 dB, re the digital level used during testing. We then derived SPL’s for each noise in the chamber, played from the laptop at each volume setting. At +20 dB, SPLs were all above the ambient noise floor, while at +10 dB, all but the lowest volume setting were. At +0 dB, SPLs were above the noise floor down to a volume setting of 44%, and the data collected at +10 and +20 dB were used to extrapolate the rest of the values.

*R2-9:* Figure 2*, panel D: I suggest flipping the color code to follow the same color scheme as other subplots (black=outdoor)*

AR-R2-9: Done.

*R2-10:* Figure 3A*: suggest using different shapes for individual data for Exp 1b/1c*

AR-R2-10: Done.

*R2-11: Page 11, line 40: suggest “consonant” -> “similar”*

AR-R2-11: Done.

*R2-12: Page 12, line 27: not sure what “mode” here refers to (same as in page 4, line 53)*

AR-R2-12: We’ve appended ‘statistical’ here to make it clearer, defining it the first time as ‘the most frequently occurring value’.

*R2-13: Page 12, line 37: The use of “finicky” seems to have a lot to unpack, and promotes readers to question the motivation of even using this paradigm as validation. Please expand on more specific details: need for high-precision audio hardware? Need for consistently behaving subjects? Demonstrate large variability across thresholds measured across studies?*

AR-R2-13: Point taken, and to avoid further lengthening the manuscript we have just taken it out.

*R2-14: Page 13, line 9: access to diverse subject pool is a clear strength. Can you include additional details beyond “around the world”?*

AR-R2-14: We have added the following details: “*As Prolific is available in most of OECD countries except for Turkey, Lithuania, Colombia and Costa Rica and also available in South Africa, most prolific participants are residents in these countries. In our sample, the 60 participants were residents from 13 different countries including United States, United Kingdom, Canada, Poland, Spain, Germany, South Africa, Belgium, Chile, Mexico, Portugal, France and New Zealand*.” (Page 20).

*R2-15: Page 13, line 60: The procedure by Milne et al. 2020 does lead to some attrition. Please report any attrition for this procedure in this implementation*.

AR-R2-15: As the attrition caused by the headphone test has been investigated our previous work (Milne et al. 2020), here we only recorded the participant’s results if they passed the test. Different from our setting in the previous work, here we did not formally record the participant’s headphone test result if they failed it. The failed participants would not be able to continue the experiment and would be told to return the test on Prolific before the main experiment started. To get an overall idea of attrition, we counted how many participants returned the test on Prolific. A total of 91 participants started the test, 7 participants quit the test after passing the headphone test, and 24 returned the test before the main experiment started. However, of these 24 returned participants, it is unclear whether they completed the headphone test or not, as they might have quit even before the headphone test started. Nevertheless, our total attrition for Experiment 2 (including before or after the headphone test and drop-out during the main experiment) is 34.1% (31/91). This information is now included in the manuscript (Page 21)

*R2-16: Page 18, line 3: did every subject reach at least 6 reversals? Can you report the number of reversals reached in general?*

AR-R2-16: The histogram of the number of reversals amongst all 180 tracks (60 participants, each had three tracks) is plotted below. In total, 16 participants had one track with less than 6 reversals (2 tracks with 3 reversals, 1 track with 4 reversals, 13 tracks with 5 reversals).

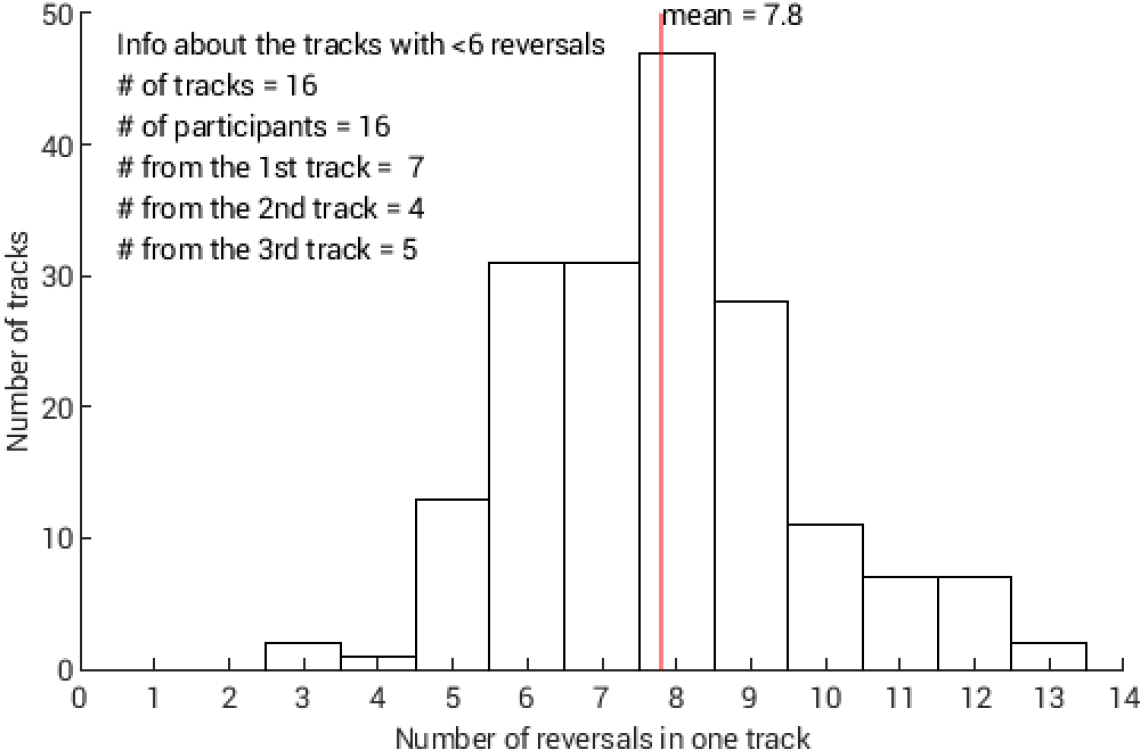

Histogram of the number of reversals amongst all three tracks and amongst all 60 participants. The red vertical line indicates the mean.

Amongst these 16 tracks, the number of reversals did not correlate significantly (pairwise all > 0.05) with age, questionnaire-derived apathy index, baseline fatigue rating, baseline motivation rating, baseline confidence rating, the accuracy of the signal detection in the later probe-signal task, the threshold computed as mean of last six reversals, or the mode-based threshold.

*R2-17: Page 18, line 3-13: A first-pass on fatigue/practice will be to test if there’s an order effect on threshold going from track 1->2->3*.

AR-R2-17: A repeated-measure ANOVA on the mode-based threshold for each track showed that there is no order effect [F(2,118) = 2.30, p = 0.11, partial eta squared = 0.038, observed power = 0.46, Sphericity Assumed]. We have inserted this result in the manuscript (page 25).

For the reviewer’s interest, we also provide this figure showing the distribution of threshold values for each track.

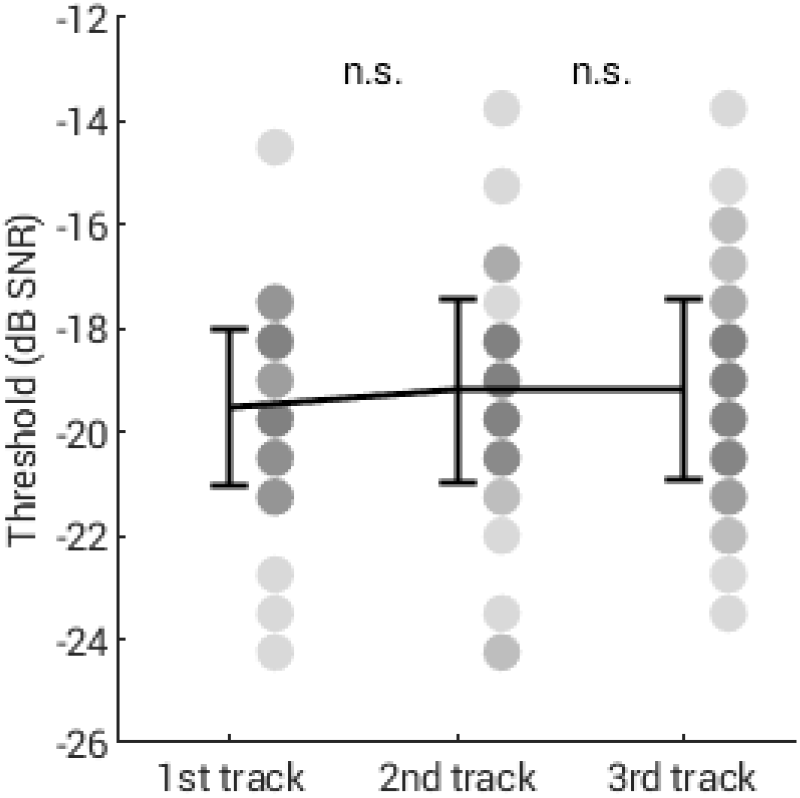

The transparent grey dots indicate individual data for each track. The black line indicates the group average with error bar 1 standard deviation. No significant difference (abbreviated as n.s.) was found between tracks (All p values > 0.16). Bonferroni correction was applied for multiple comparisons.

*R2-18: Page 18, line 28-36: I agree with this approach for validation and the evidence is compelling. I want to offer an additional approach to extract “gold standard” thresholds for validation to strengthen it even further. In human development literature, children are prone to inattention/fatigue. Thresholding using last few reversals is not always the most reliable “gold standard” in such population. To deal with this, there has been approaches to use maximum likelihood estimation to reconstruct psychometric functions using ALL trials in the adaptive track (see “psignifit” Matlab toolbox developed by Wichmann and colleagues:* https://github.com/wichmann-lab/psignifit). *Since online subjects are likely behaving between an in-lab adult and an in-lab child, having thresholds extracted from the reconstructed psychometric functions will further strengthen the validation result. In this case, all 120 trials can be used to reconstruct the psychometric function, which should be fairly robust. Also, please report the number of reversals seen on average when trial #20 was reached in the first track*.

AR-R2-18: Thank you very much for this advice. We have added this method and the results have been reported and compared with the other three methods (Figure 25-26). See the pink violin in Figure 6C.

The thresholds determined by this method is strongly correlated with those determined by mode (Pearson r = 0.94, p<10^-28; Spearman rho = 0.87, p<10^-18). On average, the threshold determined by this method is 0.475 (SD 0.516) dB SNR lower than the threshold determined by mode.

The number of reversals seen on average when trial #20 was reached in the first track was 3.45 (SD 1.05).

*R2-19:* Figure 5C*: Suggest under the green bar use “1st track” instead of “1 track”*

AR-R2-19: Done. Please see new Figure 6C.

*R2-20: Page 21, line 44-52: Do results here refer to the CHANGE in r-values reported in* Figure 7*? Please elaborate on the tests done and additional statistical details beyond p-values. The use of “post hoc explanation” seems odd. This sounds like an interpretation to me. Also, I am not quite convinced that the correlation analyses in* Fig 7 *directly leads to the conclusion stated here. There is smaller individual variability from track 1->3; but it seems that in order to demonstrate convergence of mode-based threshold toward the gold standard, we shall see the deviation from gold-standard performance REDUCES over tracks. So in that sense*, Fig 6 *already provides some of this information. I wonder if there’s a way to combine* Fig 6 *&* 7 *for more condensed/precise presentation. Also, if the recommendation is to use the first 20 trials in a single track, I’d suggest laying additional evidence here by quantifying the effect size of “error”. What is the range of deviation from the gold-standard can we expect from mode-based estimation with just 20 trials?*

AR-R2-20: We have reworked this paragraph to add both detail and clarity. We tried to combine Figs 6 and 7, but did not find a satisfactory layout, so we have kept them separate. We apologize, but we were not sure what Reviewer 2 means by ‘effect size of “error”’. With respect to the range of deviation from the gold standard with mode-based estimation from just 20 trials, this was shown in original Figure 6a (now 7A), with comparisons to mode in 30 trials (Fig 6B), and 40 trials (Fig 6C). To make this clearer, we have now added the reviewer’s compact phrasing to the description in the main text pointing to the figure (Page 26).

*R2-21: Page 22, line 20-54: Some content in these two paragraphs seems to better suit the INTRO/METHOD section for Exp2. If they remain here, please consider rewriting it to be more concise: what is expected, what do we see, and how do we conclude. For instance, the take-home message does not appear until the last sentence line 52-54. I suggest putting this content much earlier to set up the goal for the analysis, then show the analysis. If I am able to read these two paragraphs with the idea already in mind, it’ll help me better understand the justifications listed on the analyses and interpretation of results*.

AR-R2-21: Thanks - we agree this was not very easy to understand. The section has now been extensively rewritten and hopefully it is clearer as a result.

*R2-22: Page 24, line 12-15: Move this paragraph after the next. The analysis has not been shown yet to support this conclusion*.

AR-R2-22: Our larger revisions to this section of the manuscript have made this move unnecessary.

*R2-23: Page 24, line 26-28: What are the “gold-standards” from the literature? Please provide the range of metrics used here from past work to support the claim that present dataset replicated effect/finding. Line 34: Is ∼5% similar to what’s previously reported?*

AR-R2-23: Please see the revised Figure 6, and the text associated with it (page 25). On R1’s recommendation we added an additional traditional measure, and we now describe each in detail.

*R2-24: Page 24, line 42: suggest removing “As exciting…can be, “ This paragraph can be shortened to a single sentence as a simple reminder for motivation*.

AR-R2-24: Thank you. We have completely revised this section for brevity.

*R2-25: Page 24, line 50: this sentence sounds odd. Motivation affects the threshold estimation in a way that leads to poorer/more inconsistent threshold estimation. How does psychoacoustic threshold REFLECT motivation?*

AR-R2-25: We have reworded this sentence.

*R2-26: Page 25, line 19-28: This analysis captures all the analyses and conclusions in the correlations ran and presented from Page 24, line 58-Page 25, line28. Suggest removing the correlation analyses*.

AR-R2-26: In this case, we would prefer to retain the correlation analyses as they are as for some readers they may be more clear than the subsequent analysis, and do also bring out specific effects.

*R2-27: Page 25, line 30-38: Why not run a single GLM that include apathy and fatigue scales? This whole subsection can be reduced into 1-2 paragraphs to drive home the message. The redundant information makes it very distracting for me*.

AR-R2-27: We have reduced this section considerably.

*R2-28:* Figure 9F *schematic was hard for me to follow and confirm from the analyses presented. Each LMM reported in Page 25-26 was only looking at a subset of the relationships in the schematic outlined, without considering the parameters not included in the LMM. Some of these relationships will change when additional parameters are added in the model. Maybe the difficulty here is the reporting style making it difficult to follow*.

AR-R2-28: We agree that Fig 9F was not particularly helpful, so have removed it and also revised the text (see Figure 12 and page 36-37).

*R2-29: Page 25-27: The most useful information again appears in the last paragraph. This is the content that directly supports the subsection heading. I suggest removing/reducing the earlier paragraphs for concise presentation*.

AR-R2-29: This subsection has been largely trimmed.

*R2-30: Page 28, line 46: “outlier-robust” was never mentioned or supported in the Results section. Need evidence to support mode-based is robust to outlier. Also, what kind of outlier data is referred to? Attention lapses on individual trials, or individual listeners with outlier threshold?*

AR-R2-30: A fair point -- we have removed mention of “outlier-robust.”

*R2-31: Page 28, line 50-59: I believe this is the place where you promised to return to in the Results section. While I agree that mode-based threshold estimation is a good alternative, one more point to consider is how do you resolve bi-/multi-mode estimation? And even if a single mode is arrived, what does the distribution look like with SNRs with 2nd/3rd frequency? In other words, will a track with 15/20 trials tested at 0 dB SNR suggests different reliability from another track with more disperse SNRs? How does more disperse SNR distribution affect threshold estimation? This is worth discussing even though there may not be direct implication from the data just yet*.

AR-R2-31: We agree these are important questions to resolve (and have inserted a note regarding multi-modal estimation, thank you), but as the reviewer points out, we cannot really make any inferences from our data or procedures currently.

Finally, we would like to thank Reviewer 2 for taking the time to make such constructive and helpful comments.

### Authors’ Response Letter — Revision Round 2

We briefly respond (Author Response, *AR*) to each of your (editor in chief, E-i-C), the action editor’s (AE) or the reviewers’ points (R1, R2) here. Please extend our thanks for their helpful suggestions and feedback.

### Editor in chief’s comments (E-i-C) and Author Response (AR-E-i-C)

E-i-C: As you will see, everyone agrees that the revision is a great improvement, to the extent that both current reviewers are happy with this version as is. The Associate Editor has gone through the manuscript carefully himself and provides a number of comments that I think will be helpful in further strengthening the paper before its publication. Please consider all the comments carefully and provide what I hope will be a final revision that addresses them.

*AR-E-i-C: We are pleased that the reviewers and Action Editor find the manuscript improved and interesting. We respond to all points below, noting where we have made changes in the manuscript or supplementary materials*.

### Action Editor’s comments (AE) and Author Response (*AR-AE*)

AE-1: Although this MS is a big improvement on the one previously submitted, it is still has a large number of places where greater clarity is required.

AR-AE-1: *We are happy that the action editor has found the manuscript improved, and have taken the suggestions on clarity on board in our revisions as detailed below*.

AE-2: The IRB/ethics process only is given for online experiments, yet in-person experiments were conducted at Carnegie-Mellon.

*AR-AE-2: All of our experiments were conducted with ethical approval, as required. Please see detailed response below*.

AE-3: I am still not convinced that the authors have measured the noise floor of the headphone setup as they think they have, and what it means perceptually. Their quoting of headphone dBA appears to be wrong: it is not referenced to dBA in the diffuse/free field but on the headphone output measured in a coupler. Ie referenced to the ear drum.

*AR-AE-3: We have added the following to the manuscript: “Note that the coupler simulates ear canal resonance, which when paired with A-Scale weighting, magnified the associated band-pass filtering and thus likely underestimated SPL at the eardrum.” (page 10) Again, more detailed responses to the AE’s points on this topic are below*.

AE-4: There are hidden factors, such as the ear-defender effect of headphone styles that are left right to the Discussion before being mentioned as to being a potential source of online variability.

*AR-AE-4: The previous comments from the Associate Editor and at least one of the reviewers emphasized that our Introduction and Methods were too verbose, and that interleaving results and discussion was confusing. Thus, the placement of this text was based on the Associated Editor’s prior comments*.

AE-5: Additionally, there is an implicit assumption that headphone responses are near-similar across brands. Having measured many headphones and earbuds, I can say that this is not true. I give an example below, and attach some figures to illustrate.

*AR-AE-5: In fact, we took pains to acknowledge and demonstrate that headphone responses are not similar; the manuscript does not imply or assume that all headphones or earbuds are equivalent. Following the first reviews, we undertook additional testing of multiple headphone brands of varying quality and price points. As we noted in the manuscript, this was not an exhaustive survey*.

AE-6: The use of a pink, rather than white noise to “validate” their threshold measure is still a bit odd: relying on any consistency in response across headphone designs above the 1-2 kHz region is naive (I attach plots of measured response from real headphones, as well as intended diffuse field response). The meatal resonance would determine that any thresholds with such noises would be dominated by detection in the 2-3 kHz region, for properly designed headphones. However, the signal goes up to 8kHz, ie into a frequency range where there is vast inter-headphone variability. To use the threshold obtained in such a way so as to, as the authors do, add in 40 dB so as to achieve what they think of as a reasonable supra-threshold level for testing is very dangerous when they intend to use a signal frequency (or band of frequencies) that are remote from the 2-3 kHz region. There is no caveat here, which would require the experimenter to check that the level is acceptable to the participant.

*AR-AE-6: Thank you for the suggestion. We have added a statement to caution that this method should not be used when stimuli present narrow frequency ranges at high power in spectral regions distant from 1-4kHz*.

AE-7: The use of/description of the reference level of the signal amplitude in RMSv is a mess: it appears to be both voltage and digital file rms. P10 line 11 “This level was chosen as pilot testing suggested it allowed quiet” It is usual to put the units after the number (here Volts, p6 line 28 “particular root-mean-square (RMS) voltage amplitude (RMSv)”). But the statement depends heavily on the headphones’ sensitivity, which can easily range over 20 dB. (95 – 115 dB SPL/Volt), not really covered by the generalities of p11 line 21. (Low sensitivity headphones are demonstrated in Stone, Harrison, Wilbraham, Lough, Trends in Hearing 2019, https://journals.sagepub.com/doi/full/10.1177/2331216519889232). But this level description is at odds with p10 line 21 “This RMSv level of this stimulus file serves as the amplitude”. Well if it is in Volts, then one needs to specify the laptop volume setting that translates a digital signal into a voltage. And again, this is at odds with the later description of the calibration procedure which, because “white noise stimulus was digitally increased in amplitude by 10 and 20 dB,”, can only happen if the Gaussian noise digital RMS starts a about −30 dB RMS (NOT SPL) in the file. Insufficient and confused description. (The .000039 figure is unnecessarily repeated on p 11 line 23)

*AR-AE-7: We have changed these to RMS except for the cases where we have specifically measured output voltage*.

AE-8: The plotting of the background noise spectra, with a linear frequency axis, is unhelpful: it places too much emphasis on the 10-25 kHz region where there is no signal and where headphone response is very varied. Figure S2: audio engineers may use linear frequency scales, but here we are talking about perception, Most of the “interesting effects” (ie perceptually relevant) are confined to just 1/5th of the abscisaa, 0 to 5000 Hz, especially since the noise was bandlimited to 8000 Hz, and downward spread of masking is pretty negligible in people with near-normal hearing.

*AR-AE-8: We have changed the axis to log in the revised manuscript*.

AE-9: /P 3 line 8 No hyphen between audio & equipment.

*AR-AE-9: Thanks, fixed*.

AE-10: P 3 line 35 “for asking new auditory neuroscience questions quickly”. No, it is “answering”.

*AR-AE-10: We are asking questions. We hope to answer them. Asking is certain, answering unfortunately less so*.

AE-11: P 3 line 37 “diverse samples”. More like “diverse participants”.

*AR-AE-11: We intended <u>diverse samples</u>. We are not discussing the diversity of individual participants*.

AE-12: P4 line 7 “due to a half-century of” No, probably more like nearly a century, if we go back to Harvey Fletcher at Bell labs in the 1920s.

*AR-AE-12: Thank you for the historical perspective; now corrected*.

AE-13: P5 line 21-23 “so often experimental sessions will begin by running adaptive psychophysical paradigms to estimate the individual’s relevant perceptual thresholds” This sentence is misleading. One can read it as “the experiment begins……and then….” But the “and then” does not appear. The use of an adaptive track of some form (maybe modified by statistical decision) is fairly fundamental otherwise the listener will not know what to listen for, and generate spurious data. Hence there has to be adaptation to (or around threshold). (or derive psychometric functions in an efficient way)

*AR-AE-13: Unfortunately, we were unable to understand this comment, so have left the phrase as is*.

AE-14: P5 line 26, P10 line 11 (and possibly elsewhere) “quiet thresholds”. “thresholds-in-quiet” is preferred term.

*AR-AE-14: All instances now using this phrasing*.

AE-15: P6 line 31 : “setting task in uncontrolled and several laboratory environments” Needs commas (or expanding): “setting task in uncontrolled, and several, laboratory environments”

*AR-AE-15: This was indeed unclear. Now edited to ‘…in uncontrolled environments and in the laboratory’*.

AE-16: P6 line 21 onwards to p7 line 24: Is this level of detail really necessary for the IntroductionIn that section, one is trying to establish the content that follows, so a broad over-view of the experiments is necessary. Additionally it is nice that the authors show how one experiment builds on a preceding one. But by the time we are down to describing participants’ actions, as well as tracking percentages, this is best left to a relevant part of the “Methods” section. And, as they have done, finish off with a mission goal.

*AR-AE-16: Thank you for this comment. We thought carefully about this and ended up retaining the text in that the specific tasks and especially the probability manipulations (tracking percentages) are crucial to understanding the probe signal experiment, which may be unfamiliar to many readers*.

AE-17: P7 forwards. In the previous version I raised the issue of ethics/IRB approvals. The reply was “Approval was obtained in the UK for the study to be conducted online, with the express understanding that anyone could participate in any country, as is now typical for online studies conducted across multiple platforms. Clarification has been added (page 9).” which at first appears acceptable. However, p 7 line 47 onwards, we are told that, for Expt 1, the participants are “Carnegie Mellon University affiliates”, and they are being tested face to face (in-person solicitation, outdoors, campus, park, indoors) with prescribed equipment. P8 line 55 “Validation of the online amplitude setting procedure required testing in-person participants”. It is not indicated that IRB approval extended that far.

*AR-AE-17: Face-to-face participants were covered by local CMU or Pitt IRB protocols, but also by the UK ethics in that the experiments all used the online materials. Nonetheless, the revised manuscript makes note of the ethics/IRB coverage*.

AE-18: P 8 Table 1 : poor formatting with numbers split over 2 lines, what does “Measurement” refer to as a column heading?

*AR-AE-18: This is the formatting required by the journal for submission. The table will be reproduced by the copy editors, so we have not made changes to it. Regarding ‘Measurement’, we have made the heading for the list of sound level meters more explicit by renaming it “Measurement Apparatus”*.

AE-19: P10 line 11 : SOX, FLAC, A bit too much detail over “signal files were generated and stored in a lossless format and 16-bit precision”.

*AR-AE-19: We have left this detail as it is important for online testing; for instance, as the action editor would doubtless agree, the frequently used mp3 lossy encoding can have deleterious effects on reproduction quality*.

AE-20: P10 line 54 : “SPL of the stimulus at the lowest volume settings was below this noise floor,” As I pointed out in my previous review, there are two noise floors to consider, (a) the external sound field noise floor (“Ambient noise floor”), as well as (b) the electrical noise floor of the output stages of the computer (p11 line 24 “noise floor of the sound card”). The writing over the next few lines is ambiguous as to which floor is being considered. Additionally, the quoting of a 31 dB A SPL noise floor (p10 line 52) does not specify the reference conditions. Ie was the B&K meter connected to the coupler at the time, or was it measuring the diffuse field near the coupler, outside of the headphones. Really one ought to be measuring the ambient noise as that which leaks into the coupler, ie with the headphone connected physically, but not electrically. P10 line 54 does say “this noise floor” so one could infer the ambient floor, but p11 line 5 and line 11 only mention “noise floor”. P12 line 34 uses “noise floor” for the electrical system noise.

*AR-AE-20: This is, in fact, what we did. The Action Editor may have missed this, but we wrote that “…measurements were obtained using a Bruel & Kjaer 2231 precision sound level meter set to slow averaging and A-scale weighting and Bruel & Kjaer 4155 ½” microphone mounted in a Bruel & Kjaer 4152 artificial ear with a flat-plate coupler, coupled to the same set of Beyer DT-150 headphones used for data collection.” (page 10)*

AE-21: But again, we have an issue here: the SLM readings are quoted in dBA, even when the microphone is located in the coupler. This is not true. The dBA setting is a physicist’s approximation to the sensitivity of the human ear (excluding the meatal resonance) and is applied to free-field measured signals. The fact that it is reported here, as measured in a coupler, adds in the meatal resonance since the SPL measured is referenced to the eardrum. The A-weighting then applies low & high frequency cuts, to approximate what finally gets through to the cochlea (but ignores the meatal effects). The wrong reference point has been chosen for the measure.

*AR-AE-21: We acknowledge the Editor’s point and have made a note in the paper to this effect. (page 10)*

AE-22: P11 line 5 “Volume setting adjustments were determined to be linear on the Macbook Pro”. No, they were near-linear in dB with Volume %-age, ie a near-logarithmic taper on the “volume control”.

*AR-AE-22: We have retained the clarifying second phrase not noted here by the AE, “e.g., a given increment in volume setting generated a relatively consistent change in dBA SPL at both high and low overall levels.”*

AE-23: P12 line 28 “and compared the power spectrum of line noise alone” The phrase “line noise” is usually used to refer to the mains-electricity-related electrical interference. Yes, the output of a computer can be called the “line out”, so a confusion can arise. In fact the authors are not measuring mains-related noise as can be seen from the supplemental materials. Additionally, line 25 in the same para “signal would be distorted due to low bit depth,”: it may be true that there would be distortion from quantising noise, but given that the test stimulus is white noise, the distortion would be self-masked (triangular psd around each component of the noise, so adding up across all frequencies as just being a bit more noise).

*AR-AE-23: We do not anticipate that readers of Trends in Hearing will be confused on this point*.

AE-24: P12 line 50 “via wireless connections to various broadband providers.” In the context of this experiment, “wireless connections” would usually mean headphones, which would raise a few eyebrows about lossy compression. But no, we are talking about Wi-fi/radio connections. Not really relevant here….. until Expt 1d where we are told that Apple Airpods were used (which generation) which are convey audio via a perceptually lossy connection. This rather goes against all the earlier descriptions/intentions of loss-less audio.

*AR-AE-24: Re the latter point, please see our strong suggestion to readers to have participants avoid using Apple Airpods. Regarding wireless connections, we trust that readers would understand that ‘wireless connections to broadband providers’ will refer to computer internet connections*.

AE-25: P15 line 50 “seems reasonable despite the relatively large difference in ambient noise levels (31 dBA SPL indoors, and 57 dBA SPL outdoors).” This statement is pretty meaningless unless we know (a) the attenuation characteristic of the headphones, and (b) whether the ambient noise was measured in identically the same way (ie confirming that the coupler was not used for this, see above).

*AR-AE-25:* As described in the manuscript, the noise was measured identically, please see below for discussion of attenuation.

AE-26: P20 line 16 “and of these 31 dropped” Comma after “and” and also “these”.

*AR-AE-26: We agree commas were needed but added in slightly different positions*.

AE-27: P21 line 47 “participants returned the test” : “returned” is an unfamiliar term, and seems to be a phrase from Pavlovia, as becomes apparent in later lines. “completed” ?

*AR-AE-27: ‘Returned’ has a distinct meaning for online recruitment and experimental platforms so we have retained it for precision*.

AE-28: P22 line 9 : FLAC & SOX. repetitious of earlier

*AR-AE-28: This is the description of stimulus creation for the pure tones which has not been presented earlier, so not a repetition*.

AE-29: P22 line 32 “and looping through the end of the experiment.” Do the authors mean “and looping through to the end of the experiment.” ?

*AR-AE-29: We have substituted ‘until’ for ‘through’*.

AE-30: P22 line 35 “as several the experimental” Huh ?

*AR-AE-30: ‘of’ has now been inserted between ‘several’ and ‘the’*.

AE-31: P22 line 41 “noise mask”. “noise masker” is the preferred term.

*AR-AE-31: changed*

AE-32: P22 line 29 “for experimental drivers” Where did this phrase come from: it makes no sense in the context.

*AR-AE-32: As the action editor highlights, the term ‘experimental driver’ may be unfamiliar, so we have substituted ‘experimental presentation software’ instead*.

AE-33: P22 line 47 “250-ms intervals (250-ms ISI)” Errr no. Two completely different use of the word “interval”. ISI refers to the gap between stimuli intervals. So the particular interval needs to be labelled explicitly to avoid confusion with the next occurrence of “interval”.

*AR-AE-33: We now make this explicit with a new parenthetical: “* (There was 250 ms of silence between each response interval).”

AE-34: P24 line 32 “The questionnaire and ratings were added to the experiment on the second day of data collection.” Where have we been told about “days of data collection”Is this per subjectI do not think so: it appears to be an omission of the survey from the experiment design and added as an after thought. Was it given ethical approval for this change(departure from agreed procedure).

*AR-AE-34: We have raised this statement (amongst others) with the Editor-in-Chief*.

AE-35: P26 line 52 “gold standard” only defined, and there incompletely, in caption to figure 7. Needs to be in text, at least.

*AR-AE-34: Agreed, and added in the initial discussion of thresholding*.

AE-35: P28 figure 8 : It is all very well using Pearson correlations, but for this sort of data comparison (via two different methods) one really needs inter-class correlation coefficients (ICCs) which are more <u>robust.eg</u> Figure 8, Track 2, light green crosses “flip” from being near or above correlation lines, to being below line, for mode-derived thresholds being above −20 dB. ICCs would help distinguish the degree of robustness in repeatability between the different measures.

*AR-AE-34: We stand by our analysis, as it addresses the question we asked*.

AE-35: P32 line 21: “ensure”, not “assure”

*AR-AE-35: Our understanding of the meanings and senses of ‘assure’ and ‘ensure’ accords with that of the OED, rather than that of the action editor, so we have elected to retain our usage*.

AE-36: P32 line 26 “set at 14 dB SNR” Unnecessary use of SNR here, as it is used in the next line.

*AR-AE-36: we edited this line*.

AE-37: P35 line 20 “T. J. Green & McKeown” incorrect reference format with initials. Ditto p41 lines 43, 54, 56 etc)

*AR-AE-37: fixed*

AE-38: P36 line 54 : “An LMM on confidence rating showed that as the task progressed, confidence decreased slightly, but not significantly so (F(1,595) = 3.50, p = .062),” Since there is no significant change in confidence (p = 0.062), then it cannot have “decreased slightly”.

*AR-AE-38: The manuscript makes clear that the result is not significant*.

AE-39: P38 line 23 “many perceptual and cognitive phenomena are robust to level changes (Moore, 2013)”. Yes, I would agree, but many are not, a point which the authors make a few lines down line 34 “can vary systematically as a function of sound pressure level (Moore 2013…..)”. so the same reference is being used to argue both sides of the coin. Given especially that the reference is a text book, if this is to be done, one should cite the relevant page.

*AR-AE-39: That’s right. We are not in disagreement here. We do not see the need to cite specific pages. The same paragraph provides reference to empirical papers to support these points. We leave it to the interested reader to dive into Moore’s excellent book for full details, should they be interested*.

AE-40: P38 line 50-51 : again another instance where ICCs might be a better statistical tool to illustrate the point. (r = 0.7 to 0.8 indicates that over 1/3rd of the variance is still unaccounted for despite the high significance).

*AR-AE-40: We view accounting for ⅔ of the variance in an online test with naive listeners to be a positive outcome*.

AE-41: P38 line 56 “Given the ∼30 dB difference in average ambient sound levels,” Well this is not proven with the existing text : what is the headphone attenuation effect for each background noise, and, in a quiet room, does this render some of the background noise inaudible. In which case, the two soundfields are not comparable perceptually : one is a masked thresholds and the other is an unmasked threshold, so the 30-dB difference in background noise (actually 26 dBA) is misleading, more likely to be around 15-20 dB. Consider the effect of these before one starts hypothesising about “we suspect that this is due in part to participant strategy in listening in gaps, averaging in time, or in-stream segregation”, which may play a role, but secondary to masked/unmasked thresholds (and change in spectral shape).

*AR-AE-41: We added the following text: The spectra of both background noises have generally low-pass characteristics, so headphone attenuation should not be appreciably different; thus, we measured attenuation in only one of the backgrounds (the anechoic chamber). (pages 38-39). Data on headphone attenuation are presented as previously in Figure S3*.

AE-42: P39 line 36-37 “For example, avoiding both narrow-band stimuli like tones as well as stimuli that are not bandlimited like broadband noise will limit the effects of across-subject hardware frequency response differences on results.” This is only true to a limited extent. I have measured headphones responses where the meatal resonance peaks at 30 dB in Kemar, or as low as 5 dB. A broadband signal (such as white or pink noise, essentially becomes a band-limited signal with the former since the band level around 2-3 kHz is nearly 20 dB above intended. It is not uncommon to find headphone responses that have step changes in output of 10-15 dB in unexpected frequency regions. With such devices, one does not know what part f the broadband noise caused threshold, and therefore blindly adding 40 dB in level to get to a comfortable level is fraught with danger.

*AR-AE-42: See comment 41*

AE-43: P39 line 46 onwards: paragraphs on headphone choice, such as closed back to reduce influence of background noise. : all this should come earlier.

*AR-AE-43: As authors, we believe this fits appropriately into this section*.

AE-44: P40 line 6 : “These data are shown in Supplementary Materials Figure S2.” Actually in is S3. But these attenuations are meaningless since it depends on their shape. One expects them to be primarily high-frequency occluding, so their utility will depend on the spectrum of the background noise, which tends to be low frequency in real environments. Quoting attenuations to typical background spectra might be more relevant here.

*AR-AE-44: We have edited the manuscript to address this proviso on pages 38-39, and we have fixed that typo*.

### Reviewer 2 comments (R2) and Author Response to R2 (AR2)

R2-1: Thank you for the careful revision on this manuscript and responses to my comments in the author response. My concerns have been resolved. I recommend publication for this work.

*AR-R2-1: Thank you. We appreciate your expertise in helping us to improve the manuscript*.

### Reviewer 3 comments (R3) and Author Response to R3 (AR3)

R3-1: While I was not an original reviewer, I have carefully examined the original submission, the response to reviewers, and the revised manuscript. I find the responses to be very attentive to the points raised, and the current version is fully acceptable.

*AR-R3-1: Thank you. We appreciate your expertise in helping us to improve the manuscript*.

R3-2: I have a few minor comments, but they are not essential to add to the manuscript unless the authors find them useful. First, I found the method of “measuring below the noise floor” to be useful and appropriate, and would draw the authors’ attention to the literature on this topic, which might be useful to consult and perhaps mention.

Whittle, L. S., & Evans, D. H. (1972). A new approach to the measurement of very low acoustic noise levels. Journal of Sound and Vibration, 23(1), 63-76. https://www.sciencedirect.com/science/article/abs/pii/0022460X72907894

Ellingson, R. M., Gallun, F. J., & Bock, G. (2015). Measurement with verification of stationary signals and noise in extremely quiet environments: Measuring below the noise floor. The Journal of the Acoustical Society of America, 137(3), 1164-1179. https://asa.scitation.org/doi/full/10.1121/1.4908566

*AR-R3-2: Thank you for pointing out these papers, we were not familiar with either, and they are indeed helpful, so have pointed the interested reader toward them in the relevant section*.

R3-3: Similarly, while there are a large number of citations of probe-tone detection experiments, the theoretical value of the approach is not clearly articulated. I understand that the goal of this work was to show that such experiments can be replicated with remote testing methods, but it might still be useful to discuss the role of attention and motivation in the context of some of the probe-signal experiments specifically focused on these issues. In particular, it is important to note that the representation of the signal frequency was so readily established in these naive listeners. This supports models of attention in which the system adjusts its sensitivity very rapidly. A few relevant references follow.

Hafter, E. R., Schlauch, R. S., & Tang, J. (1993). Attending to auditory filters that were not stimulated directly. The Journal of the Acoustical Society of America, 94(2), 743-747. https://asa.scitation.org/doi/abs/10.1121/1.408203

Fritz, J., Shamma, S., Elhilali, M., & Klein, D. (2003). Rapid task-related plasticity of spectrotemporal receptive fields in primary auditory cortex. Nature neuroscience, 6(11), 1216-1223. https://www-nature-com.liboff.ohsu.edu/articles/nn1141

*AR-R3-3: We agree! These methods were developed to support our ongoing work using the probe signal to examine auditory attention. Since the goal of this manuscript is primarily methodological, we have not gone into detail on the theoretical and mechanistic issues that can be addressed. However, we expanded our discussion and included these references*.

## Authors’ Response Letter — Revision Round 3

Dear Dr. Oxenham,

Thank you for your letter and Action Editor’s comments on our revised manuscript “Robust and efficient online auditory psychophysics” by Zhao et al. We have addressed every point that the action editor (AE) made in the PDF. For the six points which require further explanation, we pasted the corresponding manuscript paragraph, action editor’s comment and provide our responses here. Thank you for your helpful suggestions and feedback.

With best regards, and on behalf of our co-authors,

Dr. Sijia Zhao

### Action Editor’s point 1

Pg 23: “Simultaneous presentation of a long masking sound - or indeed any long continuous sound - is challenging for experimental presentation software, particularly online. However, transient noise onsets and offsets - for instance, starting and stopping the noise masker for each trial - can have surprisingly large effects on perception (e.g., Dai et al., 1991; Franosch et al., 2003). “

Action Editor: I’m not sure how much that applies here for a simple tone-in-noise task. There is the “overshoot” effect, but that is limited mostly to brief tones (< 100 ms), gated on with the first 50-100 ms of a masker. Also, it is stronger for higher frequencies (2+ kHz), which makes it less relevant here. As far as I can tell, the Dai et al paper doesn’t address the question of gated vs continuous maskers. I also don’t really see how the Zwicker tone illusion (Franosch et al., 2003), which relies on a noise with a spectral gap, plays a role here. So long as your target tones were never presented in the gaps between the noise loop, I think you can delete everything from “Simultaneous presentation of a long…”

Author Response: Thank you very much for finding this error in the reference list: it should have pointed to Dai & Buus (1991) and not Dai, Scharf, & Buus (1991). The former paper is (highly) relevant and one we deeply regretted not having known about when we were piloting the procedure. Dai & Buus showed that gating the noise masker essentially eliminates the probe signal effect (see Fig 1) - an effect we inadvertently reproduced in piloting the probe signal with a such a gated noise masker. Therefore, we feel it is important to help others avoid this problem. The Franosch paper provides a potential explanatory mechanism for this (suggested to us by Roland Schaette and David McAlpine) but as it is out of the scope of the article to provide a fuller explanation, we have deleted the reference.

The manuscript has been updated (page 22): “However, transient noise onsets and offsets - for instance, starting and stopping the noise masker for each trial - can have surprisingly large effects on perception, with Dai & Buus (1991) showing that use of noise bursts versus continuous noise maskers essentially eliminates the probe signal effect (Dai & Buus, 1991).”

### Action Editor’s point 2

Action Editor: Unusual to cite in terms of overall level. Spectrum level of the noise (dB SPL/Hz) in its passband is more common. That would allow you to compare your results to classic literature in the field (eg Reed and Bilger, 1973, JASA).

Author Response: We have seen both in the literature - given none of the reviewers mentioned this, we have chosen not to change because it may cause more confusion (it is not very intuitive for people unversed in that particular literature).

### Action Editor’s point 3

Pg 27: “To quantify the degree of variability associated with the number of trials used to calculated the threshold, we compared the distributions of differences between the 3-track grand average and single-track thresholds calculated using the mode of 1) the first 20 trials; 2) or 30 trials; 3) all 40 trials; or 4) the mean of last six reversals.”

Action Editor: Doesn’t averaging an odd number of reversal points lead to bias? Perhaps the averaging should be based on the largest even number of reversals available (2, 4 or 6).

Author Response: If there were a bias, we would have expected to have detected it in the fairly exhaustive comparisons in the section following this one, but we did not.

### Action Editor’s point 4

Pg 39: “The spectra of both background noises have generally low-pass characteristics, so headphone attenuation should not be appreciably different; thus, we measured attenuation in only one of the backgrounds (the anechoic chamber).”

Action Editor: Given that the headphone attenuation is linear, it shouldn’t make any difference what the nature of the background sound is, so you are justified in measuring the attenuation in just one scenario, regardless of the ambient noise spectral characteristics.

Author Response: Agreed, and the sentence now reads: “Given headphone attenuation is linear, and the spectra of both background noises generally low-pass, we only measured attenuation in one background (the anechoic chamber).”

### Action Editor’s point 5

Pg 40: “For instance, if it is expected that stimulus presentation will be in somewhat noisy households at ∼ 60 dBA SPL, the average stimulus dB RMS should be ((60 - [median threshold in dB SPL = 24]) + [dB RMS of white noise stimulus in threshold-setting procedure = 26]) = 62.”

Action Editor: I don’t fully follow the logic here. Where do the numbers 24 and 26 come from, and won’t threshold depend on the spectral characteristics of the background noise(s), which might be different indoors and outdoors, not just the dBA level?

Author Response: Thanks much for flagging the serious lack of precision here; we have revised this completely, and also incorporated more information from the experiments added in the revision. Regarding the potential effect of spectral differences in indoor and outdoor environments (which we measured and have provided in supplemental materials), this would require additional experimental manipulations. However, the average noise detection threshold difference between these distinct environments was on average 4.7dB; a change in noise from white to pink evoked only a 2 dB average difference in threshold. Thus, while there may be some effect of the background scene’s spectral characteristics, these results suggest they will be quite minimal relative to the ∼25dB range in hearing thresholds that is the limiting factor in the precision of setting dB SPL using this method.

This has been updated in the manuscript (pages 39-40): “For experimenters who need to present auditory stimuli within a given range of intensities or at a particular level above perceptual threshold, the presentation level can be referenced to the RMS level of the white noise stimulus used in the amplitude setting procedure. For instance, say an experimenter wants to set her stimulus presentation level at ∼ 60 dBA SPL. an average of. If she assumes the typical participant will be in an acoustic environment similar to the outdoor setting (with an average 50 dBA SPL ambient noise level) the average stimulus soundfile *RMS* to produce an average 60 dBA SPL level in the headphones can be estimated. Recall that the RMS of the white noise file used in Experiment 1a (background noise level ∼50dBA) was 0.000399; using this stimulus, participants set their thresholds to an average of 29.4 dBA SPL (range 22.3 – 35.6 dBA SPL).

Assuming our Expt 1a-c noise detection threshold results generalize to the online population, the 10^th^ and 90^th^ percentiles of presented levels across all participants should be approximately 54- and 68-dBA SPL. Alternatively, the stimulus RMS could simply be scaled 30 dB above each individual participant’s white noise threshold level to ensure that stimuli are sufficiently audible to the vast majority of participants. One very important caveat to this approach is in the case where the spectrum of experimental stimuli is far from the 1-4kHz band that will drive much of the detectability of the white noise stimulus (for instance, pure tone stimuli at lower or very high frequencies). Here, it is important that either additional checks be placed on stimulus amplitude, or that a different thresholding stimulus be used (for instance a narrower-band noise centered around the stimulus frequency). “

### Action Editor’s point 6

Pg 44: “It is interesting that motivation showed a small influence on the probe signal effect, but not on threshold estimation.”

Action Editor: Be careful here - unclear if motivation influenced the effect, or if both were related to time of experiment. Also, there is a history showing the relative insensitivity of simple tone-in-noise detection to motivation, reward, etc., going back to the 60s (eg Watson, C.S., Clopton, B.M. Motivated changes of auditory sensitivity in a simple detection task. Perception & Psychophysics 5, 281–287 (1969). https://doi.org/10.3758/BF03209563) that would be worth mentioning briefly.

Author Response: Thank you for pointing out this relevant reference for us. This reference has been added into our discussion. It’s in line with our finding that motivation influences the auditory sensitivity, but the influence was rather small. Please do note that this statement is made based on the LMM analyses that included time-on-task (block index) along with the fatigue and motivation variables, so is well-supported.

The corresponding paragraph has been updated: “One might worry that the online threshold estimation may be affected by the on-task motivation of the participants. It is interesting that, at least that in this study, we did not observe any influence of self-reported motivation on the threshold estimation amongst the remotely tested participants. On the other hand, motivation showed a small influence on the probe signal effect. This is in line with the previous work (Watson & Clopton, 1969) which found motivation — regulated by applying electric shock on incorrect trials— increased sensitivity in a simple tone-in-noise detection task but the increase was rather small. One explanation for the absent effect of motivation on the threshold estimation here is that the effect of motivation on performance is sensitive to the length of the experiment; the probe signal experiment was longer (around 20 minutes) and was run after the threshold estimation (a length of around 10 minutes). This, with no observed effect of motivation in Expt 2, indirectly suggests an advantage of keeping experiment time shorter. In summary, any generalizations of the motivation-related findings here should be taken carefully.”

## Footnotes

1 Note that we use the term ‘volume setting’ to refer exclusively to the computer controls which are labelled as such; otherwise ‘level’ is used.

2 Note that this is a very weak form of statistical inference. To our knowledge, a formal test for an interaction between differences in correlations within and across levels of a repeated-measures design has yet to be developed, so the reader should not infer a significant interaction from these pairwise tests. We have also not applied any correction for multiple comparisons.

3 An LMM was used to investigate the effects of the current block’s fatigue rating, the previous block’s confidence rating, and task progression on motivation loss. Unsurprisingly, longer time on the task (*β* = −0.41, F(1,544) = 7.78, p = 0.0055) and higher fatigue (*β* = −0.29, F(1,544) = 66.08, p < 10^-14^) were associated with sharper motivation loss. Confidence, on the other hand, appeared to exert a restorative effect on motivation loss (*β* = 0.249, F(1,544) = 31.31, p < 10^-7^). Adding the questionnaire-derived apathy index to the LMM revealed that apathy counteracted the restorative effect of confidence (*β* = −0.0078, F(1,521) = 10.22, p = 0.0015). That is, in motivated individuals’ high confidence more strongly prevented motivation loss over time, while in apathetic people this effect was diminished.

