## Supplemental Materials for "Robust and efficient online auditory psychophysics"

### Comparison of experimental signal versus line noise in two laptop setups.

We used the line in of an early 2011 MacBook Pro running OSX 10.9.5 and Adobe Audition CS5.5 to record the analog headphone jack output of a 2020 MacBookPro (M1 chip) running macOS 11.6.1 (44.1kHz sampling rate/16-bit depth). The white noise (Expts 1a-c) and pink noise (Expt 1d) stimulus were played via the experimental setup used for participants (below), at each of MacBook Pro volume settings reported below as perceptual detection thresholds. Each recording started ~5s before the 'play' button was clicked; the pulsed white noise was allowed to play for another ~5s; recording was stopped after a final 5s. To evaluate the signal quality of an older and less-expensive laptop (potentially more characteristic of the hardware available to online participants), we used the same setup to record the analog headphone jack output from a ~2014 Asus X550C laptop running Windows 8 and Google Chrome. White noise perceptual thresholds from two observers were used to estimate analogous volume settings between the Asus and MacBookPro laptops.

We analyzed the input power spectrum of each recording using Adobe Audition CS5.5 and Matlab 2020b. Stimulus recordings used the ~5s after the initial signal spike from the sound card being engaged (more prominent in the MacBookPro). The noise floor power spectrum at each laptop volume setting was established using the last 5 seconds of the recording.

Figure S1a-e shows the spectral power (in dB from peak amplitude) for white and pink noise for each computer around threshold volume settings observed for reported participants with the MacBookPro and for two observers with the Asus laptop. Power across stimulated frequencies is consistently above noise floor for all thresholds (from ~+5dB to +~14dB for MacBookPro volume setting 18 to 44%), and floor noise levels are consistent across volume settings. (As would be expected, the Asus laptop output line noise level is generally ~3-4 dB higher that of the MacBook Pro, with an additional ~2-3 dB noise levels in frequencies ~< 400Hz).

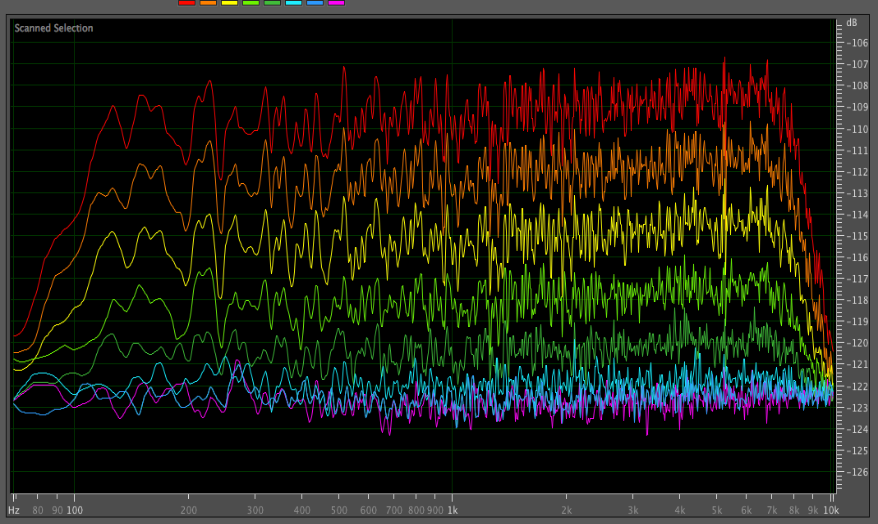

Figure S1a, MacBookPro: **white noise** stimulus power spectrum at volume settings 6% to 44%, with cool to hot colors showing increasing laptop volume settings. The lowest volume settings achieved by participants corresponded to the lowest green line. Magenta line shows sound card on, no stimulus playing.

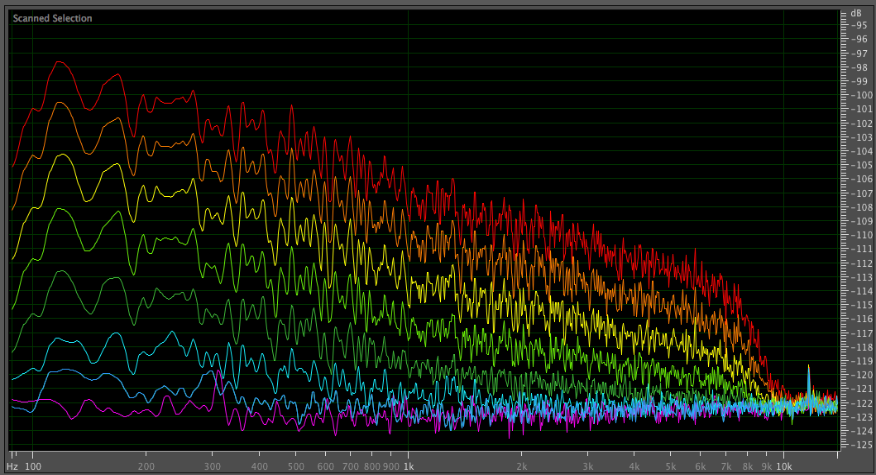

Figure S1b, MacBookPro: **pink noise** stimulus power spectrum at volume settings 6% to 44%, with cool to hot colors showing increasing laptop volume settings. The lowest volume settings achieved by participants corresponded to the lowest green line. Magenta line shows sound card on, no stimulus playing.

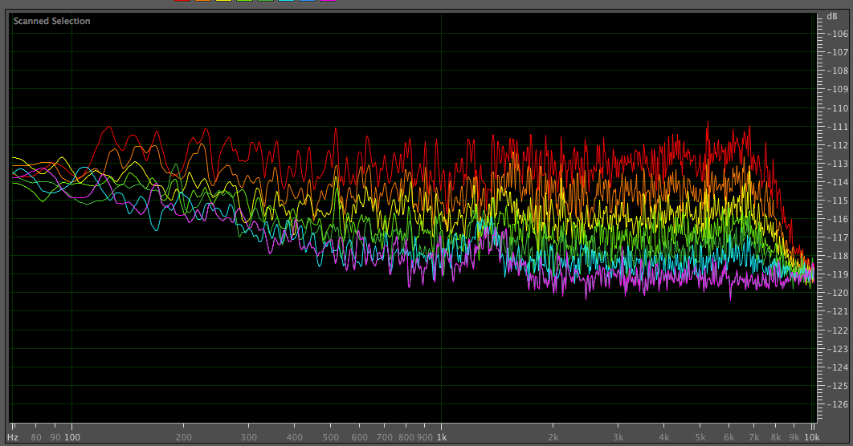

Figure S1c, Asus laptop: **white noise** stimulus power spectrum at sound card output levels typical for normal hearing listeners, with cool to hot colors showing increasing laptop volume settings. Magenta line shows sound card on, no stimulus playing.

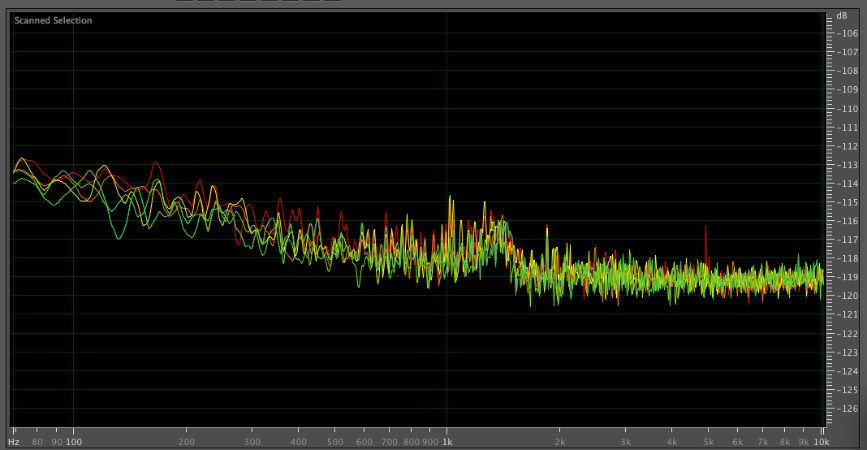

Figure S1d, Asus laptop: recording of silent period at same laptop volume settings as above (sound card still active).

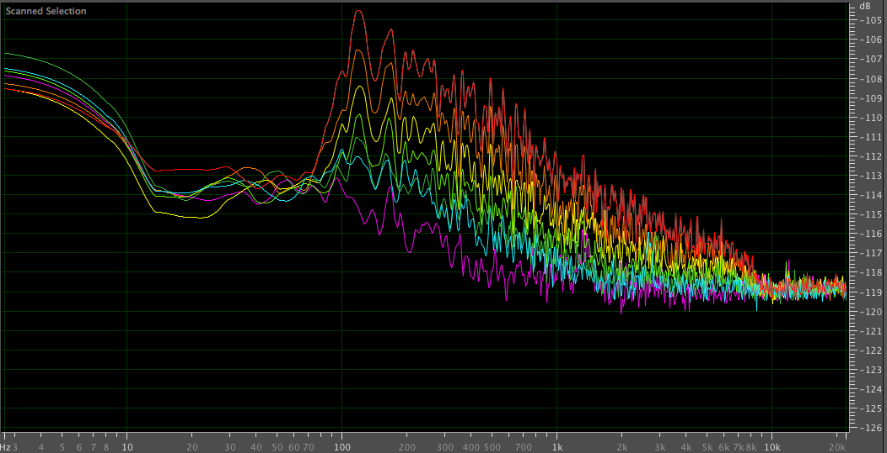

Figure S1e, Asus laptop: **pink noise** stimulus power spectrum at sound card output levels typical for normal hearing listeners, with cool to hot colors showing increasing laptop volume settings. Magenta line shows sound card on, no stimulus playing. (Note that x-axis differs from white noise so as to show more detail in lower frequencies).

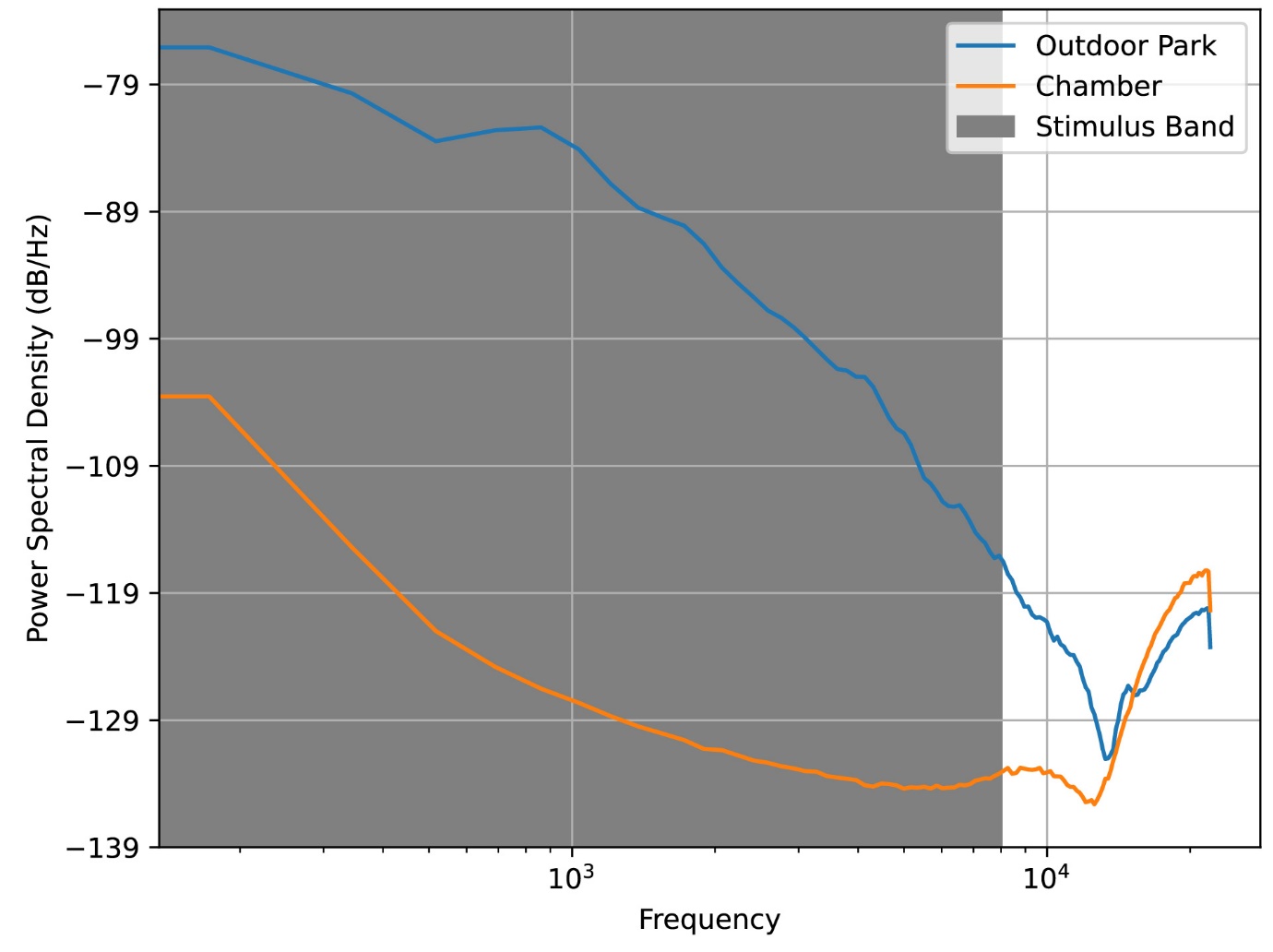

**Figure S2:** The power spectral densities of each acoustic environment used in Expt 1c, derived from 10-s recordings. The shaded area indicates the frequency band of the white noise stimulus used (80 – 8000 Hz).

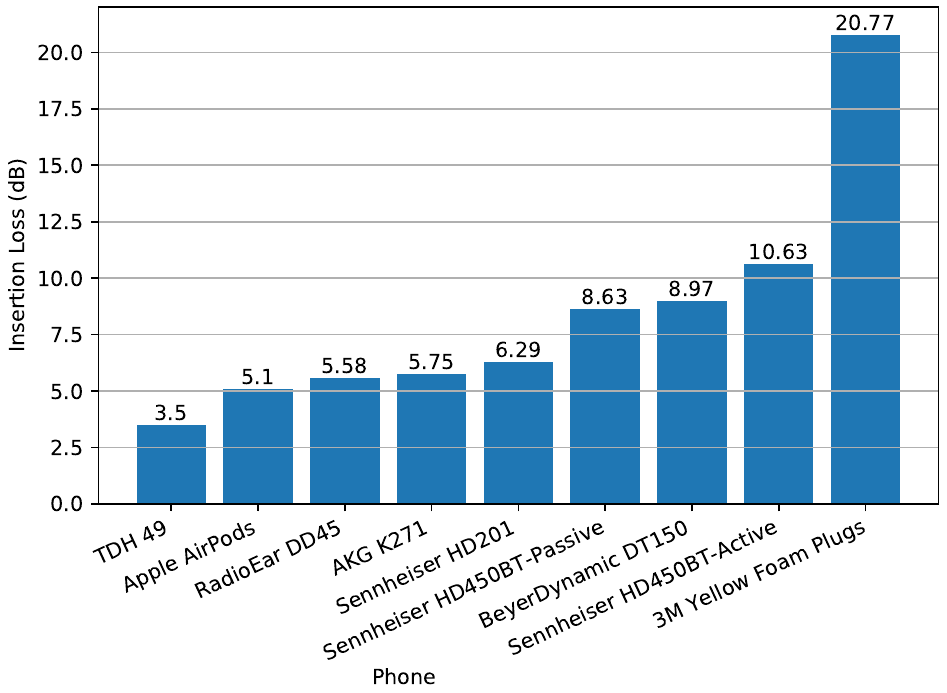

**Figure S3:** Insertion loss using various earbud and headphone models.

### Apathy Motivation Index (AMI) questionnaire

Think about your recent life. Which option best describes you over the last two weeks?

**Questions**

1. I feel sad or upset when I hear bad news

2. I start conversations with random people

3. I enjoy doing things with people I have just met

4. I suggest activities for me and my friends to do

5. I make decisions firmly and without hesitation

6. After making a decision, I will wonder if I have made the wrong choice

7. Based on the last two weeks, I would say I care deeply about how my loved ones think of me

8. I go out with friends on a weekly basis

9. When I decide to do something, I am able to make an effort easily

10. I don't like to laze around

11. I get things done when they need to be done, without requiring reminders from others

12. When I decide to do something, I am motivated to see it through to the end

13. I feel awful if I say something insensitive

14. I start conversations without being prompted

15. When I have something I need to do, I do it straightaway so it is out of the way

16. I feel bad when I hear an acquaintance has an accident or illness

17. I enjoy choosing what to do from a range of activities

18. If I realise I have been unpleasant to someone, I will feel terribly guilty afterwards

**Responses**

| **Response** | **Score** |
| --- | --- |
| Completely untrue | 4 |
| Mostly untrue | 3 |
| Neither true nor untrue | 2 |
| Quite true | 1 |
| Completely true | 0 |

**Calculation**

The level of apathy is assessed by taking the mean rating of the items within the subscale. A higher score indicates greater apathy.

| **Theme** | **Questions** | **Min** | **Max** |
| --- | --- | --- | --- |
| Behavioural activation | 5, 9, 10, 11, 12, 15 | 0 | 24 |
| Social motivation | 2, 3, 4, 8, 14, 17 | 0 | 24 |
| Emotional sensitivity | 1, 6, 7, 13, 16, 18 | 0 | 24 |

### Questions to track the dynamics of motivation and fatigue across the experiment

| **Rating** | **Question** | **Left Label** | **Right Label** |
| --- | --- | --- | --- |
| Motivation | How motivated do you feel? | Not motivated at all | Very motivated |
| Fatigue | How tired do you feel now? | Not tired | Very tired |
| Confidence | Before threshold: How well do you think you will do in the task? | Very bad | Very good |
|  | After threshold: How well do you think you did in the task? |  |  |
